# Keratinocytes Drive the Epithelial Hyperplasia Key to Sea Lice Resistance in Coho Salmon

**DOI:** 10.1101/2023.10.15.562030

**Authors:** S.J. Salisbury, R. Ruiz Daniels, S.J. Monaghan, J.E. Bron, P.R. Villamayor, O. Gervais, M.D. Fast, L. Sveen, R.D. Houston, N. Robinson, D. Robledo

**Affiliations:** The Roslin Institute and Royal (Dick) School of Veterinary Studies, University of Edinburgh, Edinburgh, UK; Institute of Aquaculture, University of Stirling, Stirling, UK; University of Santiago de Compostela, Santiago de Compostela, Spain; Atlantic Veterinary College, University of Prince Edward Island, Charlottetown, Canada; Nofima AS, Tromsø, Norway; Benchmark Genetics, 1 Pioneer Building, Edinburgh Technopole, Milton Bridge, Penicuik, UK; Sustainable Aquaculture Laboratory -Temperate and Tropical (SALTT), Deakin University, Victoria 3225, Australia

**Keywords:** snRNAseq, salmon, aquaculture, disease, parasite, sea lice, cell type, skin, immunity, wound-healing

## Abstract

**Background:** Salmonid species have followed markedly divergent evolutionary trajectories in their interactions with sea lice. While sea lice parasitism poses significant economic, environmental, and animal welfare challenges for Atlantic salmon (*Salmo salar*) aquaculture, coho salmon (*Oncorhynchus kisutch*) exhibit near-complete resistance to sea lice, achieved through a potent epithelial hyperplasia response leading to rapid louse detachment. The molecular mechanisms underlying these divergent responses to sea lice are unknown.

**Results:** We characterised the cellular and molecular responses of Atlantic salmon and coho salmon to sea lice using single-nuclei RNA sequencing. Juvenile fish were exposed to copepodid sea lice (*Lepeophtheirus salmonis*), and lice-attached pelvic fin and skin samples were collected 12h, 24h, 36h, 48h, and 60h after exposure, along with control samples. Comparative analysis of control and treatment samples revealed an immune and wound-healing response that was common to both species, but attenuated in Atlantic salmon, potentially reflecting greater sea louse immunomodulation. Our results revealed unique but complementary roles of three layers of keratinocytes in the epithelial hyperplasia response leading to rapid sea lice rejection in coho salmon. Our results suggest that basal keratinocytes direct the expansion and mobility of intermediate and, especially, superficial keratinocytes, which eventually encapsulate the parasite.

**Conclusion:** Our results highlight the key role of keratinocytes to coho salmon’s sea lice resistance, and the diverged biological response of the two salmonid host species when interacting with this parasite. This study has identified key pathways and candidate genes that could be manipulated using various biotechnological solutions to improve Atlantic salmon sea lice resistance.

## INTRODUCTION

Parasitism by sea lice is one of the greatest economic, environmental, and animal welfare issues facing the Atlantic salmon (*Salmo salar*, Linnaeus, 1758) aquaculture industry, with annual global costs exceeding £700 million [1]. Sea lice species, including the northern hemisphere’s *Lepeophtheirus salmonis* (Krøyer, 1837) and the southern hemisphere’s *Caligus rogercresseyi* [2], feed on salmon skin and fins, causing chronic open wounds in Atlantic salmon that can contribute to secondary infections [3]. Additionally, sea lice significantly reduce the market value of aquaculture fish – infestations have been estimated to cost US$0.46/kg of biomass [4] – and can also cause considerable impacts on wild salmonids [5]. A variety of treatment strategies have been developed to mitigate sea lice infestations in Atlantic salmon aquaculture, but these can be costly, ineffective, environmentally-damaging, and cause reduced animal welfare [6]. For example, sea lice have evolved increasing resistance to the costly and potentially environmentally damaging chemical parasiticides that have historically been commonly applied to salmon aquaculture pens [5,7]. Preventative methods, particularly those improving the innate resistance of Atlantic salmon to sea lice, are therefore considered a more effective route to address this problem [6].

Relatively high heritabilities for sea lice resistance in Atlantic salmon [e.g., 8-10] suggest that selective breeding should be effective, particularly when informed by genotype information via genomic selection [11, 12]. However, counts of sessile lice are the only measure of resistance that is currently used, and genetic variation in the immune response of Atlantic salmon has been difficult to assess. In addition, despite the identification of some significant QTL [*e.g*., 13-15], sea lice resistance has proven to be a polygenic trait [11]. Given the absence of loci of large effect to target, the relatively long generation time of Atlantic salmon (3-4 years), and that modern salmon breeding programs must include multiple additional traits in their breeding goal, selective breeding is unlikely to result in clear improvements to sea lice resistance in the short-term [6]. More rapid increases in genetic resistance to sea lice through gene editing or other biotechnological approaches may be informed by investigation of closely related salmonid species demonstrating greater resistance to sea lice [16].

Coho salmon (*Oncorhynchus kitsutch*, Walbaum, 1792) demonstrate an innate ability to kill and expel sea lice. Within 24 hours of louse attachment, coho salmon mount an acute epithelial hyperplasia response associated with a thickening of the skin, inflammation, cell proliferation, and an infiltration of immune cells [17–19]. This localized swelling can even encapsulate attached lice after 10 days post exposure [17, 19] and causes 90% of lice to drop off their coho salmon hosts between 7 – 14 days post exposure [18, 20]. In contrast, minimal swelling and rapid degradation of the epidermis occurs in response to an attached louse in highly susceptible Atlantic salmon [17]. The resistance of coho salmon to sea lice has therefore been proposed to be the result of an immune and wound-healing response that is greater in magnitude and very different in character compared to that of Atlantic salmon [21, 22]. This is supported by the upregulation of multiple genes associated with inflammation, tissue remodelling, and cell adhesion in the skin of coho salmon but not Atlantic salmon in response to sea lice [22, 23]. Both Atlantic salmon and coho salmon have also been suggested to mount a nutritional immune response to sea lice [24, 25], where iron availability is limited to deter iron-seeking pathogens [26]. However, the exact molecular and cellular mechanisms underlying coho salmon’s resistance to sea lice remain elusive.

This uncertainty is in part due to the cellular heterogeneity of fish skin. The skin’s multiple layers demonstrate distinct transcriptomic profiles reflecting each layer’s unique composition of cell types [27]. The outermost layer of skin, the epidermis, is populated primarily by filament-filled keratinocytes [28] in three layers: an upper layer of flattened superficial keratinocytes, an intermediate layer of amorphous keratinocytes, and a lower layer of cuboidal basal keratinocytes [29, 30]. Specialized mucous cells are found individually throughout the epithelium and play an important role in maintaining skin integrity through mucus production [30, 31]. The dermal layer below contains fibroblasts, blood vessels, and chromatophores [30, 31] as well as scales in the trunk and fin rays in the fins, both maintained by osteoblasts [30, 32, 33]. Both epidermal and dermal layers are punctuated by endothelial blood vessels and neural structures [34]. Muscle and fat lie below the dermis and are not considered part of the skin [30]. There is also a variety of resident immune cells in the skin including T cells, B cells, neutrophils, dendritic cells, and macrophages [35].

The large diversity of specialised cell types present within the skin therefore poses a problem for traditional bulk transcriptomic approaches which average gene expression across all cell types within a tissue and may therefore be unable to detect biologically relevant cell-type specific differential gene expression in highly heterogeneous tissues [36]. Single nuclei RNA sequencing (snRNAseq) offers a solution to this issue by generating individual transcriptomes for thousands of individual cells [37]. Cells can be grouped based on their individual transcriptomes into distinct cell type clusters, whose identities can be ascertained from diagnostic marker genes, uniquely expressed in each cluster. These technologies allow the study of biological processes with unparalleled resolution, facilitating the comparison of the same cell type across groups or species.

The aim of this work was therefore to use snRNAseq to investigate the cell types and gene expression patterns characterizing the response to sea lice in the skin of Atlantic salmon and coho salmon. We specifically targeted the first 60 hours post infection by *L. salmonis* copepodids. This time frame has been largely unexplored from a transcriptomic perspective despite being associated with significant histological changes leading to lice rejection in coho salmon [23]. Comparing the cell type-specific responses of resistant and susceptible species to sea lice allowed us to identify cell types and molecular pathways involved in determining the mechanisms of resistance in coho salmon and to pinpoint candidate genes that could be targeted to improve sea lice resistance in Atlantic salmon aquaculture.

## RESULTS

A total of 10 and 12 snRNAseq libraries passed filtration for Atlantic salmon and coho salmon, respectively. These had over 244 million reads each, and at least 73% and 86% of those reads aligned uniquely to the genome, for Atlantic salmon and coho salmon, respectively (Table S1,S2). The final total number of cells obtained for each species was 50328 for Atlantic and 48341 for coho salmon (Table S3).

### 1. Cell Type Identities and Marker Genes

A total of 23 cell clusters were observed within each species, after clustering cells independently by species (Fig.1a,b). These clusters demonstrated distinct transcriptomic profiles and their inferred identities were consistent across species (Fig.1). Marker genes were frequently identical for the same cell type across species (Fig.1 c,d, see Fig.S1,S2 for dot plots of additional cell markers, Table 1 for functional relevance of all marker genes for ascribed cell type identity, Table S4,S5 for counts per cell type and sample) and highly concordant between fin and skin tissue types (Fig.S3,S4). We identified all cell types expected in these tissues [30, 92] as well as several previously unreported cell types including a tuft-like “secretory” cell type.

**Fig. 1.**
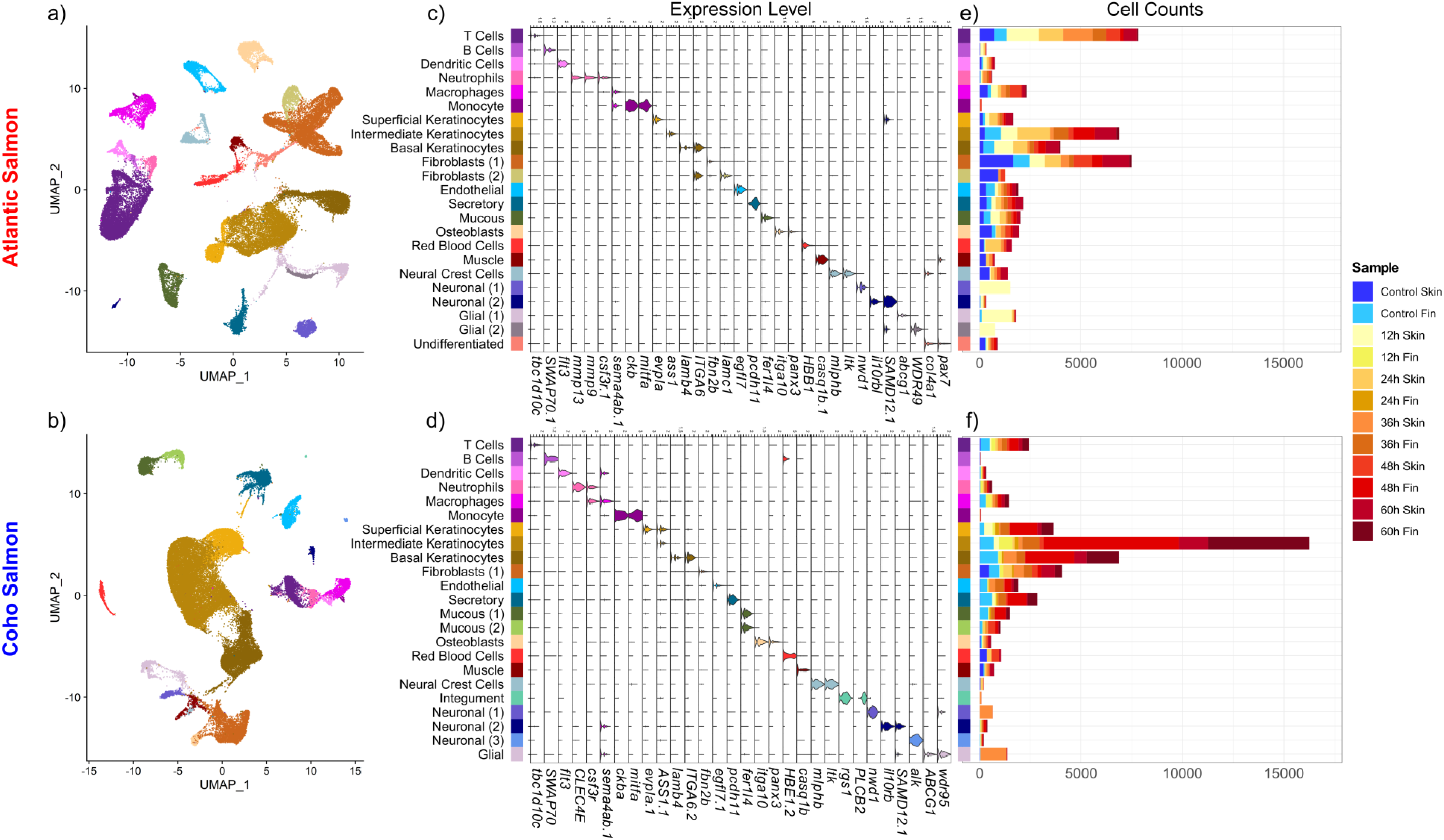
Cell types detected in Atlantic (a-c) and Coho (d-f) Salmon. UMAPs of cell clusters coloured by putative identity for (a) Atlantic salmon and (b) coho salmon. Violin Plots of marker genes for each cell cluster for (c) Atlantic salmon and (d) Coho Salmon. Counts of each cell type by sample for (e) Atlantic salmon and (f) coho salmon. Note there is no 12h Fin or 24h Fin sample for Atlantic salmon.

**Table 1.**
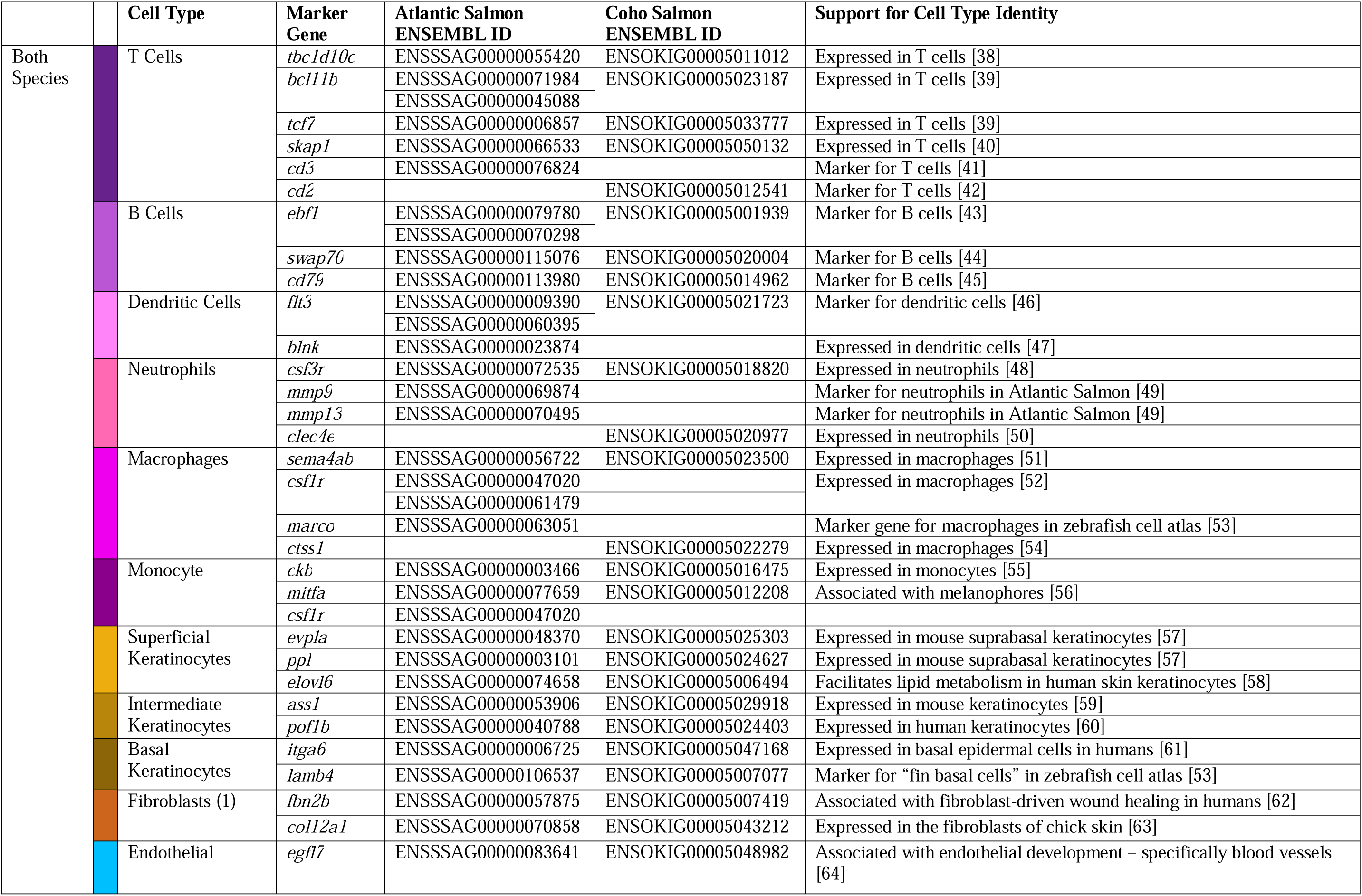
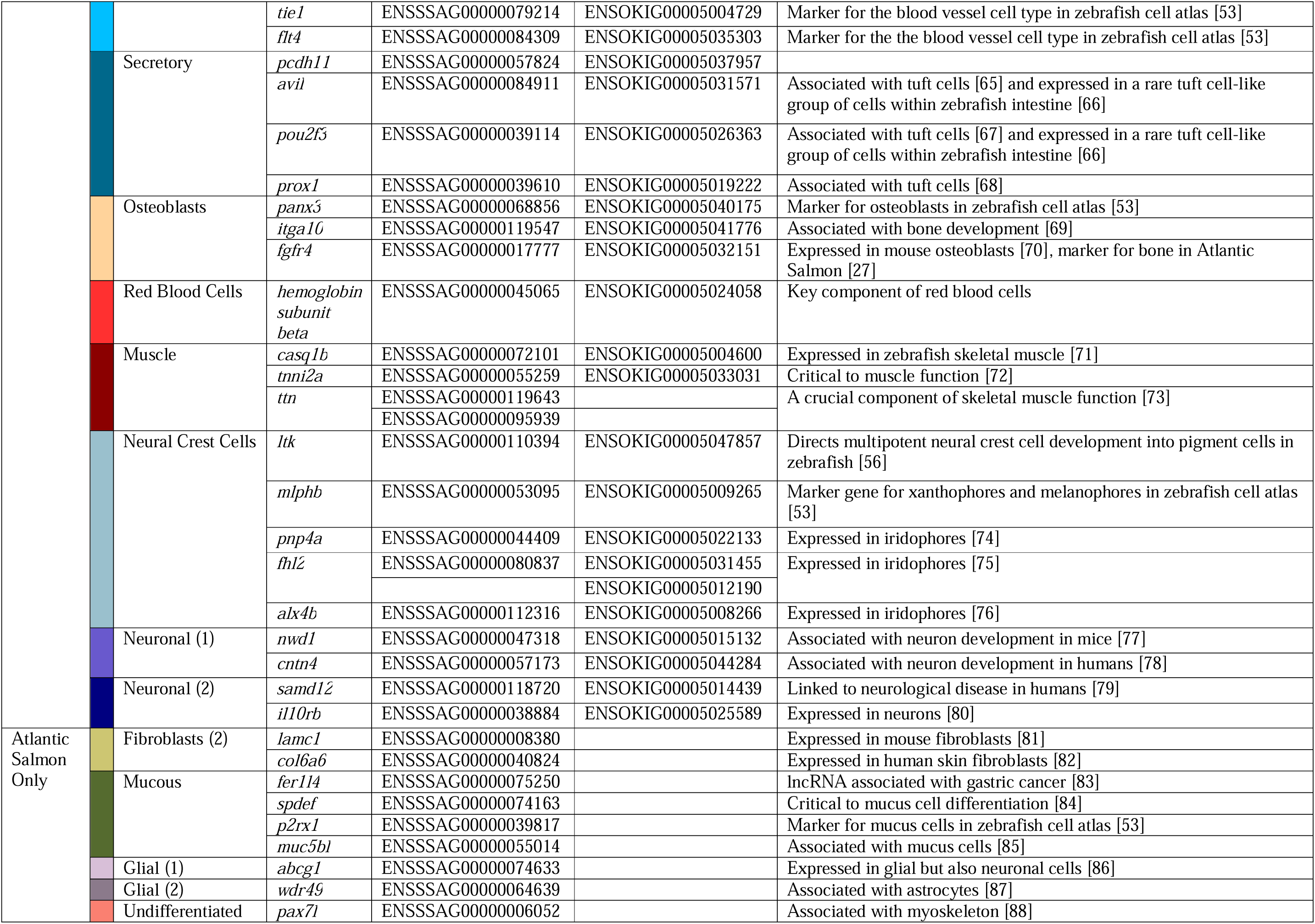
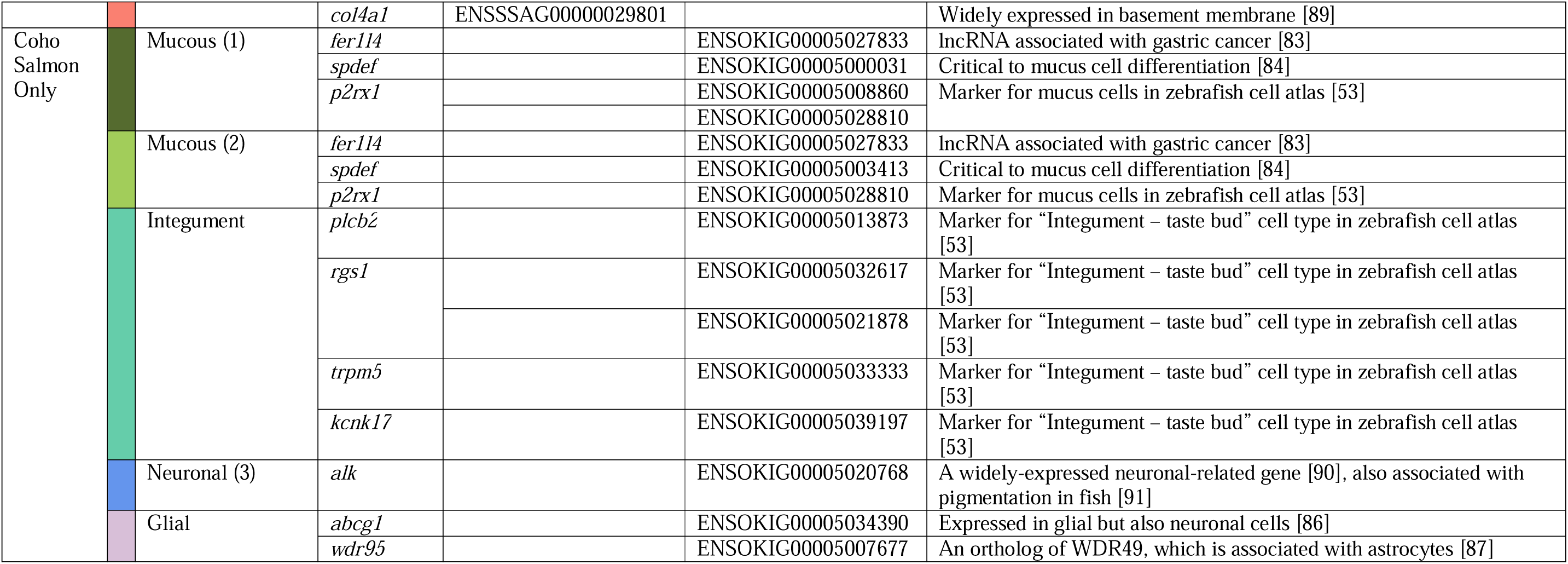
Marker genes for cell types found in skin and fin samples of Atlantic salmon and coho salmon. All noted genes were significantly (p << 0.001) upregulated in the given species’ cell type cluster relative to all other cells.

The integration of samples from both species demonstrated the majority of cell types observed in each of the species-specific datasets (Fig.2 a). Two clusters of immune cells were uncovered in the combined dataset which we designated “lymphocyte” and “myeloid” given their expression of *itgae* [93] and *cd163* [94], respectively. The marker genes for each cluster of the combined dataset were often identical to those marker genes in the corresponding cluster in the species-specific dataset and always highly expressed (Fig.2 b,c,d), confirming the presence of identical cell types in the skin of Atlantic salmon and coho salmon. However, the species-specific datasets presented additional clusters and had a greater number of marker genes given more genes were used in the clustering (salmonids present a recent whole-genome duplication and the establishment of 1:1 orthologies are not straightforward, which resulted in many genes being removed when the datasets of the two species were combined). Therefore, all further analyses were conducted using the species-specific datasets, which we refer to exclusively from this point forward.

**Fig. 2.**
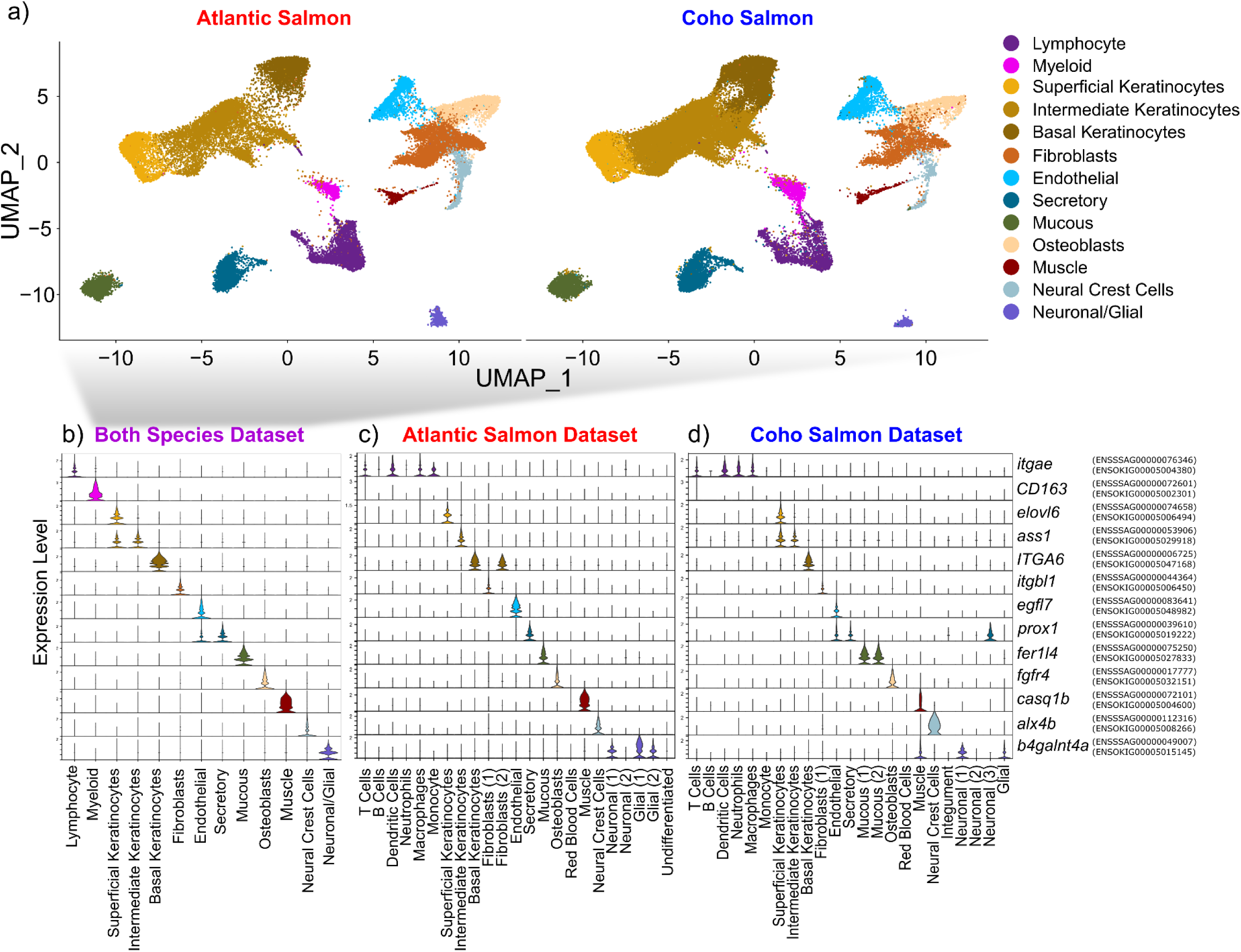
Cell clusters identified integrating both Atlantic salmon and coho salmon samples using 1:1 orthologous genes. a) UMAP of cell clusters split by species, b) violin plot of expression of a marker gene for each cluster. Violin plots visualize the expression of these same features in the species-specific datasets: c) Atlantic salmon, d) coho salmon. The Atlantic salmon and coho salmon ortholog ENSEMBL codes are noted to the right of each gene.

#### 1.1 Non-Immune Cell Types

Keratinocytes were among the most abundant cell types. Three keratinocyte clusters were identified: basal keratinocytes, superficial keratinocytes and a third cluster of “intermediate keratinocytes”, likely located between the former two keratinocyte layers and consistent with the three layers of keratinocytes observed in fish skin [29, 30]. Keratinocytes were abundant in all samples, but notably increased at 48h and 60h post infection only in coho salmon (Fig.1 e,f).

Other abundant cell types include fibroblasts, endothelial cells, and osteoblasts. Mucous cells were split into two clusters in coho salmon with many overlapping markers (Fig.S5,S6), but differing in their relative expression of different paralogs of *spdef* and *p2rx1* (see Fig.S2,S5,S6). Interestingly, *muc5* (associated with mucous cells, [85]) was expressed only in Atlantic salmon mucous cells (Fig.S1,S2). A “secretory” cell type was abundant in both species, and expressed tuft-cell marker genes (Table 1). Tuft cells line the epithelium of the gut and airway in mammals, and although their function is not well-characterized, they are associated with initiating immune responses (e.g., activating Th2 cells in response to helmith endoparasitism in mice) [95]. We speculate these may be a sacciform cell type, previously noted in coho salmon [20]. However, the noted absence of sacciform cells in Atlantic salmon [20], means that the location, morphology, and function of this newly identified cell type requires further investigation.

Neural crest cells were characterized by multiple pigment cell genes (Table 1) including *ltk*, which directs multipotent neural crest cell development into pigment cells in zebrafish [56], suggesting these cells are pigment cell progenitors. The detection of neural crest cells, red blood cells, and muscle cells predominately in trunk skin samples (Fig.1 e,f), is consistent with expectations of greater abundance of these cell types in the trunk skin than in the fins [30] given the potential to cut deeper into the dermal layer. Additionally, several clusters of neuronal and glial cells were observed, but most were observed in a single sample per species (Fig.1 e,f) suggesting they comprise neural structures which are present sporadically throughout the skin (e.g., peripheral axons [34], or the lateral line). Given their inconsistent presence within our samples we do not further consider the response of these cell types to sea lice, but note their potential to confound bulk RNAseq skin data.

Several cell types were identified in only one species. A small cluster of cells detected in coho salmon demonstrated a number of marker genes observed in cluster 196 “Integument-Taste Bud” of a zebrafish cell atlas [53] (Table 1), which we refer to as “integument” cells henceforth. We speculate this cell cluster may represent a rare chemosensory cell type in coho salmon, which may also be present in Atlantic salmon but was unobserved due to its rarity (N = 93 cells in coho salmon). Fibroblasts (2) were detected in Atlantic salmon but not coho salmon and expressed *lamc1* and *col6a6* but also marker genes of the keratinocyte clusters (e.g., *itga6* and *pof1b*) (Fig.S1). A final cell cluster unique to Atlantic salmon was termed “Undifferentiated” because of its few distinctive marker genes (Fig.S1,S7).

#### 1.2 Immune Cell Types

The immune cell marker gene *cd45* [96] was expressed in four and two clusters for Atlantic salmon and coho salmon, respectively (Fig.S8). These clusters were re-clustered to investigate for additional immune cell types expected to be present in the skin and potentially involved in sea lice response [18]. Sub-structuring within *cd45*+ cells revealed six main types of immune cells in both species: T cells, B cells, dendritic cells, neutrophils, macrophages, and monocytes (Fig.3 a,b). Myeloid and lymphocyte cells were clearly differentiated by the expression of *spi1b,* a marker for the myeloid lineage in zebrafish [39]. Marker genes for all immune cell types were consistent with the literature (Table 1) with the curious exception of the monocyte marker gene *mitfa*, typically associated with melanophores [56], suggesting these monocytes might develop into melanomacrophages known to be present in salmonid skin [19] (Fig.1 c,d, Fig.3 c,d, see Fig.S9-31 for violin plots of top marker genes).

**Fig. 3.**
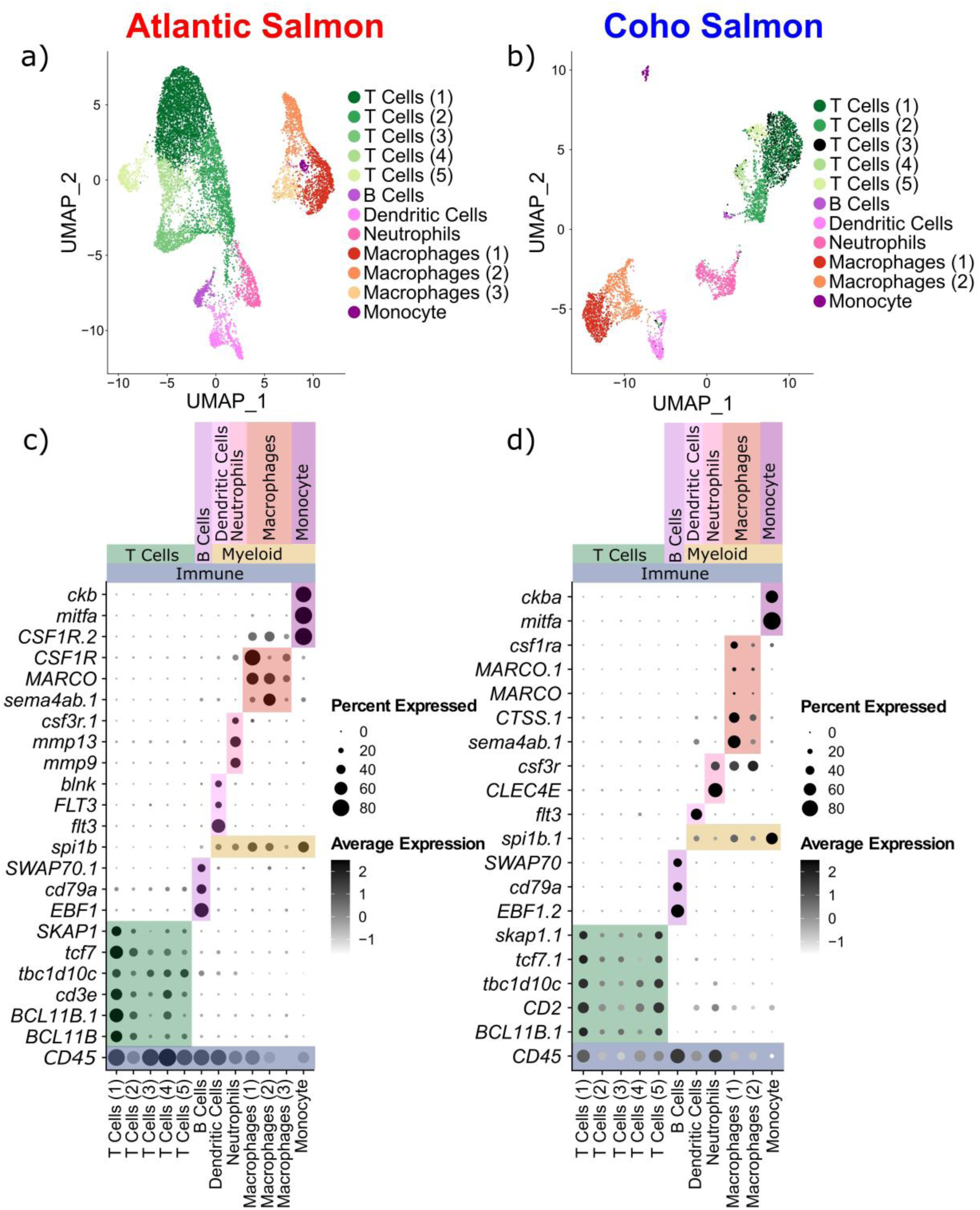
Sub-clustering of putative immune cells expressing CD45. UMAP visualization of immune clusters in Atlantic (a) and Coho (b) Salmon. Dot Plots of features characterizing immune cell types in Atlantic (c) and Coho (d) Salmon.

While multiple macrophage and T cell subclusters were apparent in each species, their top marker genes were either largely overlapping among subclusters, mostly ribosomal genes, or had unknown biological relevance (Fig.S9-12,S17-19,S21-23,S25,S29-30), suggesting these are clustering artefacts or previously undescribed immune cell types. For instance, expression of *cd4* and *cd8* also did not conclusively differentiate T cell subclusters (Fig.S32), however, T cells (5) in Atlantic salmon (Fig.S13) and T cells (4) in coho salmon (Fig.S24) expressed *gata3*, associated with Th2 cell activation [97]. Given this general lack of clear, biologically relevant expression differences within T cell and macrophage subclusters, and to maximize power for subsequent differential expression analyses (given the low numbers of cells in each T cell and macrophage subcluster, Table S6,S7) we grouped together all T cell subclusters and all macrophage subclusters for downstream analysis.

### 2. Common Responses to Sea Lice in Resistant and Susceptible Salmonid Species

A total of 4567 and 1799 unique genes were found to be differentially expressed between any treatment time point and the control in Atlantic salmon and coho salmon, respectively (see Fig.S33-35 for the distribution of differentially expressed genes within a given cell type, see Fig.S36,S37 for GO enrichment results). Some conserved wound-healing and immune responses to sea lice infection were detected in Atlantic salmon and coho salmon.

#### 2.1 Wound-Healing Response to Sea Lice

Both species showed a clear activation of wound-healing mechanisms in response to the parasite in a variety of cell types (Fig.4). Upregulation of genes linked to limb development such as *pax9* [98] and *meis2* [99] were evident in keratinocytes, mucous cells, and/or fibroblasts. Genes associated with extracellular matrix integrity including *pdgfra* [100] and *col21a* [101] were upregulated in fibroblasts of both species. Another gene associated with healing of individual cells, *abr* [102], was significantly upregulated in macrophages and T cells in coho salmon and in mucous cells, keratinocytes, and T cells in Atlantic salmon. The upregulation of *agr2* observed in mucous cells of both species probably reflects an increased production of mucus in response to sea lice [103] potentially to aid in wound-healing [30, 92]. A gene previously found to be upregulated at louse attachment sites in Atlantic salmon [104], *aloxe3*, was upregulated in mucous cells of both species but only significantly in Atlantic salmon. Mutations to *aloxe3* are associated with ichthyosis, a condition resulting in the build-up of skin cells [105], suggesting this gene could contribute to wound-healing-associated cell growth. Similarly, epidermal reinforcement-related genes *cldn8* [106] and *cntn1* [107] were more upregulated in Atlantic salmon. However, *bnc2*, associated with wound-healing and fibrosis [108], as well as black pigmentation [109], was upregulated earlier and more strongly in coho salmon basal keratinocytes. Similarly, *hspe2,* associated with cell proliferation and extracellular matrix strengthening [110], was upregulated in coho salmon fibroblasts but downregulated in Atlantic salmon fibroblasts. Therefore, while general wound-healing mechanisms are activated in both species, differences can be detected.

**Fig. 4.**
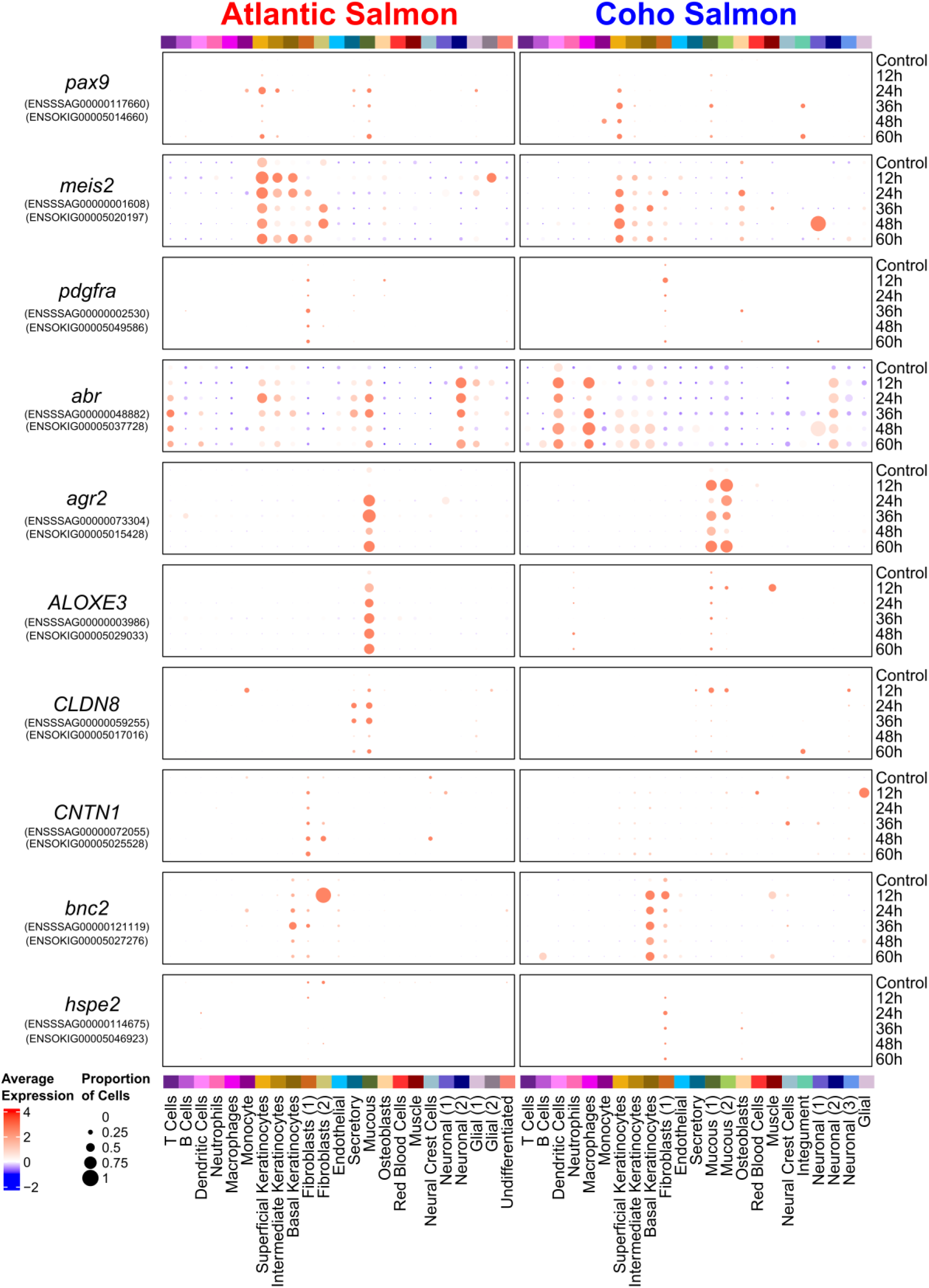
Dotplots of wound healing-related gene expression in Atlantic salmon and/or coho salmon in response to sea lice. All genes shown were significantly differentially expressed (p_adj_ < 0.001) in at least one pairwise comparison between the control and any treatment timepoint in either species.

#### 2.2 Immune Response to Sea Lice

A clear immune response was observed in both species in response to sea lice (Fig.5). Multiple paralogs of genes associated with immune cell development including *runx3* [111], *rarab* [112], and *gnai2* [113] were upregulated in response to sea lice in a variety of immune cell types including T cells, macrophages, and dendritic cells (Fig.5a). *Myo9b*, a gene associated with immune cell motility and activation [114] was upregulated in dendritic cells, neutrophils, and macrophages in both species, though showing a faster and more intense upregulation in coho salmon (Fig.5a). Major histocompatibility components were significantly upregulated in macrophages and T cells (*MHCII* only) but surprisingly in non-immune cell types too, mainly keratinocytes, and particularly superficial keratinocytes (Fig.5b). The involvement of the complement immune system was unclear. Two paralogs of *c4* were upregulated in Atlantic salmon fibroblasts while in coho salmon fibroblasts, one paralog was not differentially expressed, and the other was upregulated at 24h but downregulated at 36h and 60h (Fig.5c). C*fd* was significantly downregulated in Atlantic salmon fibroblasts but was not significantly differentially expressed in coho salmon (Fig.5c). This is consistent with previous observations of the downregulation of this gene in Atlantic salmon in response to *L. salmonis* sea lice [104]. Though Atlantic salmon demonstrated robust activation of T cells through the significant upregulation of *cd28*, *ifit9*, *sox4* [115], *cxcr4* [116], and *ly-9* [117], they also significantly upregulated anti-inflammatory *socs3* [118] (Fig.5d).

**Fig. 5.**
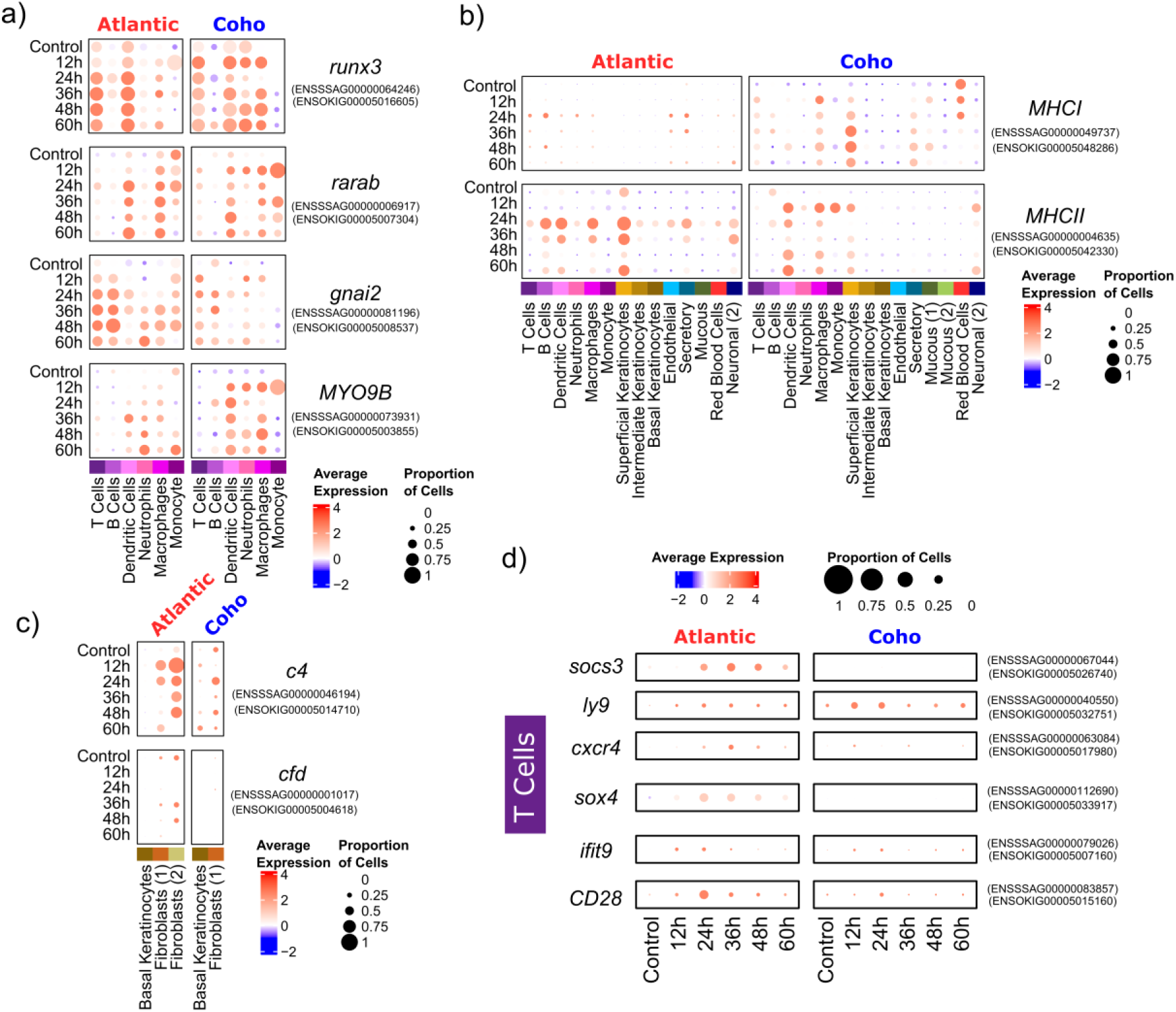
Dotplots of immune-related gene expression in Atlantic salmon and/or coho salmon in response to sea lice. a) immune genes upregulated in both species, b) MHC genes upregulated in both species, c) complement immune system gene expression, d) immune-related genes particularly upregulated in Atlantic salmon in response to sea lice. All genes shown were significantly differentially expressed (p_adj_ < 0.001) in at least one pairwise comparison between the control and any treatment timepoint in either species.

**Fig. 6.**
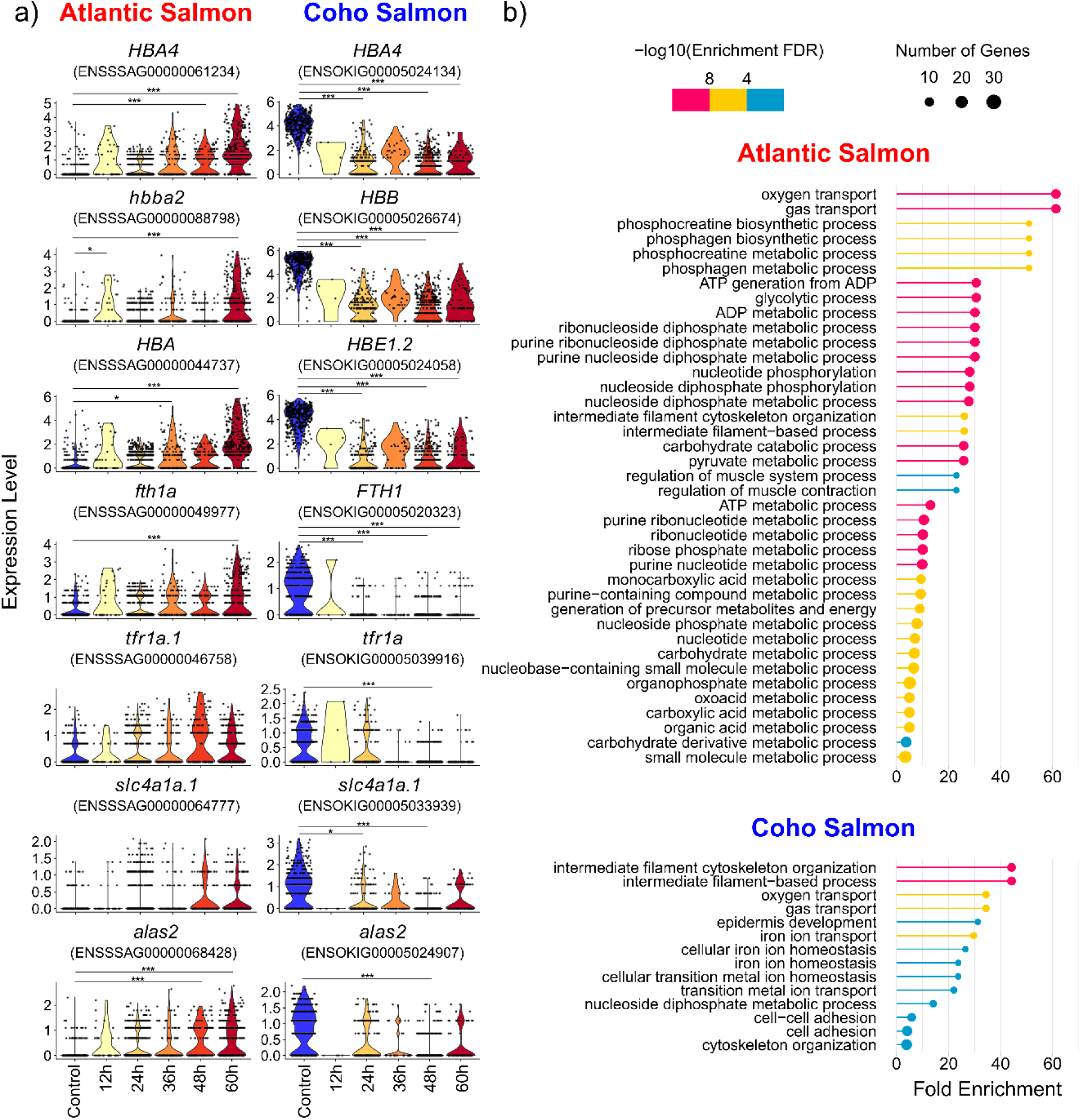
Red blood cell response to sea lice in Atlantic salmon and coho salmon. a) violin plots of gene expression in Atlantic salmon and coho salmon of genes significantly upregulated in coho salmon keratinocytes (p_adj_ < 0.001) in response to sea lice in at least one treatment timepoint relative to the control(* - p_adj_ < 0.001, ** - p_adj_ < 0.0001, *** - p_adj_ < 0.00001), b) significantly enriched biological GO terms (padj < 0.001) for red blood cells in response to sea lice in Atlantic salmon and coho salmon.

**Fig. 7.**
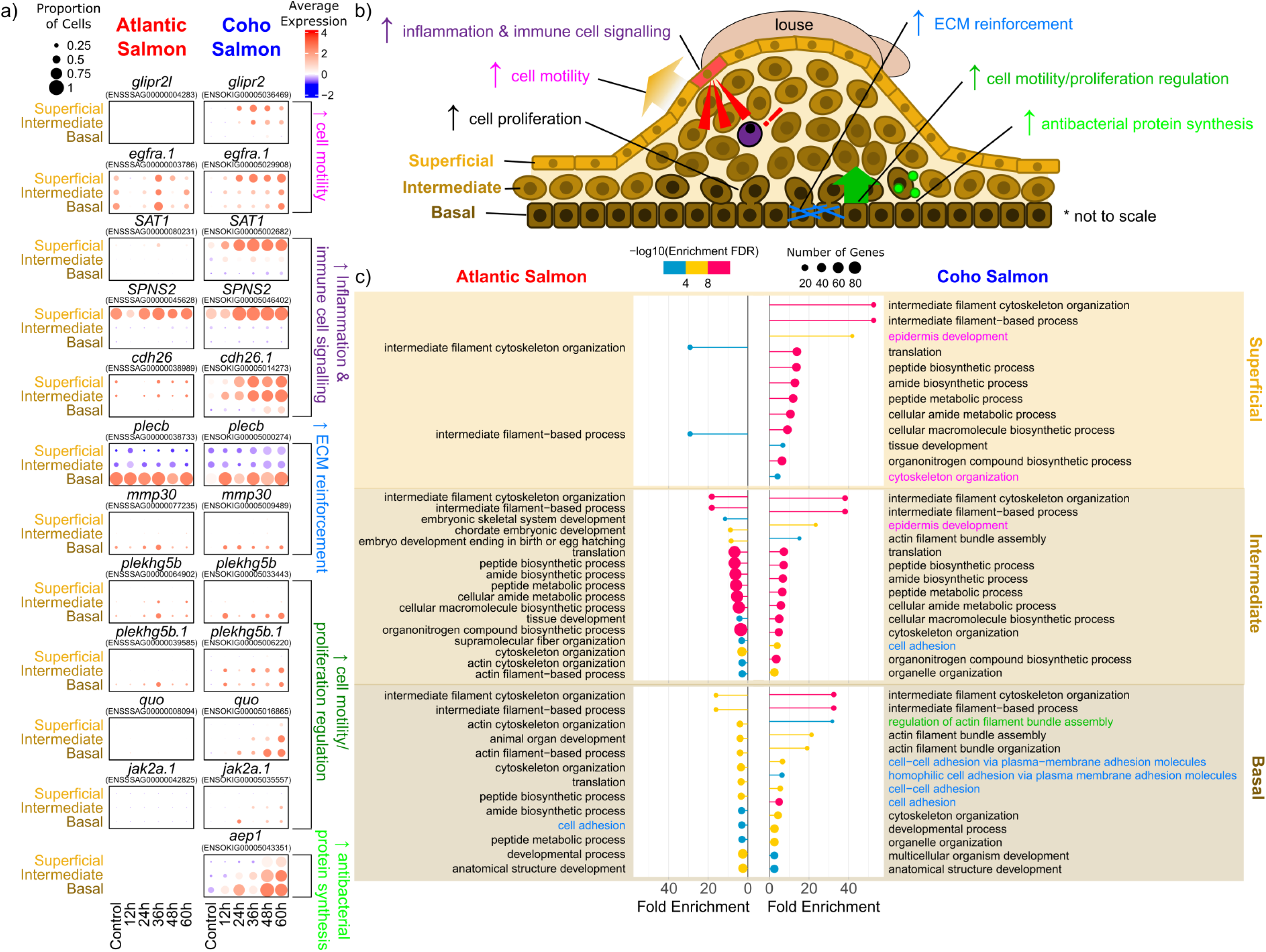
Keratinocyte response to sea lice underlies coho salmon resistance to sea lice. a) dotplots of gene expression in Atlantic salmon and coho salmon of genes significantly upregulated in coho salmon keratinocytes (p_adj_ < 0.001) in response to sea lice in at least one treatment timepoint relative to the control, b) proposed unique contributions of superficial, intermediate, and basal keratinocytes to epithelial hyperplasia immune response to sea lice in coho salmon, c) significantly enriched biological GO terms (padj < 0.001) for superficial, intermediate, and basal keratinocytes in response to sea lice in Atlantic salmon and coho salmon. Differentially expressed genes in a) and GO terms in c) are colour-coded by the biological processes depicted in b) that they are potentially associated with.

### 3. Responses to Sea Lice Unique to Coho Salmon

#### 3.1 Downregulation in Coho Salmon Red Blood Cells in Response to Sea Lice

Atlantic salmon red blood cells upregulated a number of genes associated with iron binding including several hemoglobin and ferritin subunits, and *tfr1a* [119] and other genes key to red blood cell function including s*lc4a1a* (ion transportation, [120], and *alas2* (heme biosynthesis, [25]) (Fig.6a). On the contrary, there was a significant downregulation of these genes in coho salmon red blood cells (Fig.6a). A regulation of iron in coho salmon red blood cells was further supported by the enrichment of a variety of iron-related GO terms (e.g., iron ion transport – GO:0006826) in sea louse infected samples of coho salmon but not Atlantic salmon (Fig.6b).

#### 3.2 Keratinocytes are Key to Epithelial Hyperplasia Response to Sea Lice in Coho Salmon

Coho salmon keratinocytes exclusively significantly upregulated a variety of genes associated with epidermal re-organization (Fig.7a,b). Keratinocytes in both species were enriched for intermediate filament cytoskeleton organization (GO:0045104) and intermediate filament-based process (GO:0045103), consistent with the known abundance of filaments observed in salmon keratinocytes [28] (Fig.7c). However, the fold enrichment was much higher in coho salmon, indicating greater cell movement and restructuring of keratinocytes in this species (Fig.7c).

Coho salmon superficial keratinocytes expressed genes more associated with cell motility and immune cell localization, consistent with their location in the outermost layer of the epidermis and in direct contact with attached lice [30] (Fig.7b). The GO term epidermis development (GO:0008544) was enriched in coho salmon superficial keratinocytes and to a lesser extent in intermediate keratinocytes (Fig.7c). Increased cell motility in coho salmon superficial keratinocytes and intermediate keratinocytes was also evident by the increased expression of *glipr2*, associated with cell migration particularly in response to hypoxia [121], and *egfra*, associated with epidermal cell proliferation [122] (Fig.7a). Coho salmon superficial keratinocytes also upregulated genes related to inflammation and immune cell infiltration including: *sat1* [123], *spns2* [124], and *cdh26* [125], (Fig.7a).

In contrast, the basal keratinocyte response in coho salmon was characterized by the upregulation of genes associated with extracellular matrix reinforcement, consistent with their location in the outermost layer of the dermis [30] (Fig.7b). Genes associated with cell adhesion and the extracellular matrix including *plecb* [126] and *mmp30* [127] were significantly upregulated in coho salmon (Fig.7a). GO terms associated with extracellular matrix development (e.g., cell-cell adhesion via plasma-membrane adhesion molecules – GO:0098742, and cell adhesion – GO: 0007155, which is also enriched in intermediate keratinocytes) were also significantly enriched in coho salmon basal keratinocytes (Fig.7c). This layer of keratinocytes may also be responsible for directing the movement of upper layers of keratinocytes through the upregulation of genes known to regulate cell motility including *plekhgb5b* [128] and *quo* [129] (Fig.7a) and supported by the significant enrichment for GO: 0032231, regulation of actin filament bundle assembly (Fig.7b,c). Coho salmon basal keratinocytes also upregulated the immune gene *jak2a* (Fig.7a), which regulates hematopoiesis [130], promotes cell proliferation [131] and is inhibited by *socs3* [132] (upregulated only in Atlantic salmon (Fig.5d)). An aerolysin-like protein, which breaks down cell membranes [133] and is upregulated in fish in response to bacterial infections [e.g., 134, 135], was also significantly upregulated exclusively in coho salmon basal keratinocytes (Fig.7a), confirming earlier observations of the upregulation of this gene exclusively in the skin of coho salmon but not of Atlantic salmon in response to sea lice [23].

The differentially expressed genes characterizing the intermediate keratinocytes response to sea lice in coho salmon largely overlapped with either the basal or superficial keratinocytes (Fig.7a). This less specialized role is consistent with their location between the superficial and basal keratinocytes. It may also reflect their recent generation from basal keratinocytes [136] as evidenced by the particular increase in abundance of this layer of keratinocytes at 48-60h (Fig.1f).

#### 3.3 Other Cell Types Potentially Contributing to Coho Salmon Epithelial Hyperplasia in Response to Sea Lice

Several additional cell types express genes related to inflammation in coho salmon (Fig.8). Secretory cells significantly upregulate *ttc7a* from 24h onward in coho salmon but this gene was only significantly upregulated at 36h in Atlantic salmon. This gene is associated with epithelial inflammation in mice [137]. Alternatively, *mrc1*, a gene linked to inflammation [138] and associated with increased *C. rogercresseyi* sea lice count on Atlantic salmon [139], was significantly upregulated in coho salmon but not Atlantic salmon endothelial cells. Coho salmon macrophages also demonstrated upregulation of the inflammation-associated gene *usp47* [140]. Multiple cell types may therefore potentially regulate the keratinocyte epithelial hyperplasia response to sea lice observed in coho salmon.

**Fig. 8.**
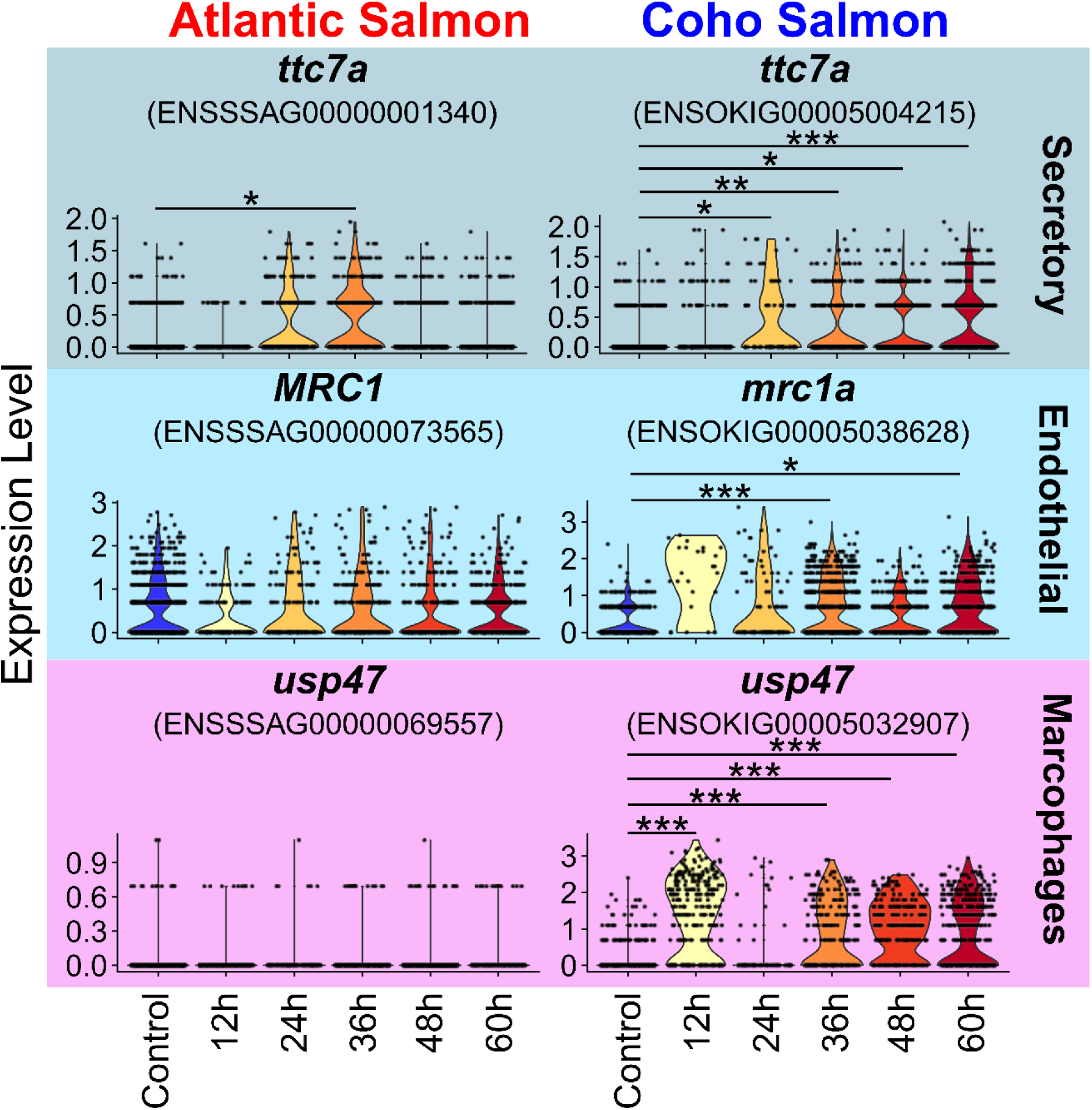
Violin plots of gene expression in Atlantic salmon and coho salmon in response to sea lice that are potentially regulating coho salmon’s epithelial hyperplasia response to sea lice. The cell type for which the expression of each gene is shown is noted to the right of each plot. (* - p_adj_ < 0.001, ** - p_adj_ < 0.0001, *** - p_adj_ < 0.00001.)

## DISCUSSION

Our results suggest that Atlantic salmon and coho salmon skin share a common set of cell types consistent with their recent divergence 30 million years ago [141]. Many of these cell types demonstrate a clear response to sea lice, which includes the activation of wound-healing and immune mechanisms, often common to both species. Conversely, lice immunomodulation of a variety of cell types was evident only in Atlantic salmon. Additionally, the coho salmon response to sea lice presented unique signatures, characterized by iron-limitation in red blood cells and a dramatic stimulation and re-organization of keratinocytes. These processes are likely to be major contributors to the greater resistance of this species to sea lice, and the underlying genes and regulatory networks detected here are potential candidates whose expression and functioning could be disrupted to “rewire” the host response to sea lice in Atlantic salmon via biotechnological approaches such as gene editing [16].

### Wound-Healing Response

Both species appear to employ a common wound-healing response to sea lice using a combination of keratinocytes, fibroblasts, mucous cells, and immune cells, in agreement with the critical role of these cell types in response to skin laceration [92]. The expression of limb development-related genes in multiple cell types also confirms a large-scale rearrangement of the skin in response to wounding [30]. Fibroblastic repair of the dermis, as expected shortly after wounding [30], was also evident through the upregulation of genes related with extracellular matrix reconstruction in fibroblasts in both species. Mucous cell upregulation of *abr2* also suggests both species increased mucus production in response to sea lice. Though sea lice feed on mucus [142], increased mucus production is a characteristic wound-healing response in Atlantic salmon [30, 92]. Alternatively, mucus upregulation may be particularly adaptive in coho salmon since, unlike Atlantic salmon, mucus of this species does not prompt a protease increase from sea lice, suggesting coho salmon mucus may contain protective qualities [143].

### Immune Response

Both species mount a common immune response to sea lice invoking the innate, adaptive, and complement immune systems. The upregulation of major histocompatibility proteins in the skin of both species is consistent with previous observations [24, 144]. However, the expression of *MHCII* in the superficial keratinocytes was surprising given that these cells are not typically associated with antigen presentation. Nonetheless, this result is consistent with and may explain previous observations of *MHCII* expression in Atlantic salmon epidermis in response to sea lice [24, 145]. Our results support the potential importance of superficial keratinocytes for sensing pathogens via antigen presentation and initiating immune and inflammatory responses [146].

Similarly, keratinocytes and fibroblasts seem to be key to the activation of the complement immune system. However, our results do not provide clear support for the importance of the complement immune response to sea lice resistance. This is consistent with previous observations of both the upregulation [139, 147] and downregulation [104] of complement proteins in Atlantic salmon in response to sea lice. Our results therefore support earlier suggestions that activation of the complement pathway may not be sufficient to grant sea lice immunity in Atlantic salmon [139].

Our results also potentially indicate that Atlantic salmon and coho salmon preferentially employ different immune cells in response to sea lice. Atlantic salmon had far more T cells than coho salmon (Fig.1e,f) perhaps as a consequence of artificial selection in this aquaculture strain of Atlantic salmon for greater disease resistance [148]. Atlantic salmon also demonstrated greater upregulation of genes associated with T cell activation. This observation may be partly attributable to differences in power among species to detect differential expression in T cells but is consistent with previous evidence suggesting a T cell dominated response to sea lice in Atlantic salmon [149]. In contrast, coho salmon potentially show a greater use of their macrophages in response to sea lice, as evidenced by the significant enrichment for “antigen processing and presentation” (GO:0019882) in coho salmon but not Atlantic salmon macrophages. Our results also support the key role of macrophages in directing coho salmon skin inflammation in response to sea lice [17], specifically through the upregulation of *usp47* and *ndst1a*, genes which are both associated with macrophage-driven inflammation [140, 150]. We speculate that coho salmon employ a macrophage-dominant innate immune response to sea lice, while Atlantic salmon try (and fail) to employ a T cell-led adaptive immune response. More sampling or targeted snRNAsequencing of immune cells, allowing for greater power to detect cell type heterogeneity within macrophages and T cells in each species, could be helpful to test this hypothesis.

A surprising result was the seeming lack of response in neutrophils to sea lice in either species. Few differentially expressed genes were observed in this cell type and no GO terms were enriched for either species, likely a result of low power due to the few neutrophils detected in each species. This scarcity of neutrophils was itself somewhat surprising given that previous histological work has suggested increased abundance of neutrophils at the site of wound healing in both species [17]. Upregulation of genes identified in this study as markers for neutrophils (e.g., *mmp9*, *mmp13*, *csf3r*) have also been observed to be upregulated at the site of sea lice attachment in both species [e.g., 22, 151]. This discrepancy may reflect a true relative rarity of neutrophils in comparison to other skin cell types (e.g., keratinocytes and fibroblasts which dominated our samples). Alternatively, this may be a sampling bias due to the demonstrated difficulty in capturing this cell type with scRNAseq [152]. More sampling, adjustment of nuclei isolation protocols to target immune cells, or integration of snRNAsequencing data with spatial transcriptomic data may therefore help us to learn more about what immune cells are doing in response to sea lice.

### Potential Immunomodulation of Atlantic Salmon By Sea Lice

Given the known susceptibility of Atlantic salmon to sea louse immunomodulation [153, 154], differences in immune and wound-healing response between Atlantic salmon and coho salmon may not only reflect host physiological differences but also the differential capacity of sea lice to immunomodulate each species. For example, upregulation of the inflammation-dampening *socs3* [118] in Atlantic salmon may be induced by sea louse immunomodulation. This gene is also upregulated in Atlantic salmon skin and head kidney in response to *C. rogercresseyi*, but is downregulated when Atlantic salmon are fed an immunostimulatory diet associated with lower lice counts, suggesting that this upregulation in response to *C. rogercresseyi* is maladaptive [155]. *Socs* genes are commonly targeted by fish pathogens to dampen host immunity [156] and may be particularly effective at preventing macrophage activation (e.g., in turbot in response to bacterial pathogens [157]). Our results therefore suggest that *L. salmonis* induce *socs3* upregulation in Atlantic salmon in order to weaken their hosts.

Lice immunomodulation may have also caused the dampened expression of *hspe2* and *bnc2* in Atlantic salmon, potentially resulting in reduced capacity for wound-healing, and, in the case of *bnc2*, melanism [109]. Melanism is frequently observed at the louse attachment sites in Atlantic salmon [30] and is more pronounced in Atlantic salmon with more sea lice resistance [158]. Therefore, sea lice may downregulate *bnc2* in Atlantic salmon to prevent effective wound healing.

Upregulation of haemoglobin and ferritin in Atlantic salmon red blood cells could also reflect lice immunomodulation for the purposes of increasing the parasite’s access to the host’s iron. Many pathogens manipulate iron homeostasis to increase available iron both for nutritional purposes and potentially as a method of weakening their host [159, 160], as excess iron can contribute to Fenton Chemistry production of harmful reactive oxygen-containing species [161]. Ferritin and genes related to heme biosynthesis have previously been observed to be upregulated in the skin of Atlantic salmon in response to *L. salmonis* [162]. This was suggested to be an adaptive compensatory response to blood loss from *L. salmonis* parasitism, however, we suggest that this may instead be a maladaptive response due to *L. salmonis* immunomodulation of Atlantic salmon. This is supported by the observation that haemoglobin is downregulated in Atlantic salmon infected with *C. rogercresseyi* when they are fed an immunostimulatory diet [155]. *L. salmonis* secretion of prostaglandin E2 or other vasodilators may underlie this response in Atlantic salmon [163]. Our results therefore suggest the potential for sea lice to manipulate a wide-range of molecular pathways and phenotypes in Atlantic salmon related to immune response, wound healing and iron availability. Additional molecular research from the perspective of the sea louse would be useful to substantiate these findings and elucidate the precise molecular strategies employed by the sea louse to elicit these responses in Atlantic salmon.

### Potential Nutritional Immune Response in Coho Salmon Red Blood Cells May Discourage Sea Lice

In contrast to Atlantic salmon, coho salmon red blood cells downregulate multiple iron-binding genes in response to sea lice. This could reflect differential wound-healing strategies in each species or may potentially indicate an adaptive nutritional immune response. Nutritional immunity, where hosts reduce the availability of iron in their tissues, is commonly employed to dissuade iron-seeking pathogens [26]. Pink salmon downregulate iron-associated genes in response to sea lice [164] and a nutritional immune response resulting from the upregulation of *hepcidin 1* has been suggested for both Atlantic salmon and coho salmon [24]. However, we found low expression of hepcidin in both species in all samples. Instead, our results suggest that this nutritional immune response in coho salmon is derived from the downregulation of a variety of iron-binding genes in red blood cells.

Yet, Atlantic salmon are clearly capable of mounting a similar nutritional immune response to other pathogens. For example, plasma iron significantly decreased in Atlantic salmon exposed to live and dead *Piscirickettsia salmonis* bacteria [119]. Intriguingly, Atlantic salmon seem capable of mounting a similar nutritional immune response by upregulating genes associated with heme degradation when parasitized by *C. rogercresseyi* but not *L. salmonis* [162]. *L. salmonis*’ longer co-evolutionary history with Atlantic salmon [165] may have resulted in its greater capacity to immunomodulate Atlantic salmon in comparison to *C. rogercresseyi.* Given the susceptibility of Atlantic salmon to both sea louse species, restoring Atlantic salmon’s adaptive nutritional immunity may not be sufficient to confer resistance to *L. salmonis*. However, this may still result in positive animal welfare consequences given that iron limitation can prevent opportunistic microbial infections [166] that are often associated with the sites of sea lice attachment [167].

### Keratinocytes Key to Coho Salmon Epithelial Hyperplasia Immune Response to Sea Lice

Our results strongly suggest that keratinocytes are responsible for the epithelial hyperplasia response characterized by filament development, inflammation, and cell proliferation that coho salmon employ to expel sea lice [17, 18, 21]. This is evidenced by our observations of a significant upregulation of genes associated with cell proliferation, cell motility, and extracellular matrix strengthening in keratinocytes, in addition to their dramatic increase in abundance during sea lice infection. However, our results further reveal keratinocytes play an active immunological role in response to sea lice. Given their capacity for antigen presentation through the expression of *MHCII*, superficial keratinocytes may play a sentinel role in the detection of sea lice and subsequently attract immune cells to the site of an attached sea lice. Superficial keratinocytes, and to a lesser extent intermediate keratinocytes also seem to be responsible for the dramatic increase in filament cell proliferation typifying coho salmon response to sea lice [17, 18] as evidenced by their upregulation of genes related to cell motility and filament reorganization. The intermediate keratinocytes, which we suggest lie between the superficial and basal keratinocytes due to their shared marker and differentially expressed genes, rapidly increase in abundance at 48-60h post sea lice infection and are likely responsible for the observed skin thickening in coho salmon in response to sea lice [17, 18]. Basal keratinocytes, alternatively, regulate the cell motility and proliferation of the upper layers of keratinocytes, strengthen the basement membrane of the epidermis, and emit antibacterial aerolysin proteins to prevent secondary microbial infections. Therefore, each layer of keratinocytes plays a unique but integrated role in the observed epithelial hyperplasia characterising coho salmon’s response to sea lice.

## CONCLUSIONS

In this study, we revealed the cell-specific mechanisms underlying responses to sea lice in a susceptible and a resistant salmonid species. Single nuclei RNA sequencing allowed us to identify the importance of genes with cell type-specific expression patterns, teasing apart cell-type specific responses, including variation in the functional roles among keratinocytes. Our results suggest a complex interplay of genes and cell types associated with sea lice response in both Atlantic salmon and coho salmon. The susceptibility of Atlantic salmon to sea lice infection despite clear activation of the complement, innate, and adaptive immune systems, confirms the insufficiency of this species immune response to effectively repel sea lice. Coho salmon, alternatively, demonstrate multiple interesting strategies in response to sea lice but keratinocytes seem to be key to the epithelial hyperplasia underlying coho salmon sea lice resistance.

The candidate genes we identified underlying coho salmon’s resistance and Atlantic salmon’s susceptibility hold significant promise for enhancing sea lice resistance in Atlantic salmon via biotechnological approaches such as gene editing. Knocking out genes in Atlantic salmon that we identified as upregulated during lice infestation and potentially linked to immunodeficiency and sea lice immunomodulation through CRISPR-Cas9 editing holds the potential to significantly enhance the species’ resistance to sea lice. Furthermore, promoting the expression of those genes associated with a dampened immune response in Atlantic salmon or those associated with epithelial hyperplasia in coho salmon could also effectively strengthen lice resistance in Atlantic salmon. Our findings thus offer actionable insights to mitigate the economic and ecological toll of sea lice infestations in the Atlantic salmon aquaculture industry.

## METHODS

### Experimental Design

Atlantic salmon eggs with poorer than average estimated breeding values for resistance to sea lice were sourced from Benchmark Genetics Iceland. Coho salmon (1 - 2 g) were provided by the Quinsam River Hatchery, Quinsam River, BC, Canada. Both species were reared in a Recirculating Aquaculture System at the Center for Aquaculture Technologies (PEI, Canada) in freshwater until post-smolt stage (approximately 15 g), after which fish were gradually transferred to saltwater and reared to a target weight of approximately 25 g. During the experiment, fish were kept in 135 L tanks at approximately 12 °C. Triplicate tanks of each species were treated with locally-sourced (n = 49 / fish, [147]) *Lepeophtheirus salmonis* copepodids and maintained for 60 hours and sampled every 12 hours. Untreated control fish were maintained in parallel tanks and sampled at 36 hours into the experiment. Fish were sedated before sampling with Tricaine methanesulfonate (100 mg L^-1^), and then subjected to a lethal blow to the head. Tissue samples (skin and pelvic fin), from louse attachment sites for treated fish, were collected and immediately frozen in dry ice.

### Library Preparation and Sequencing

Nuclei were isolated from one skin and one fin sample from each of the 5 treatment timepoints (12, 24, 36, 48, and 60 hours post exposure) as well as the control for each species (N = 24 tissue samples total) using a custom protocol optimized for salmon epidermis [168]. In brief, approximately 45 mg tissue samples were cut with scissors in 1 mL of TST buffer for 10 minutes on ice before being filtered through a 40 µm Falcon™ cell strainer (Thermo Fisher Scientific, catalog no. 08-771-2). A further 1 mL of TST and 3 mL of 1X PBS + BSA buffer were added to each sample before centrifuging at 4°C for 5 minutes at 500 g. Samples were resuspended in 1 mL 1X ST buffer filtered again through a 40 µm cell strainer, stained with Hoechst 33342 Solution (Thermo Fisher Scientific, catalog no. 62249) and then nuclei integrity was visually assessed using a fluorescent microscope. A disposable flow haemocytometer (C-Chip Neubauer Improved (100 µm depth), NanoEnTek, catalog no. DHC-N01) was then used to estimate nuclei counts.

Samples were processed with Chromium Next GEM Single Cell 3’ Reagent Kits v3.1 (Dual Index) (10X Genomics) using the protocol outlined in the User Guide (CG000315 Rev C). Samples were diluted with nuclease-free water to a target concentration that would recover approximately 7000 nuclei in the final library. Samples were then loaded on the Chromium Controller for nuclei droplet formation. After subsequent nuclei and UMI barcoding and reverse transcription, resulting cDNA was then amplified, fragmented, and indexed with Truseq adapters and Illumina sample indexes. Sequencing was performed on a NovaSeq 6000 platform (Illumina) by Azenta for approximately 220 million paired end 2×150bp reads per sample.

### Genome Indexing and Read Alignment with STAR

Genome indexing and library mapping was performed with STAR (version 2.7.10a, [169, 170]). We appended the mitochondrial genome from the ENSEMBL V2 Atlantic salmon genome (Salmo_salar.ICSASG_v2.dna_rm.toplevel.fa.gz, v2, release 105, masked genome, assembly ID: GCA_000233375.4) to the ENSEMBL V3 Atlantic salmon genome (Salmo_salar.Ssal_v3.1.dna_rm.toplevel.fa.gz, v3.1, release 106, masked genome, assembly ID: GCA_905237065.2) for both the .gff and .fna files prior to indexing. For coho salmon, we appended this species mitochondrial genome (version NC_009263.1, NCBI) to the ENSEMBL V2 coho salmon genome (Oncorhynchus_kisutch.Okis_V2.dna_rm.toplevel.fa.gz, v2, release 106, masked genome, assembly ID: GCA_002021735.2) for both the .gff and .fna files prior to indexing. Prior to this concatenation, the coho salmon mitochondrial genome .gff file was manually edited to convert “CDS” annotations to “exon” annotations (consistent with the Atlantic salmon mitochondrial genome .gff file) as STAR assigns transcripts to “exon” annotations in the .gff file. gffread (v0.10.1) was used to convert .gff to .gtf files [171]. Both genomes were indexed using STAR (--runMode genomeGenerate). Each library was then mapped against its corresponding genome with the 10X V3 cell barcode whitelist (3M-february-2018.txt) and using standard parameters for single cell libraries (--soloMultiMappers Unique --soloBarcodeReadLength 28 --soloType CB_UMI_Simple --soloUMIlen 12 -- soloCBwhitelist 3M-february-2018.txt --soloFeatures GeneFull --clipAdapterType CellRanger4 --outFilterScoreMin 30 --soloCBmatchWLtype 1MM_multi_Nbase_pseudocounts --soloUMIfiltering MultiGeneUMI_CR --soloUMIdedup 1MM_CR --readFilesCommand zcat --outSAMtype BAM Unsorted). The raw (unfiltered) files (*genes.tsv*, *barcodes.tsv*, and *matrix.mtx*) generated for each sample were then used for downstream analysis. On average, there were 300 million reads per sample with 94% of reads with valid barcodes, and a 62% saturation (for more details see Fig.S38, Table S1,S2).

### Quality Control, Clustering, Integration

Samples were then analysed in an R (v4) environment using Seurat (v4.1, [172]). We created Seurat objects for each library after removing nuclei with less than 200 features and features occurring in fewer than three nuclei. One Atlantic salmon sample (Atlantic_12h_fin) retained only 60 nuclei after this initial filtration and was therefore discarded from downstream analysis (Table S3). We then merged samples by species into a single Seurat object. Nuclei where mtDNA features accounted for 10% or more of their total UMIs were removed (Table S3, Fig.S39) before removing all mtDNA features (leaving 48608 and 39312 features remaining for Atlantic salmon and coho salmon, respectively). After sub-setting the Seurat object into individual samples, upper and lower thresholds for UMI and feature counts per nuclei were then applied individually to each sample based on knee plot visualization. For all Atlantic salmon samples, only nuclei with more than 500 UMIs but less than 6000 UMIs and more than 500 features and less than 3500 features were retained (Fig.S40). For coho salmon samples, a lower UMI and feature count limit of 300, 500, or 750 was applied to each sample; an upper UMI limit of 2000 or 6000 was applied while an upper feature limit of 1500 or 3500 was applied (Fig.S41). A single Atlantic salmon sample (Atlantic_24h_fin) retained only 338 nuclei after this initial filtration and was therefore discarded from downstream analysis (Table S3).

Samples were then merged again into a single Seurat object by species before splitting samples again into individual sample datasets. This was done to ensure that the same features were considered across samples. Counts were then normalized for each sample using the “NormalizeData” function prior to calculating cell cycle scores using the “CellCycleScoring” function (see Tables S8,S9 for list of genes used). The “v2” SCTransform version with the glmGamPoi method (v 1.8.0, [173]) was used to normalize RNA counts for each sample, regressing out scores for the S and G2M cell cycle stages. Linear dimension reduction was conducted for each sample using the “RunPCA” function with 50 PCs. After consulting Elbowplots for each sample, a UMAP using 20 PCs was run for each sample and the “FindNeighbours” function was applied using 20 PCs, before using the “FindClusters” function with a resolution of 0.2. DoubletFinder (v 2.0.3, [174]) was then applied independently to each sample selecting pK values with the highest associated BCmvn value. We assumed a 4% doublet formation rate (based on the Chromium instrument specifications) and adjusted for homotypic doublets (see Table S3 for remaining cells per sample after doublet removal).

Samples were integrated by species using 5000 features and anchors that were identified with the “rpca” reduction method and the “FindIntegrationAnchors” function. A PCA was rerun on the integrated dataset using 50 PCs, and 30 PCs were used for subsequent UMAP generation and clustering with a resolution of 0.2 (Fig.S42a,S43a). Markers for each cluster were assessed using the logistic regression method and the FindAllMarkers function on the “SCT” assay and “data” slot, using sample ID as a latent variable to help reduce batch effects among samples. We used a pseudocount of 0.001, set a p-value threshold of 0.01, and only considered genes that were upregulated, expressed in at least 25% of all nuclei (in either of the compared groups), and demonstrated the default threshold of 0.25 X difference (log-scale) between the two compared groups.

Two clusters (0 and 4) were removed from the Atlantic salmon dataset due to low average feature/UMI counts (Fig.S42c,d). Many of the marker genes for cluster 0 were ribosomal genes, suggesting poor quality nuclei (Fig.S44). Cluster 4 was also found almost exclusively in a single sample (Atlantic_Control_skin), again suggesting it was poor quality (Fig.S42b). Similarly, cluster 1 from the coho salmon dataset was removed for having low average feature/UMI counts and because many of its markers were ribosomal genes (Fig.S43c,d,S45). The SCTransformation was then redone for each sample based on the RNA assay as described above, and integration of samples for each species was conducted as described above using 30 PCs for UMAP generation and a resolution of 0.2 for clustering for Atlantic salmon and 20 PCs for UMAP generation and a resolution of 0.2 for clustering for coho salmon. An additional cluster (11) was subsequently removed from the coho salmon dataset for having many ribosomal marker genes (Fig.S46,S47). The SCTransformation of each sample and integration of samples was again redone for the coho salmon dataset after removing this cluster, again using 20 PCs for UMAP generation and a resolution of 0.2 for clustering. (See Table S3 for remaining cells per sample and Fig.S48 for the distribution of UMIs and features per sample after all filtering.)

### Sub-clustering

Clusters identified as immune cells based on the expression (Fig.S8) of *cd45* (*ptprc*) (a marker gene for immune cells, [96]) were then considered separately for each species to investigate for the presence of additional immune cell types. For immune cells identified within Atlantic salmon samples, a PCA was rerun on the integrated assay using 10 PCs, and UMAP generation and clustering were conducted using 9 PCs and a resolution of 0.3, respectively. For coho salmon immune cells, a PCA was rerun on the integrated dataset using 20 PCs, UMAP was generated using 15 PCs and clustering was conducted using a resolution of 0.4. Marker genes comparing each immune cell cluster with all other immune cells were then identified using the same marker gene detection method described above using the “FindAllMarkers” function but UMI counts were not re-corrected based on the sub-setted datasets (recorrect_umi = FALSE). Marker genes were investigated and visualized to assess cell type. All clusters identified as macrophages were grouped together as were all clusters identified as T-Cells (see Results).

Within a single cluster of the coho salmon dataset (cluster 12) we observed expression of the *ltk* gene (a marker of neural crest cells in Atlantic salmon, see results below) in a small subset of cells within this cluster while other cells within this cluster demonstrated expression of *casq1b* (a marker of muscle cells in Atlantic salmon, see results below) (Fig.S49a,b). To investigate the potential for multiple cell types within this cluster, we reran a PCA on cells from this cluster using the integrated assay and 10 PCs, before performing UMAP generation using 3 PCs and clustering with a resolution of 0.02. The resulting UMAP revealed two clusters of cells, one expressing *ltk*, the other expressing *casq1b* (Fig.S49c-f).

These detected subclusters were then incorporated into the larger dataset for each species including all cell types (see Fig.S50 for distribution of UMIs and features per cluster). Marker genes were then assessed for all newly identified immune cell types using the “FindAllMarkers” function (as described above) in the context of all other cell types. The top markers based on the average log 2-fold change were then considered for each cluster to assess cell type identity. Gene annotations from the ENSEMBL genome were supplemented with EntrezID (NCBI, [175]) and UniProt [176] annotations based on querying BioMart (v 2.50.3, [177]).

### Differential Gene Expression Detection

We next identified genes which were differentially expressed between the control samples and each of the infection timepoints (12, 24, 36, 48, 60 hrs post infection) for both species and all cell types using the “FindMarkers” function and the default Wilcox method. We used the SCT assay and “data” slot, imposed a minimum percent threshold (percent of cells in either considered group that had to express the gene) of 0.1, set a minimum threshold p-value of 0.01, and used the default threshold of 0.25 X difference (log-scale) between the two compared groups. We excluded results from cell types that had fewer than 50 nuclei in the control samples and comparisons where the treatment timepoint had fewer than 50 nuclei. Genes were considered differentially expressed if their adjusted p-value < 0.001. Enriched GO Biological Processes for differentially expressed genes detected for each cell type for each species were identified using ShinyGO (v 0.80, [178]). We used default parameters and limited the gene universe to all features in the RNA assay for each species (N = 48608, N = 39312 genes for Atlantic salmon and coho salmon, respectively). GO terms were considered significantly enriched if the FDR-adjusted p-value < 0.001.

### Integration of Samples Across Species

We then directly compared Atlantic salmon and coho salmon samples using 6494 genes identified using Orthofinder v2.5.4 [179] as 1:1 orthologs between the two species. The transcriptomes of the Atlantic salmon and coho salmon Ensembl genomes used as reference for the snRNAseq analyses were used (Salmo_salar.Ssal_v3.1.cdna.all.fa and Oncorhynchus_kisutch.Okis_V2.cdna.all.fa). A single isoform per gene was retained using a custom python script that selects the longest transcript for each gene, and Orthofinder was run using default parameters. The orthogroups with one gene per species were considered 1:1 orthologs between Atlantic salmon and coho salmon.

Atlantic salmon and coho salmon samples were re-processed using the same quality control methods as described above, but features were winnowed down to this set of 1:1 orthologous genes just prior to the SCTransformation of individual samples. Samples from both species were then integrated together using 2000 features using anchors identified with the “rpca” reduction method with the “FindIntegrationAnchors” function. A PCA was run on the integrated dataset using 50 PCs with clustering and a UMAP was generated using 20 PCs and a resolution of 0.2. Markers were then detected for each cluster and species using the “FindAllMarkers” function as described above. The distribution of features and UMIs as well as the top markers based on the average log 2-fold change were then considered for each cluster. A single cluster (cluster 0) was removed due to a lack of defining marker genes (Fig.S51,S52), following reclustering as above a second cluster (cluster 1) was again removed due to a lack of defining marker genes (Fig.S53,S54). After removing these clusters the SCTransformation was redone for each sample based on the RNA assay, and integration of samples for each species was conducted as described above (using 2000 features for integration, 50 PCs for the PCA, 20 PCs and a resolution of 0.2 for clustering and UMAP generation, see Fig.S55 for distribution of UMIs and features per cell type and cell type counts per sample). Markers were then detected for each cluster using the “FindAllMarkers” function as described above. The top markers based on the average log 2-fold change were then considered for each cluster to assess cell type identity.

### Ethics Approval and Consent to Participate

CATC and UPEI Animal Care Committees (AUP 21-008) approved all fish handling procedures, which were conducted in accordance with the Canadian Council for Animal Care regulations (http://www.ccac.ca/) and ARRIVE guidelines.

### Consent for Publication

Not applicable.

### Availability of Data and Materials

Sequencing data for all samples used in this study will be deposited to NCBI SRA upon article acceptance. Scripts used to analyse and visualize data are available at https://github.com/SarahSalisbury/Atlantic_Salmon_vs_Coho_Salmon_Lice_Response_snRNAseq.

## Authors’ Contributions

MDF, SJM, JEB, MDF, RDH, NR, and DR created the experimental design. MDF, RRD, and DR conducted the experiment and collected tissue samples. RRD led the lab work with assistance from SJS, PRV, and OG. SJS led the data analysis, interpretation, and figure generation, with assistance from DR. SJS led the writing in consultation with DR and with assistance from RRD, SJM, JEB, PRV, MDF, LS, RDH, and NR. All authors approved the final manuscript.

## Acknowledgements

We gratefully acknowledge Paige Ackerman, Edward Walls, Sherri Higgins, Cindy Ederis for contributing coho salmon from Quinsam River Hatchery as well as Ólafur H. Kristjánsson of StofnFiskur, Benchmark Genetics in Iceland and Morten Rye of Benchmark Genetics in Norway for contributing Atlantic salmon eggs. Our thanks go to Dr. Mark Polinksi and Russell Anderson from FAS delivery for ensuring the safe and efficient transfer of animals; and Dr. Mark Braceland and Dr. Hiatham Mohammed from CATC, as well as Dylan Michaud for ensuring the welfare of animals throughout the experiment.

