## Supplementary Material for "Keratinocytes Drive the Epithelial Hyperplasia Key to Sea Lice Resistance in Coho Salmon"

**Fig.S1** Expression of marker genes within 23 identified cell clusters within Atlantic salmon fin and skin samples.

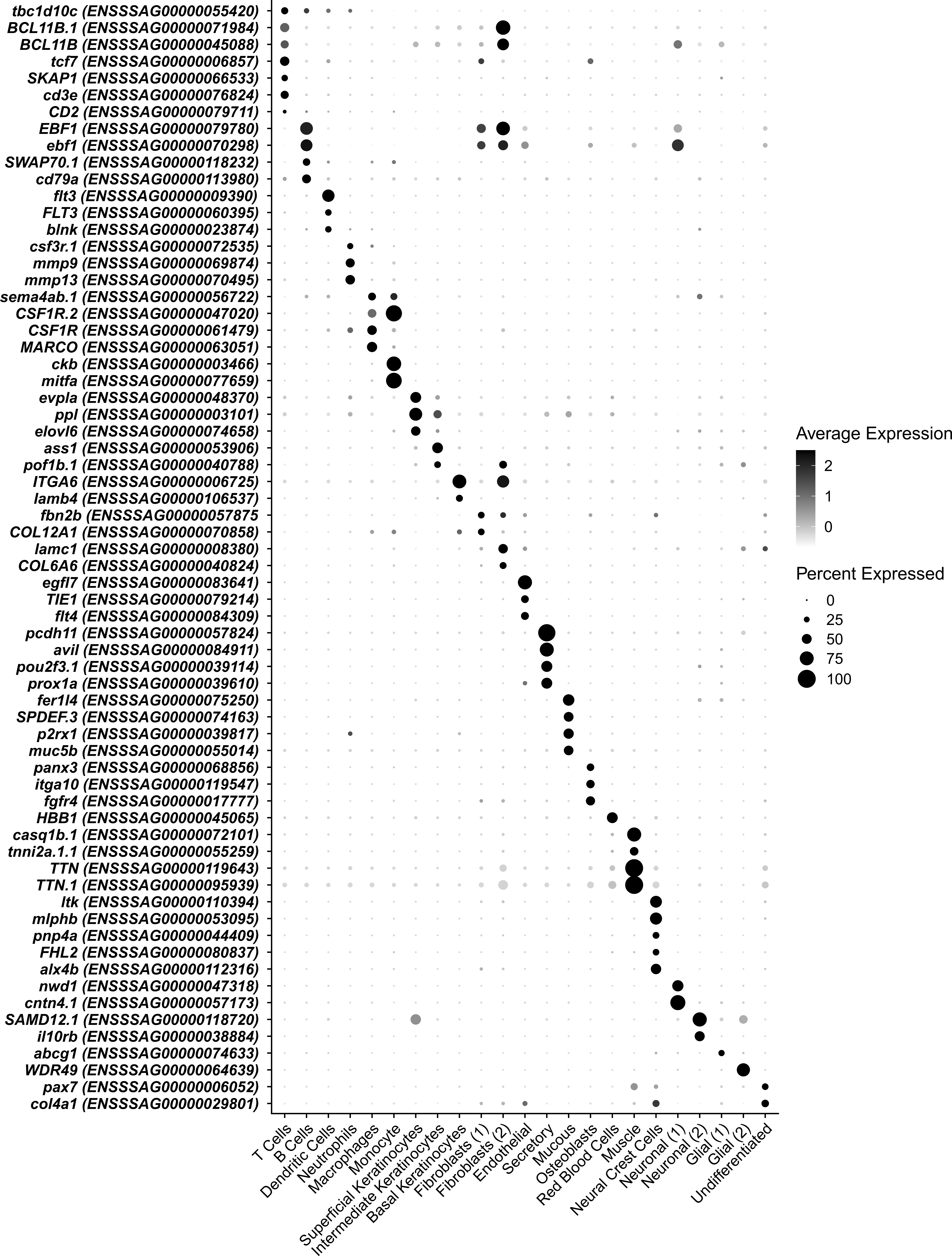

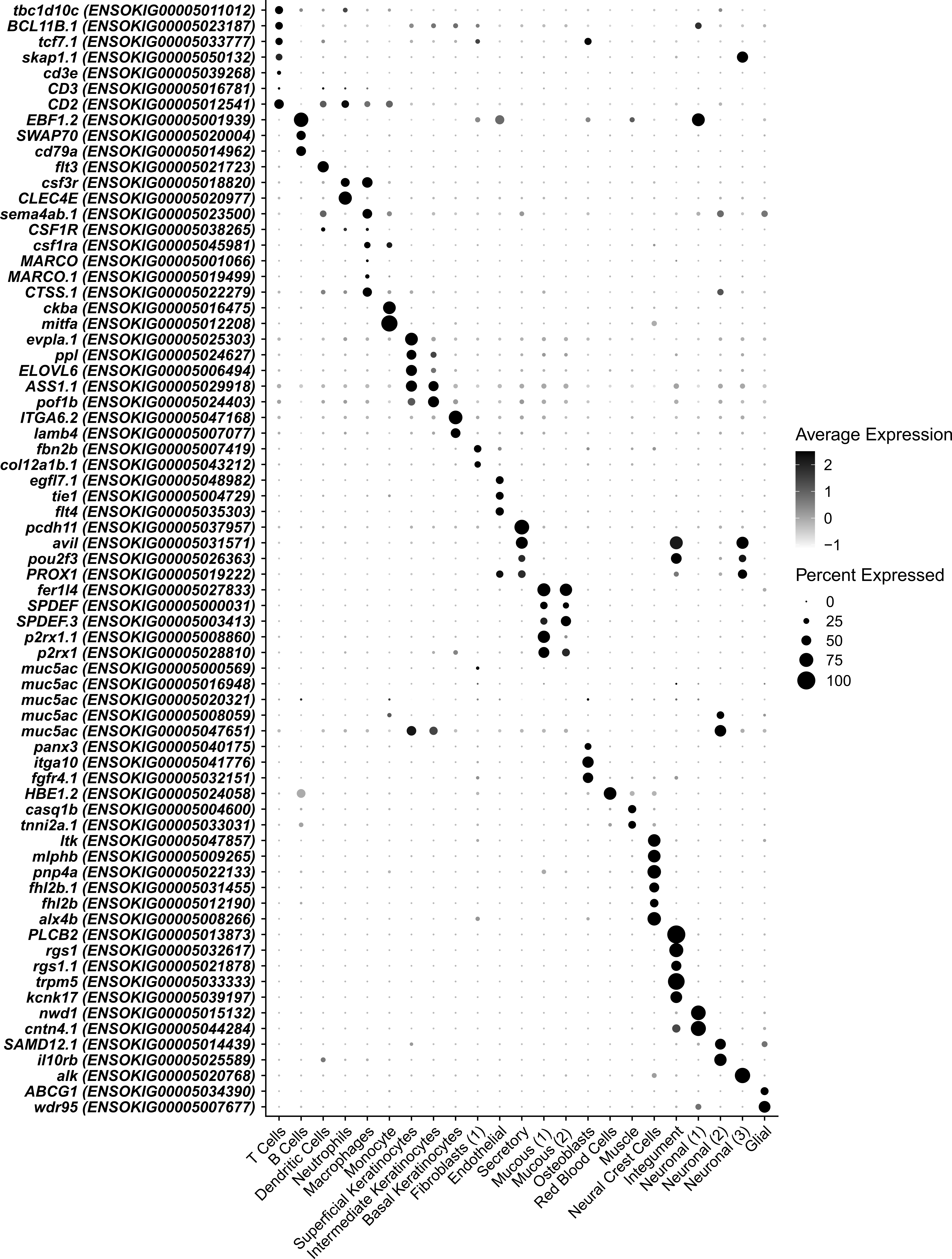

**Fig.S2** Expression of marker genes within 23 identified cell clusters within coho salmon fin and skin samples.

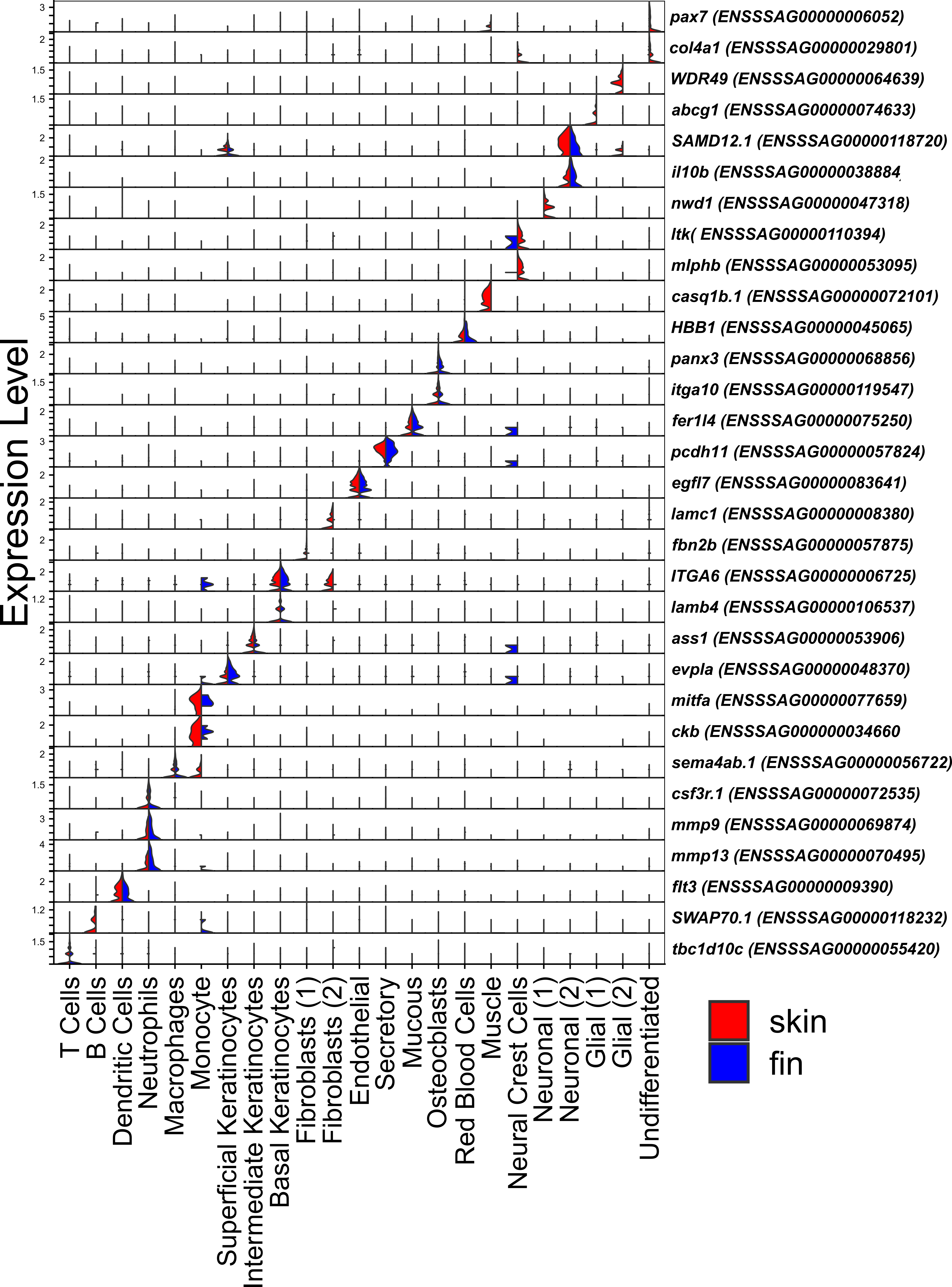

**Fig.S3** Violin plots of the expression of marker genes from Fig.1c for each cell type detected within Atlantic salmon samples split by tissue type (skin or pelvic fin). Note that the cell types: red blood cells, muscle, neural crest cells, neuronal (1), and glial cell clusters were almost exclusively observed in skin samples.

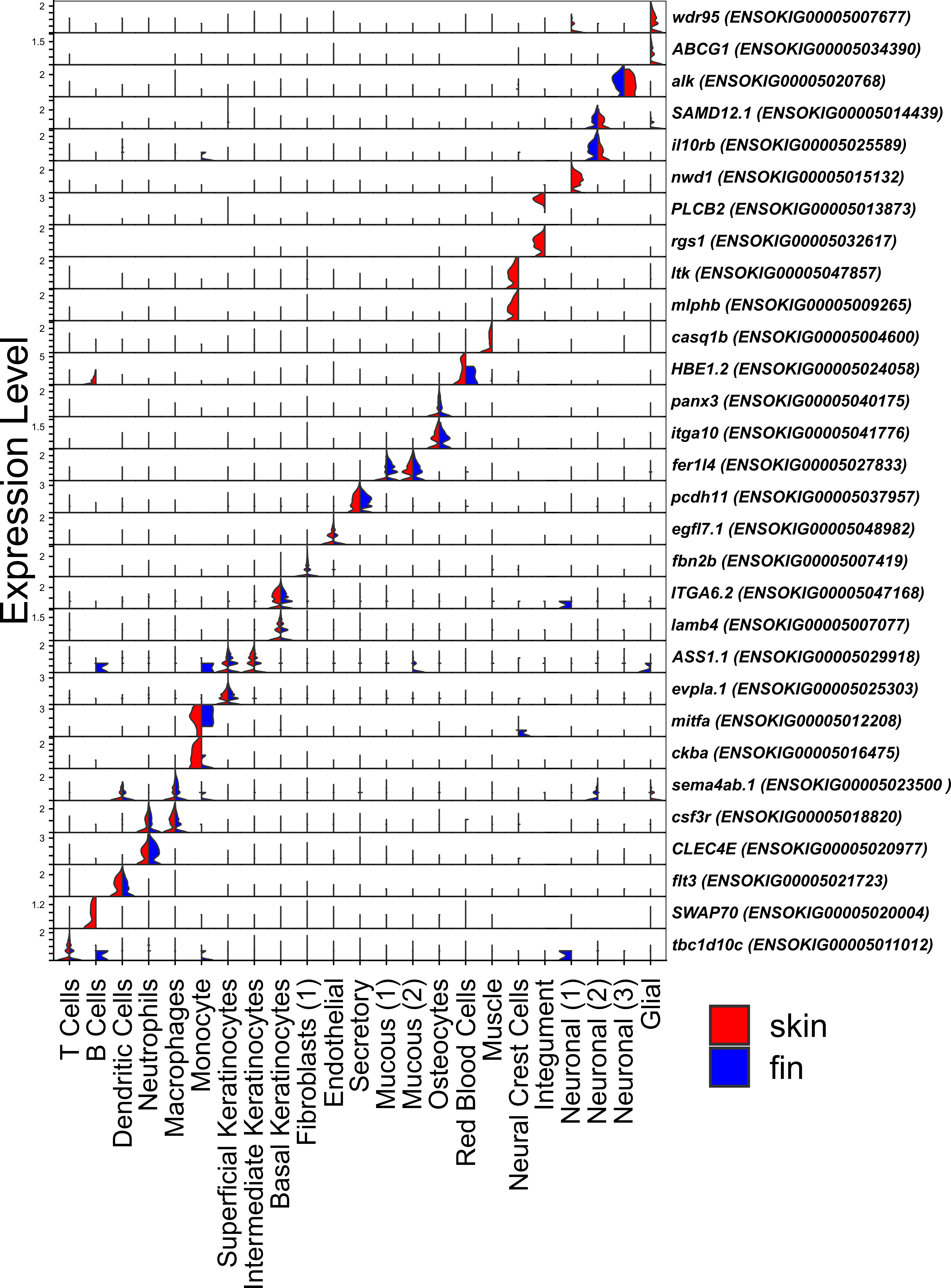

**Fig.S4** Violin plots of the expression of marker genes from Fig.1d for each cell type detected within coho salmon samples split by tissue type (skin or pelvic fin). Note that the cell types: red blood cells, muscle, neural crest cells, neuronal (1), and glial cell clusters were almost exclusively observed in skin samples.

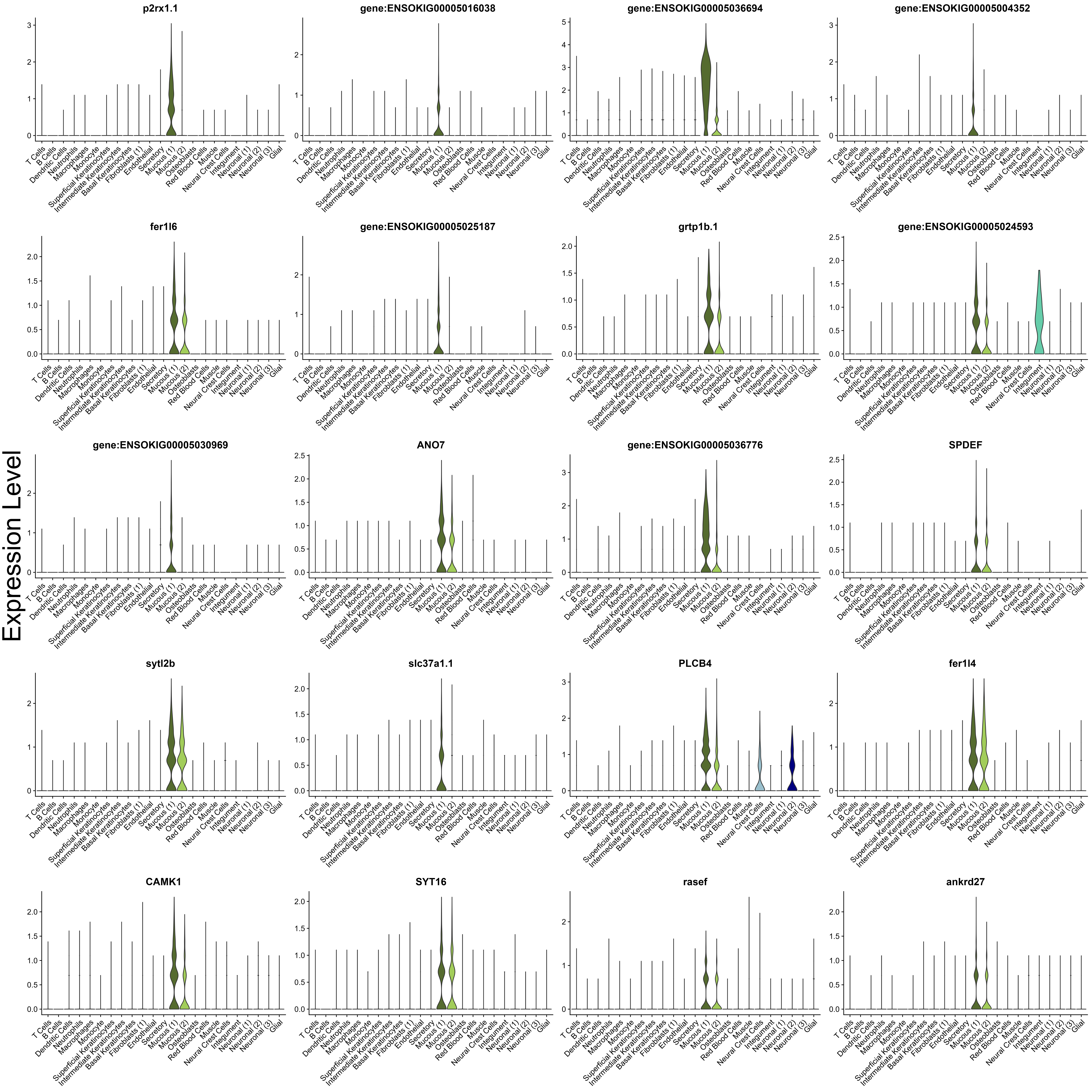

**Fig.S5** Violin plots of expression levels (based on the SCT assay) for the top 20 significant (adjusted p-value < 0.05) marker genes for the mucous (1) cluster (based on log-scale two-fold change in expression) of the coho salmon dataset (for UMAP see Fig.1).

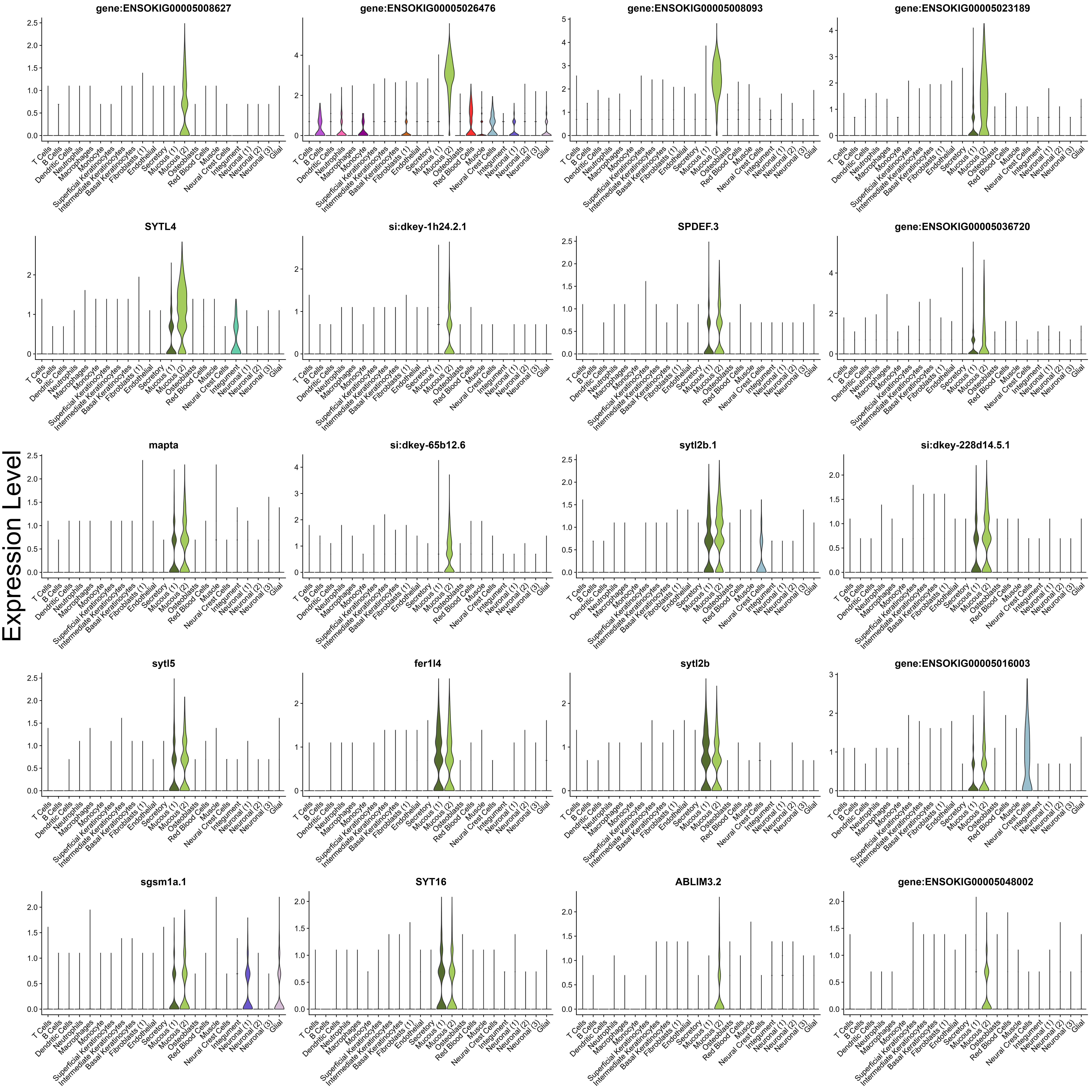

**Fig.S6** Violin plots of expression levels (based on the SCT assay) for the top 20 significant (adjusted p-value < 0.05) marker genes for the mucous (2) cluster (based on log-scale two-fold change in expression) of the coho salmon dataset (for UMAP see Fig.1).

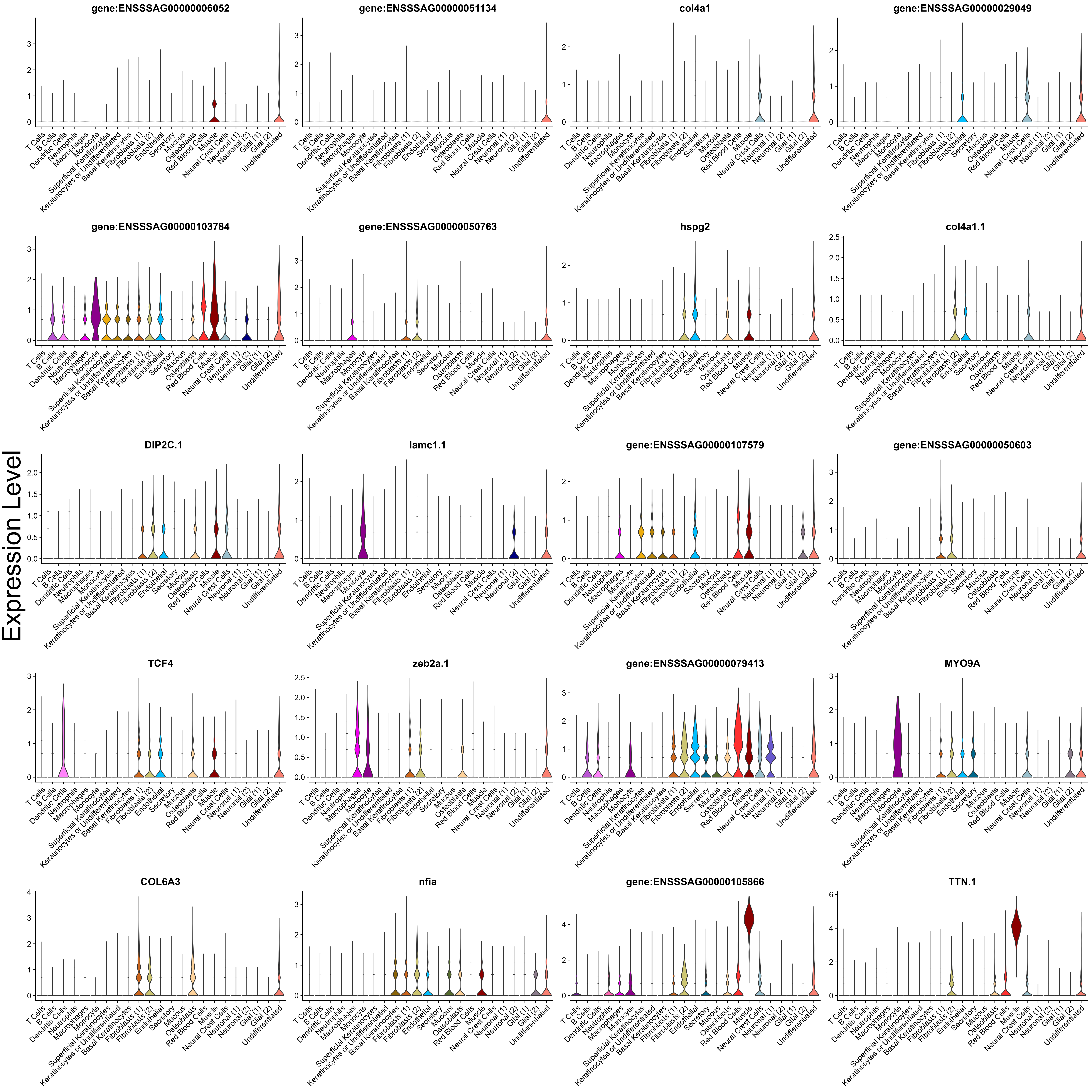

**Fig.S7** Violin plots of expression levels (based on the SCT assay) for the top 20 significant (adjusted p-value < 0.05) marker genes for the undifferentiated cluster (based on log-scale two-fold change in expression) of the Atlantic salmon dataset (for UMAP see Fig.1).

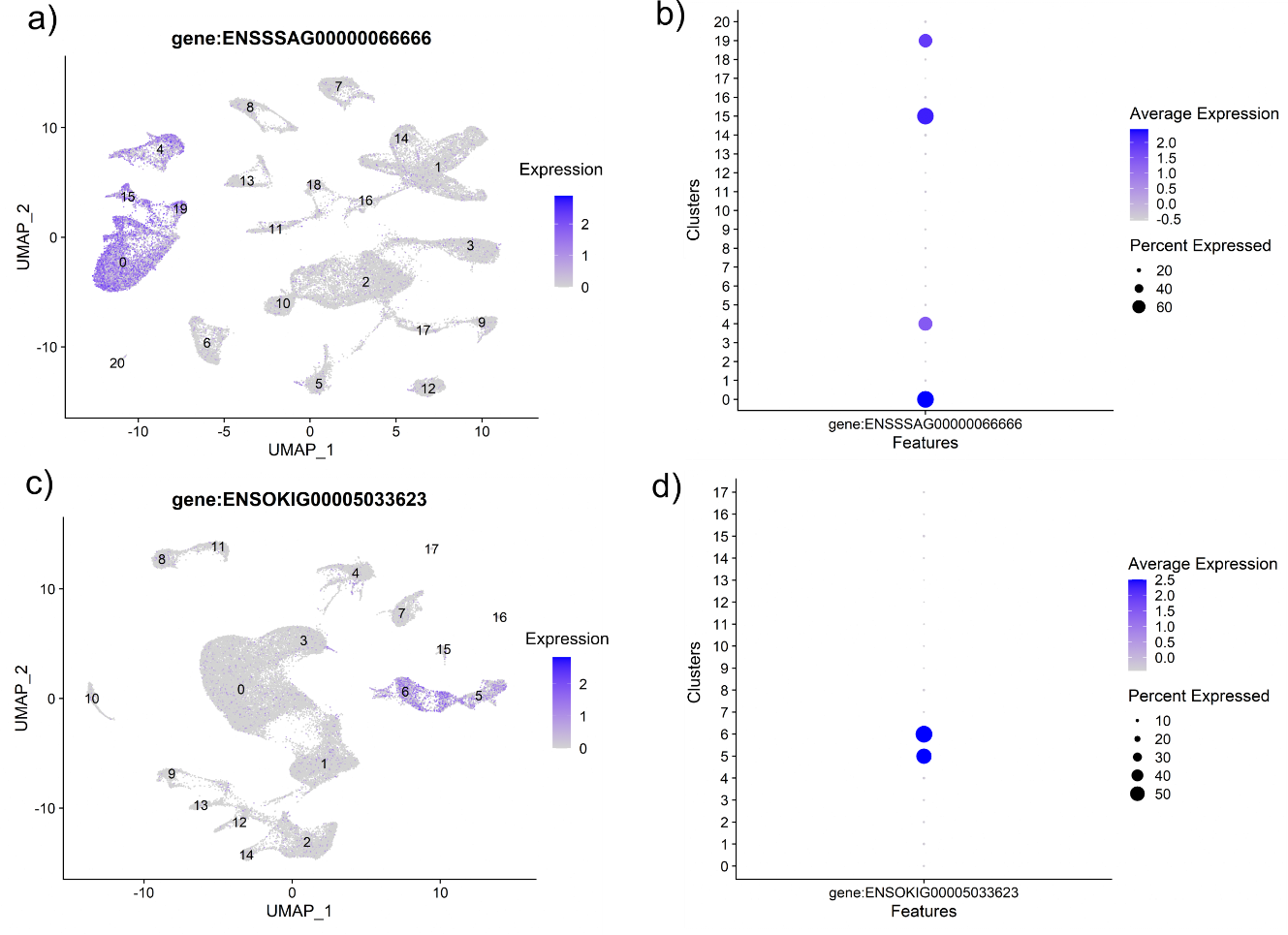

**Fig.S8** Expression of *CD45* (*ptprc*) in Atlantic salmon (a, b) and coho salmon (c, d). Feature Plots (a, c) indicate expression (based on the SCT assay) of *CD45*. Dotplots (b, d) indicate average expression (based on the SCT assay) of *CD45* for each cluster.

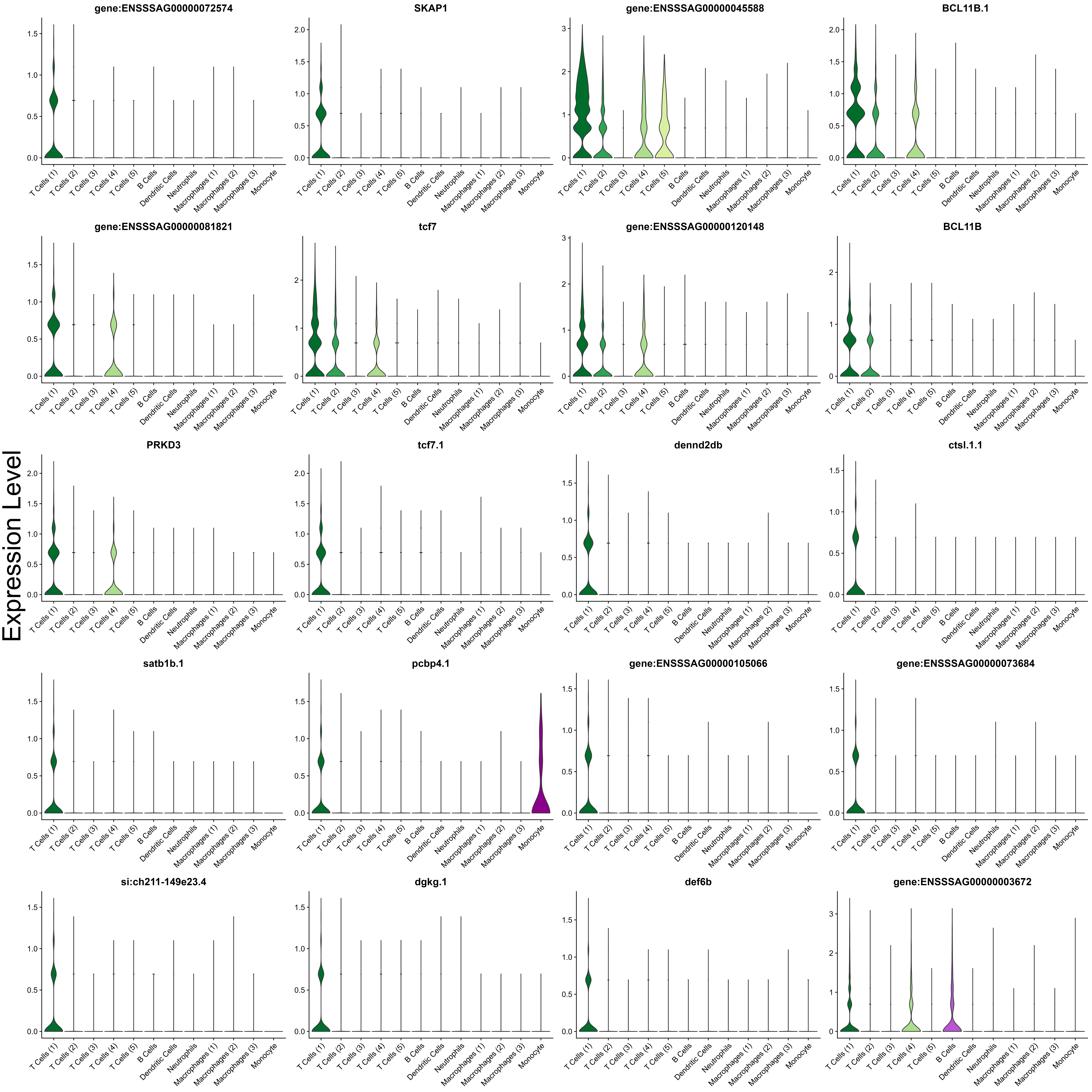

**Fig.S9** Violin plots of expression levels (based on the SCT assay) for the top 20 significant (adjusted p-value < 0.05) marker genes for the T cells (1) cluster (based on log-scale two-fold change in expression) of the Atlantic salmon immune cells only data subset (for UMAP see Fig.3a).

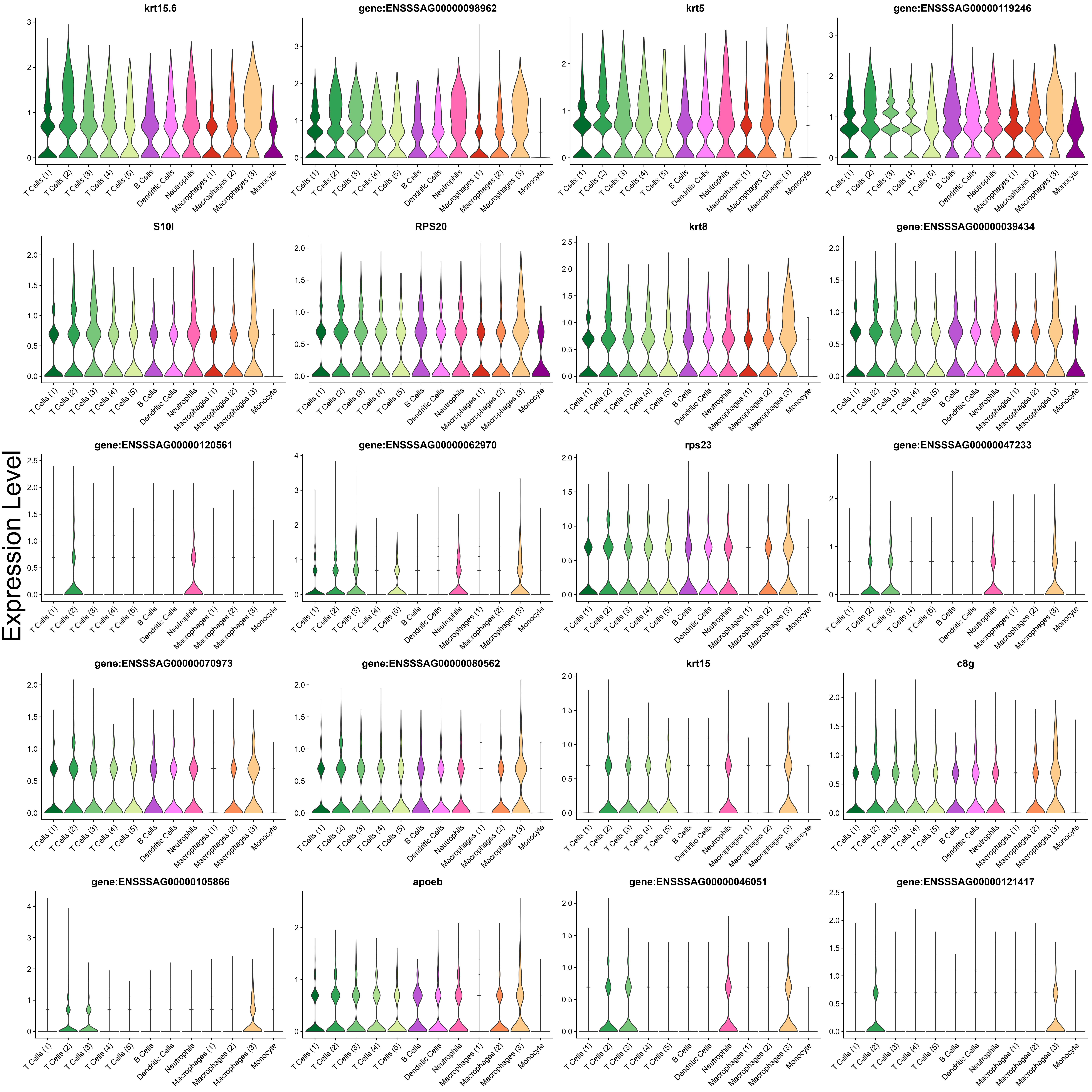

**Fig.S10** Violin plots of expression levels (based on the SCT assay) for the top 20 significant (adjusted p-value < 0.05) marker genes for the T cells (2) cluster (based on log-scale two-fold change in expression) of the Atlantic salmon immune cells only data subset (for UMAP see Fig.3a).

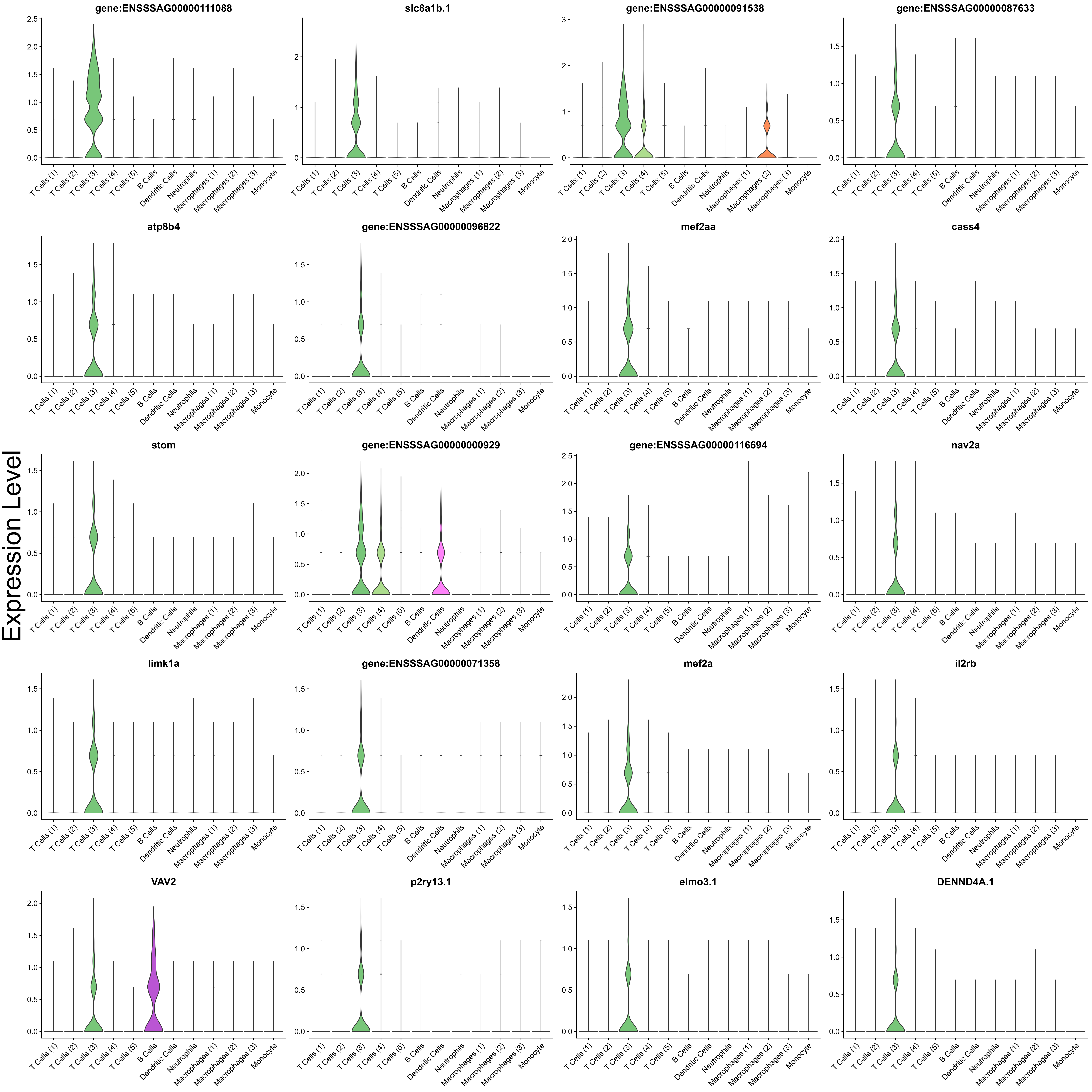

**Fig.S11** Violin plots of expression levels (based on the SCT assay) for the top 20 significant (adjusted p-value < 0.05) marker genes for the T cells (3) cluster (based on log-scale two-fold change in expression) of the Atlantic salmon immune cells only data subset (for UMAP see Fig.3a).

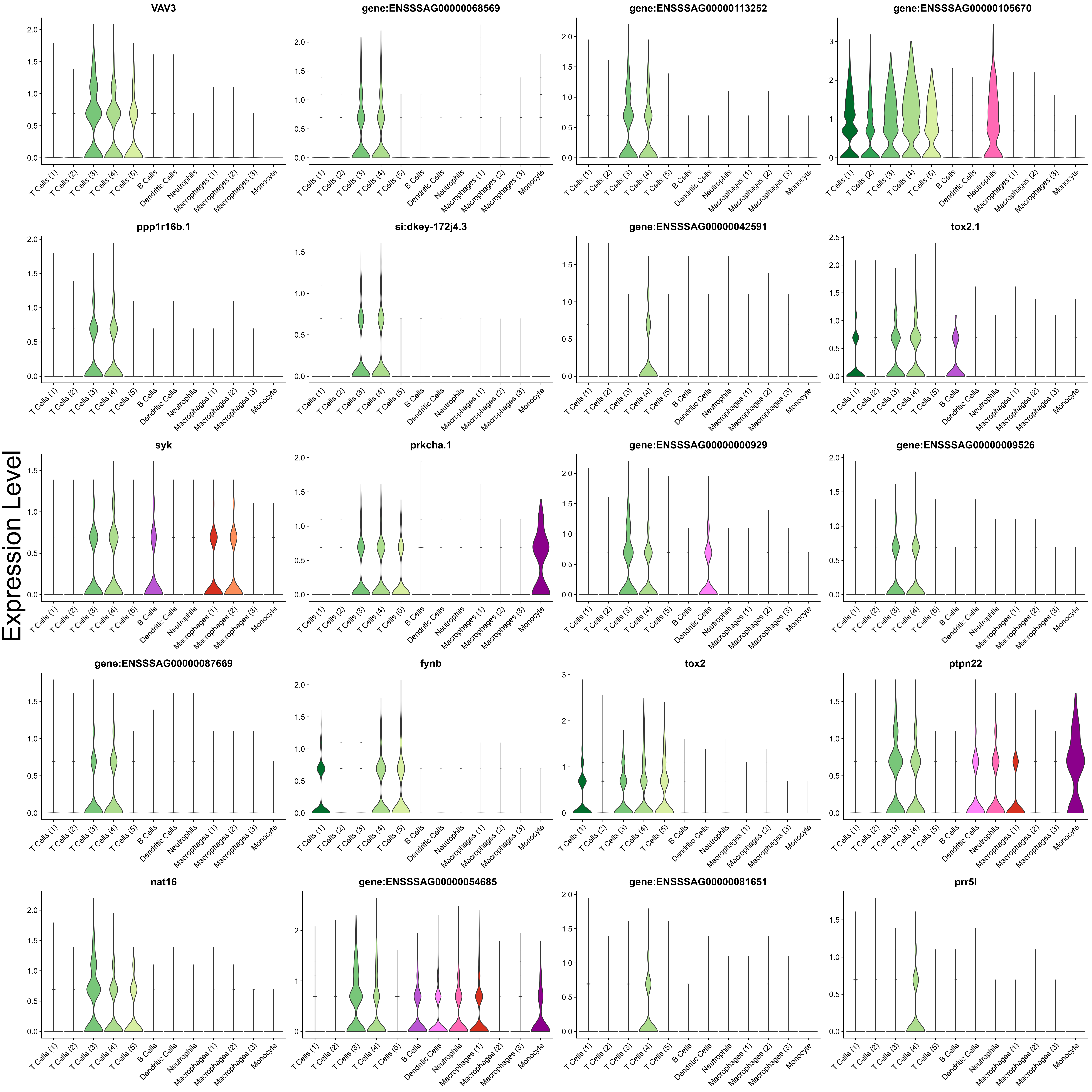

**Fig.S12** Violin plots of expression levels (based on the SCT assay) for the top 20 significant (adjusted p-value < 0.05) marker genes for the T cells (4) cluster (based on log-scale two-fold change in expression) of the Atlantic salmon immune cells only data subset (for UMAP see Fig.3a).

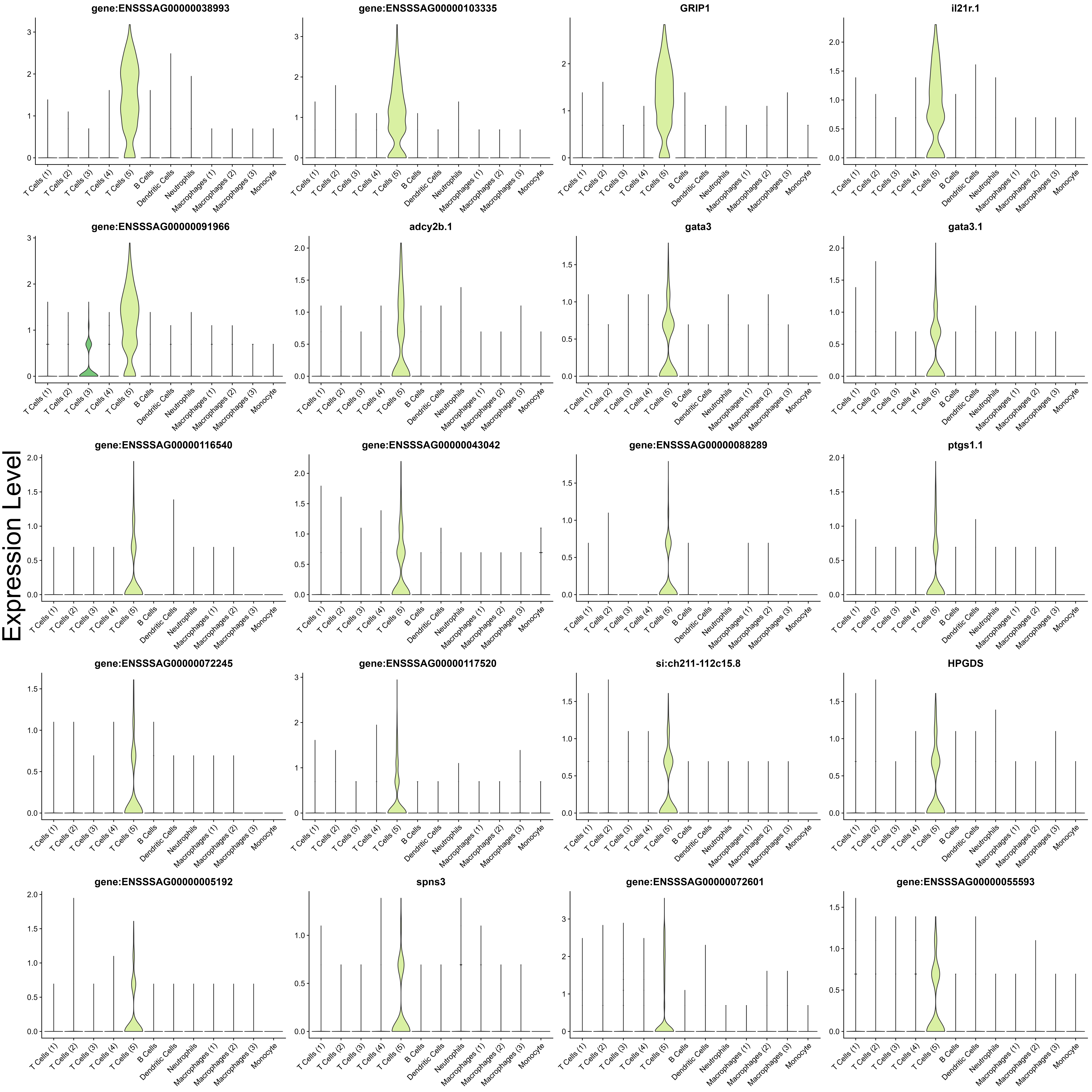

**Fig.S13** Violin plots of expression levels (based on the SCT assay) for the top 20 significant (adjusted p-value < 0.05) marker genes for the T cells (5) cluster (based on log-scale two-fold change in expression) of the Atlantic salmon immune cells only data subset (for UMAP see Fig.3a).

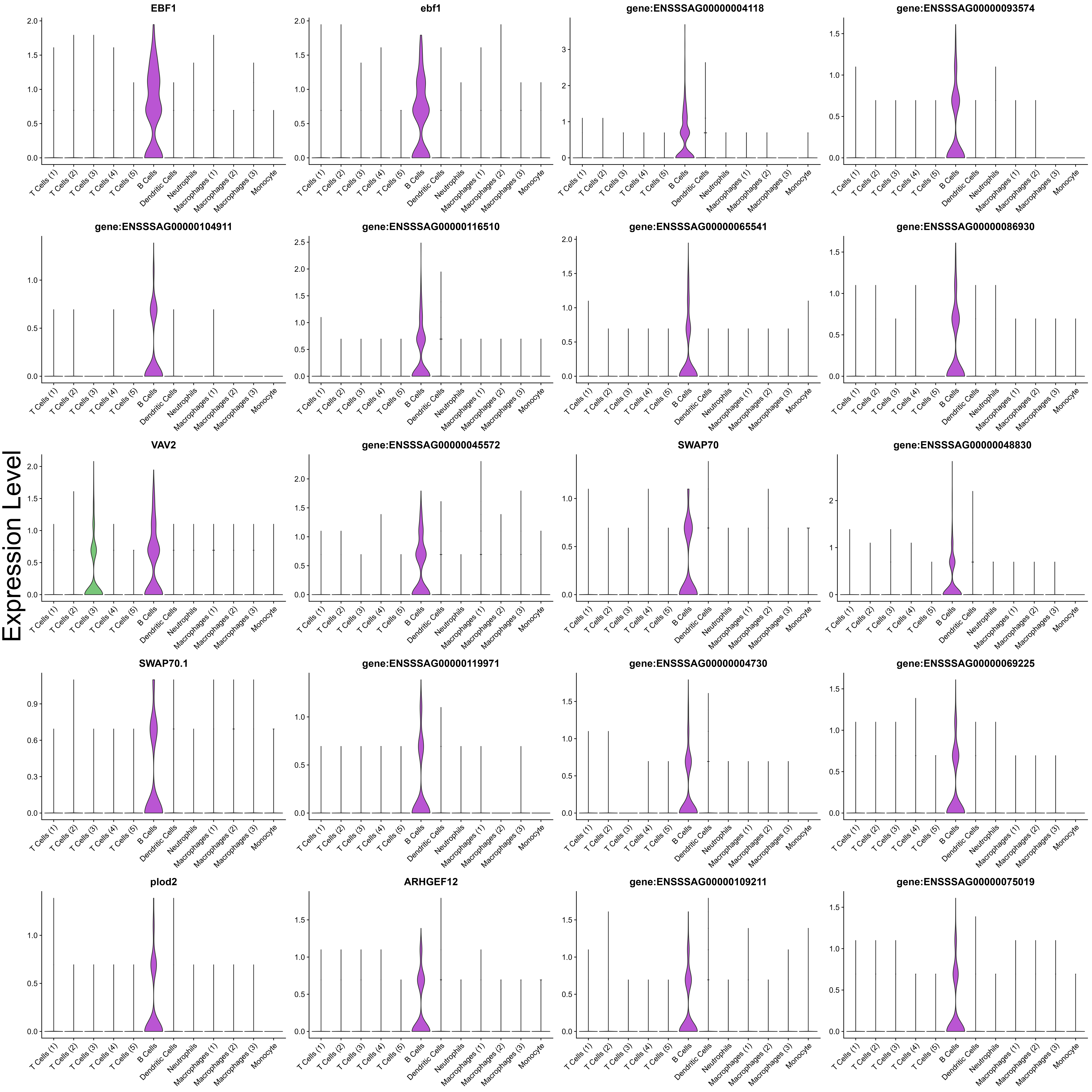

**Fig.S14** Violin plots of expression levels (based on the SCT assay) for the top 20 significant (adjusted p-value < 0.05) marker genes for B cells (based on log-scale two-fold change in expression) of the Atlantic salmon immune cells only data subset (for UMAP see Fig.3a).

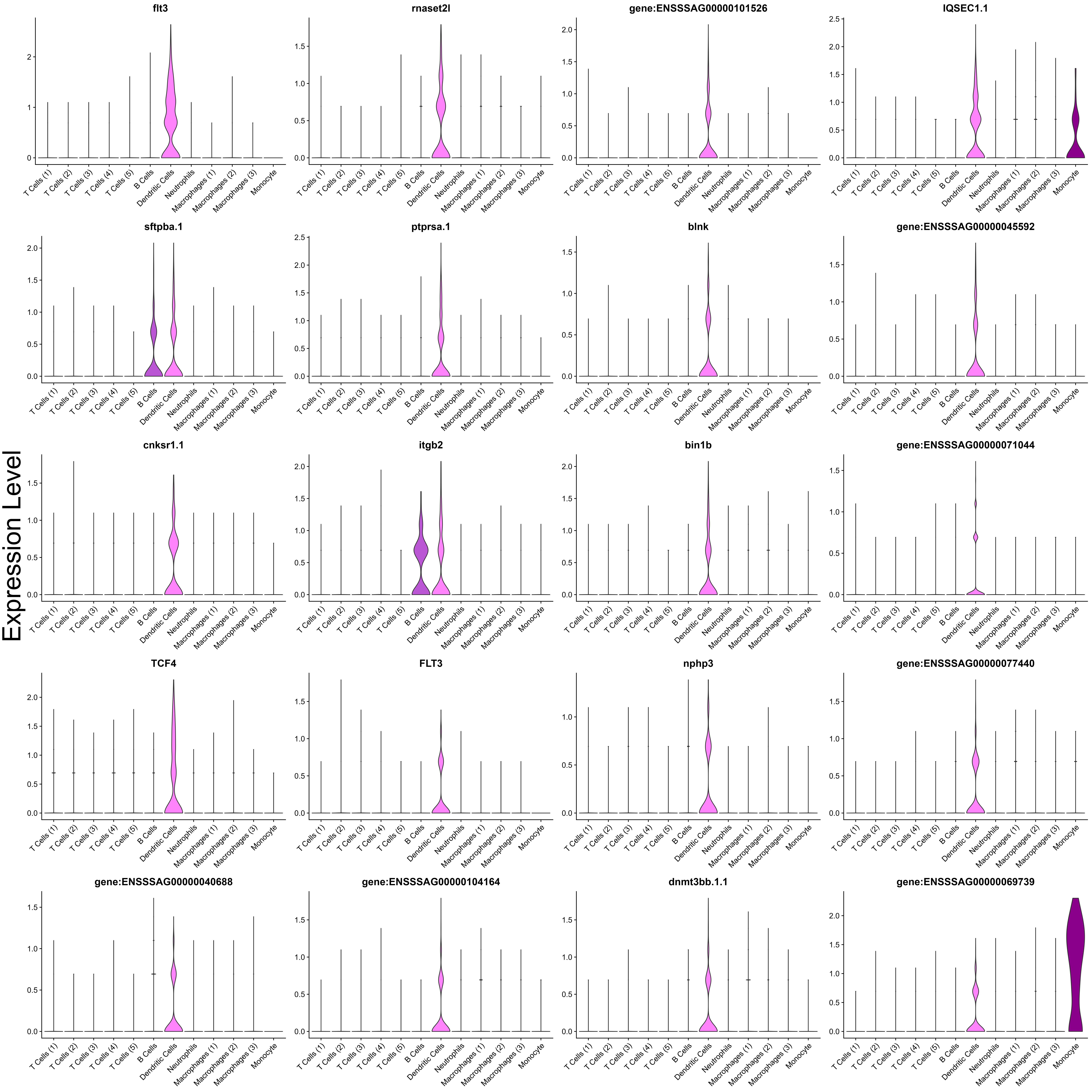

**Fig.S15** Violin plots of expression levels (based on the SCT assay) for the top 20 significant (adjusted p-value < 0.05) marker genes for dendritic cells (based on log-scale two-fold change in expression) of the Atlantic salmon immune cells only data subset (for UMAP see Fig.3a).

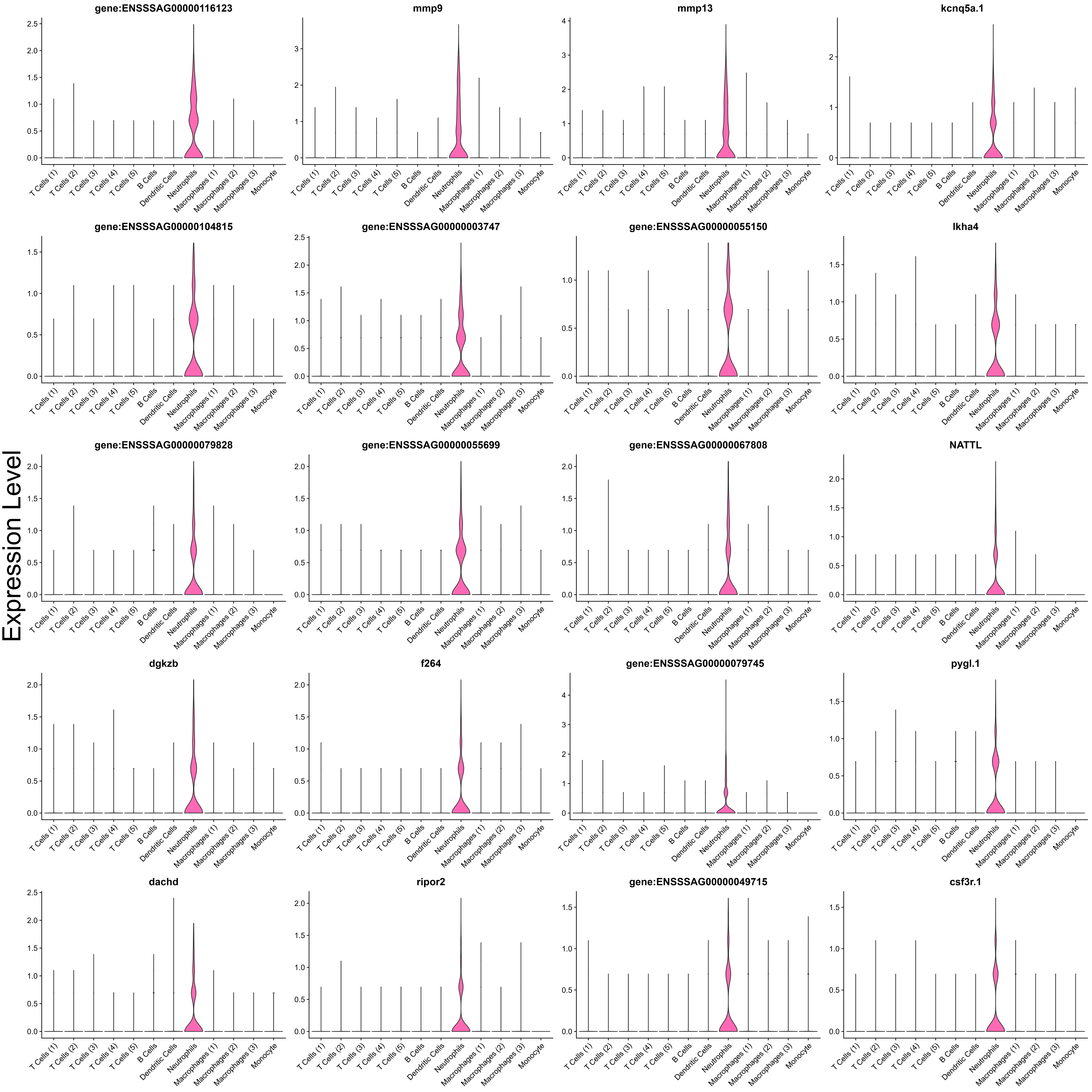

**Fig.S16** Violin plots of expression levels (based on the SCT assay) for the top 20 significant (adjusted p-value < 0.05) marker genes for neutrophils (based on log-scale two-fold change in expression) of the Atlantic salmon immune cells only data subset (for UMAP see Fig.3a).

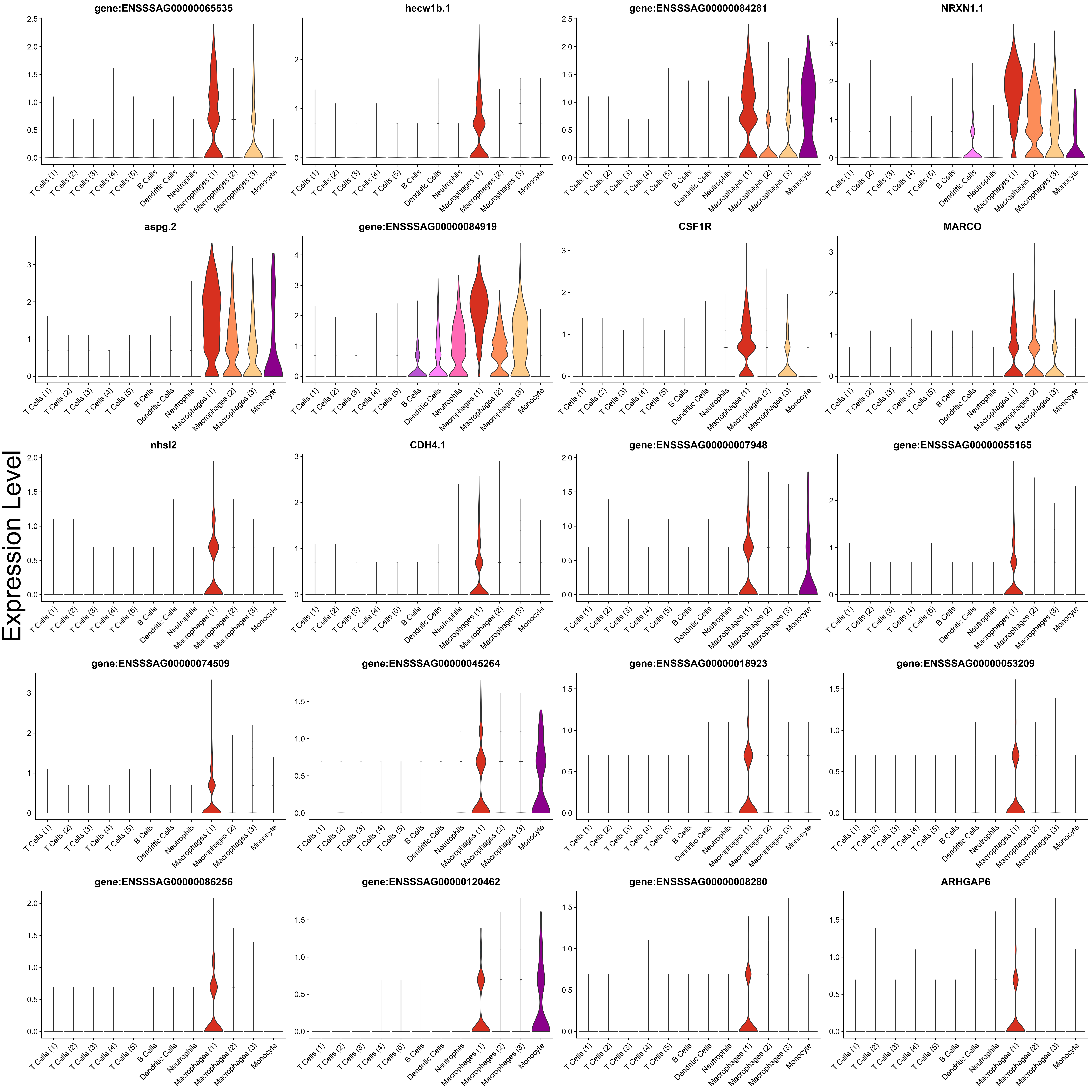

**Fig.S17** Violin plots of expression levels (based on the SCT assay) for the top 20 significant (adjusted p-value < 0.05) marker genes for the macrophages (1) cluster (based on log-scale two-fold change in expression) of the Atlantic salmon immune cells only data subset (for UMAP see Fig.3a).

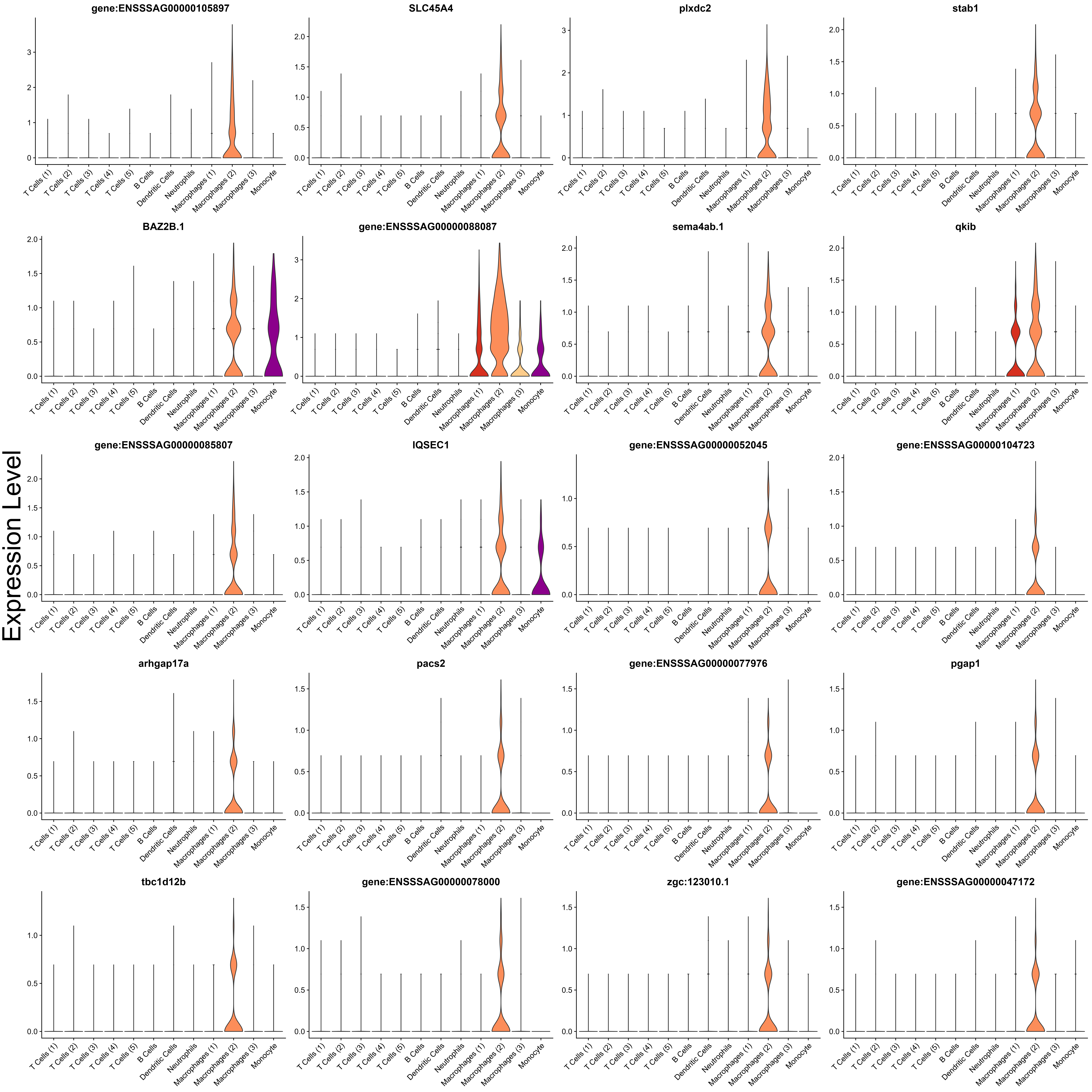

**Fig.S18** Violin plots of expression levels (based on the SCT assay) for the top 20 significant (adjusted p-value < 0.05) marker genes for the macrophages (2) cluster (based on log-scale two-fold change in expression) of the Atlantic salmon immune cells only data subset (for UMAP see Fig.3a).

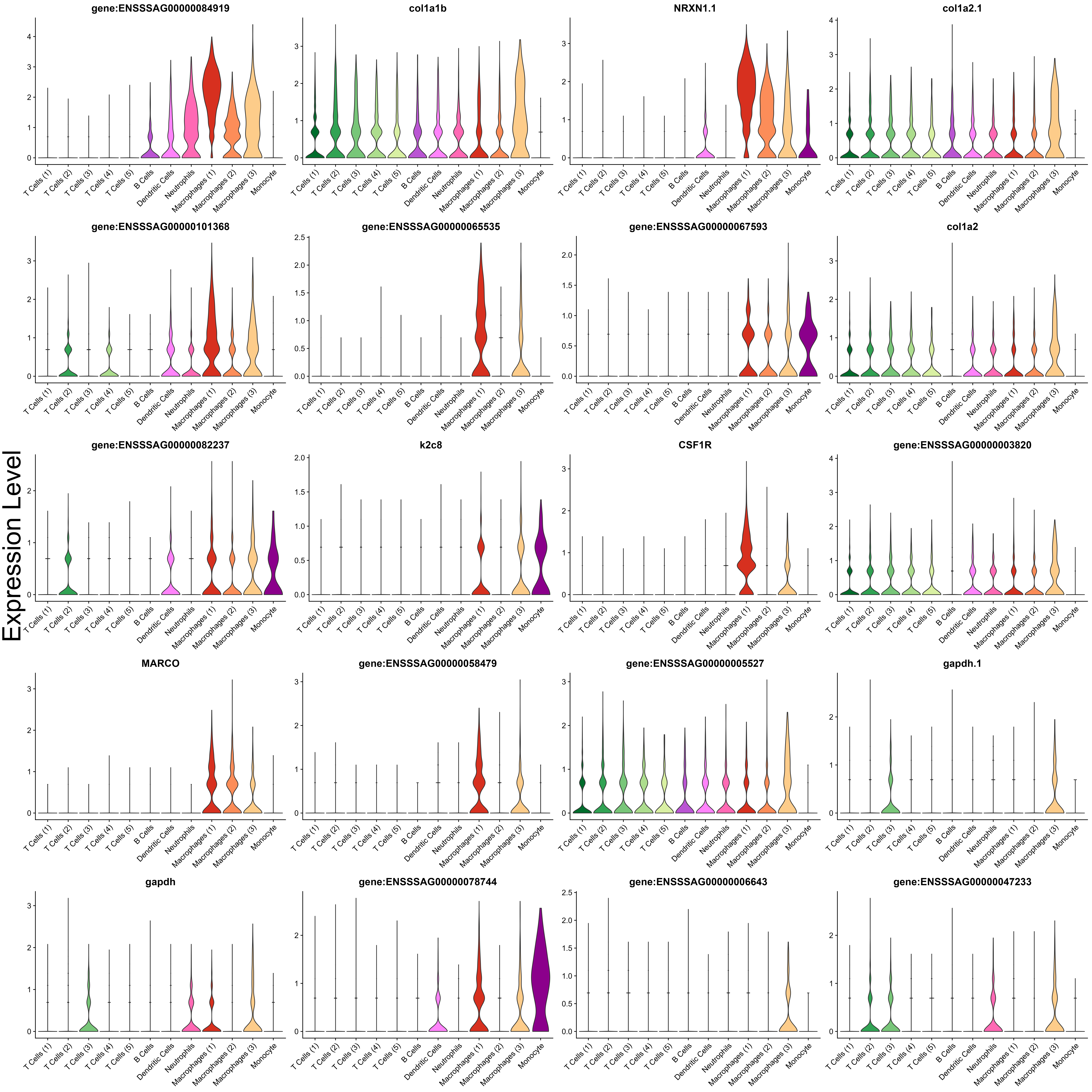

**Fig.S19** Violin plots of expression levels (based on the SCT assay) for the top 20 significant (adjusted p-value < 0.05) marker genes for the macrophages (3) cluster (based on log-scale two-fold change in expression) of the Atlantic salmon immune cells only data subset (for UMAP see Fig.3a).

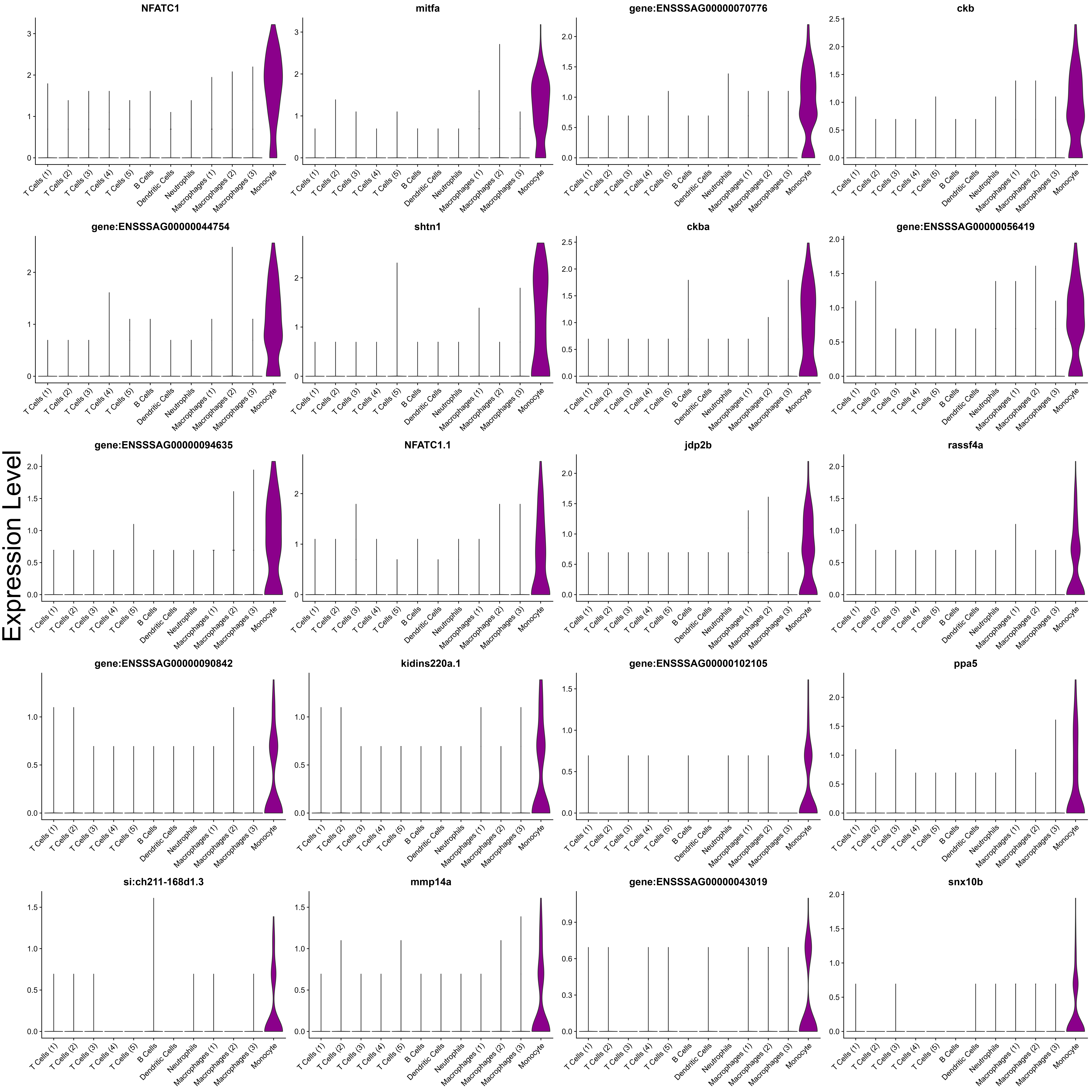

**Fig.S20** Violin plots of expression levels (based on the SCT assay) for the top 20 significant (adjusted p-value < 0.05) marker genes for monocytes (based on log-scale two-fold change in expression) of the Atlantic salmon immune cells only data subset (for UMAP see Fig.3a).

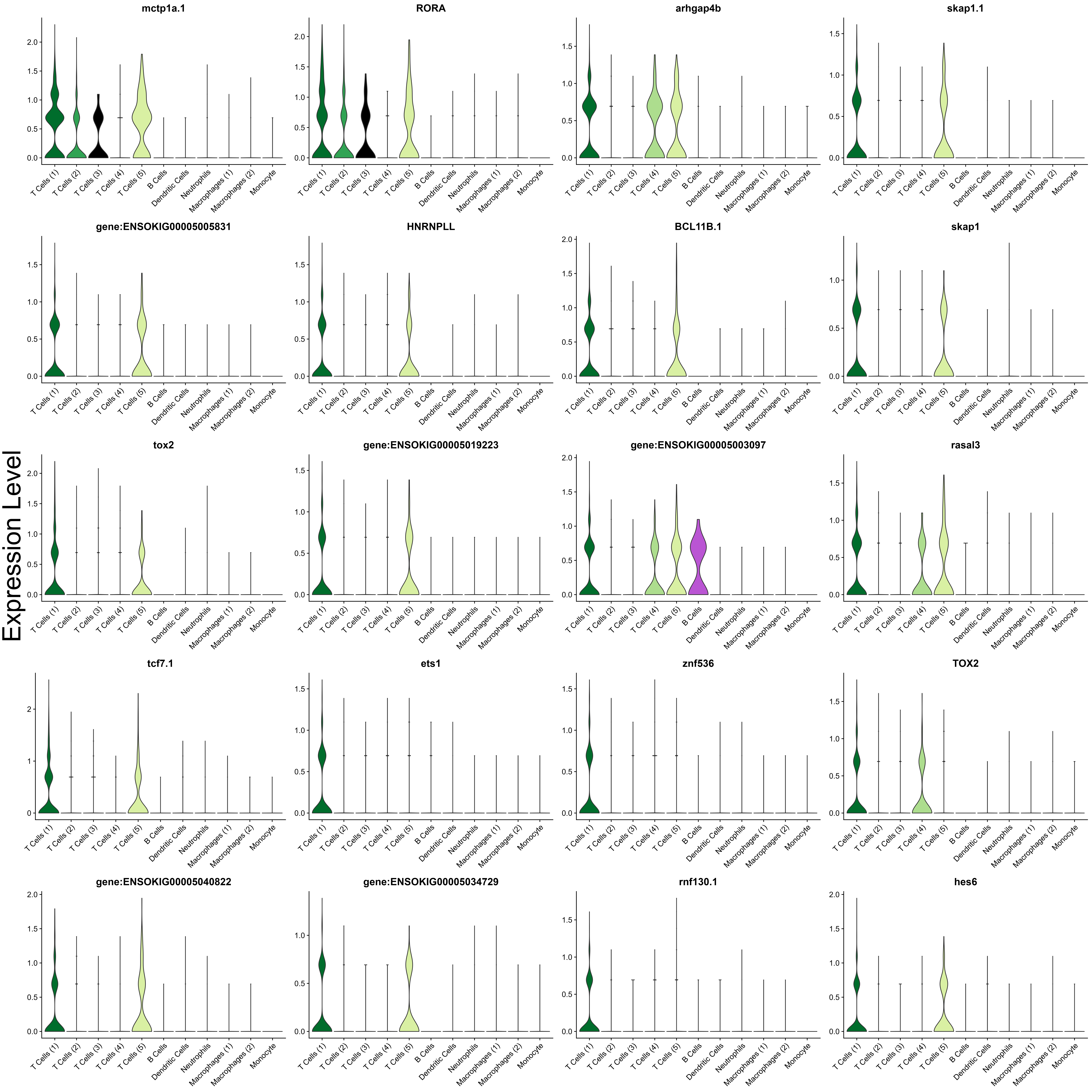

**Fig.S21** Violin plots of expression levels (based on the SCT assay) for the top 20 significant (adjusted p-value < 0.05) marker genes for the T cells (1) cluster (based on log-scale two-fold change in expression) of the coho salmon immune cells only data subset (for UMAP see Fig.3d).

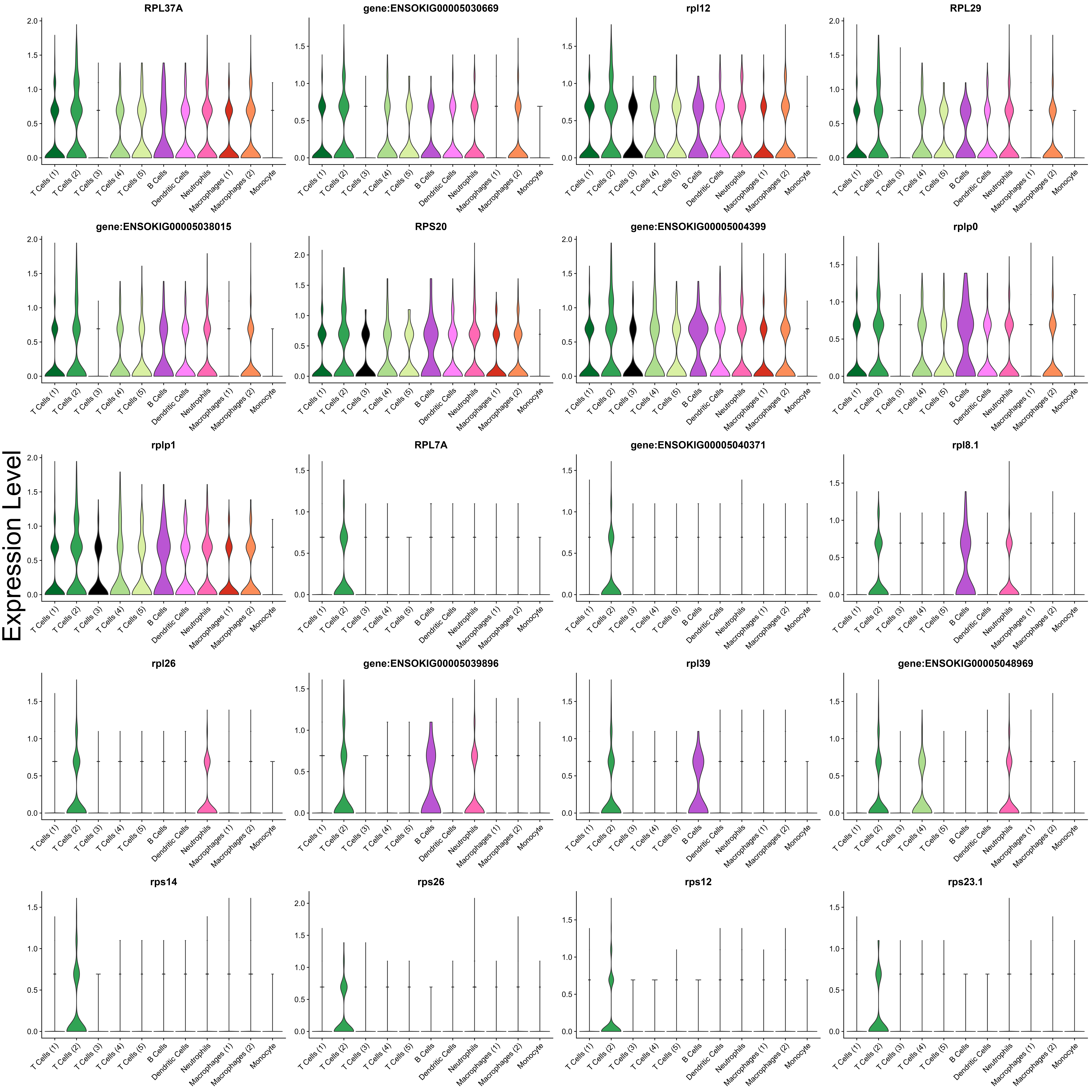

**Fig.S22** Violin plots of expression levels (based on the SCT assay) for the top 20 significant (adjusted p-value < 0.05) marker genes for the T cells (2) cluster (based on log-scale two-fold change in expression) of the coho salmon immune cells only data subset (for UMAP see Fig.3d).

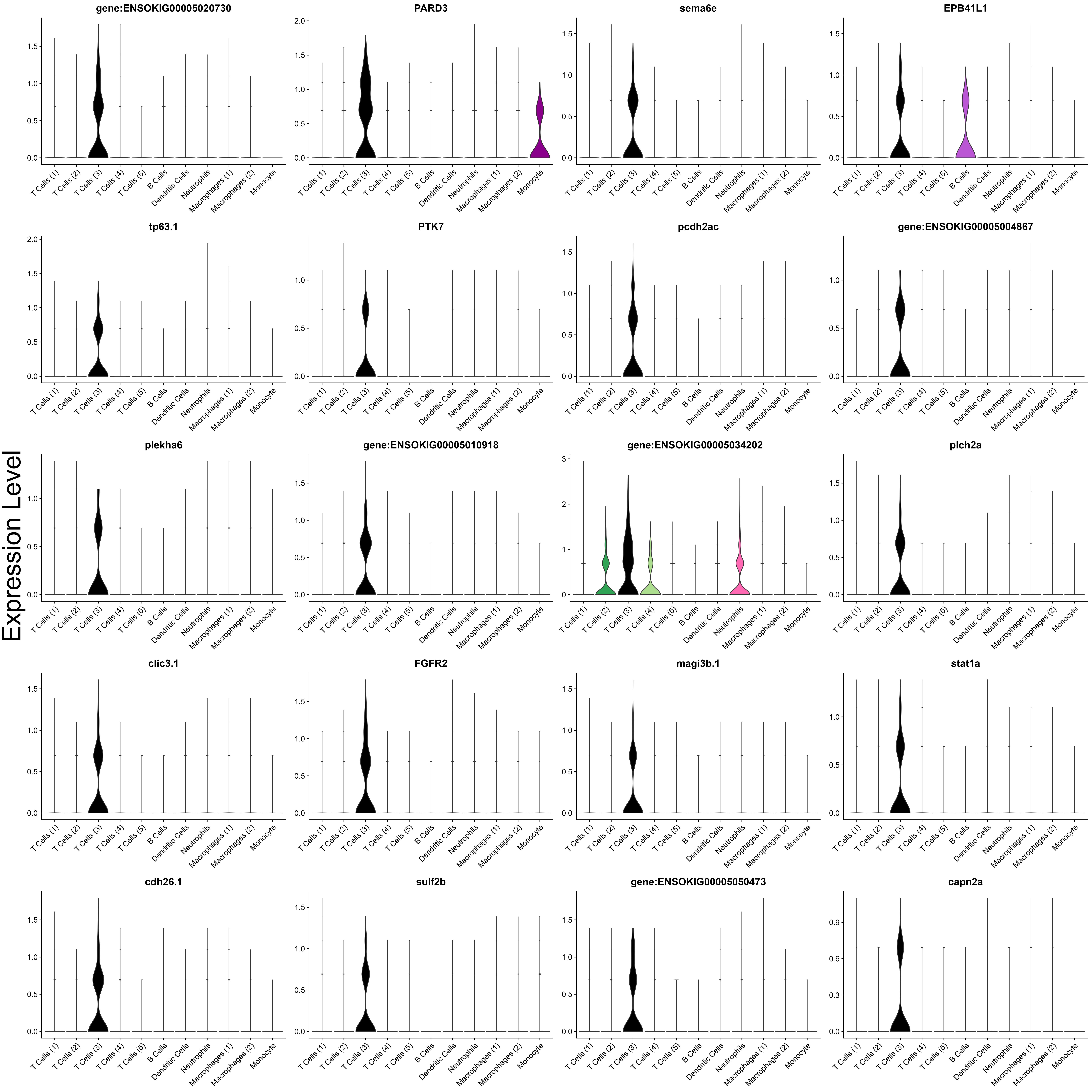

**Fig.S23** Violin plots of expression levels (based on the SCT assay) for the top 20 significant (adjusted p-value < 0.05) marker genes for the T cells (3) cluster (based on log-scale two-fold change in expression) of the coho salmon immune cells only data subset (for UMAP see Fig.3d).

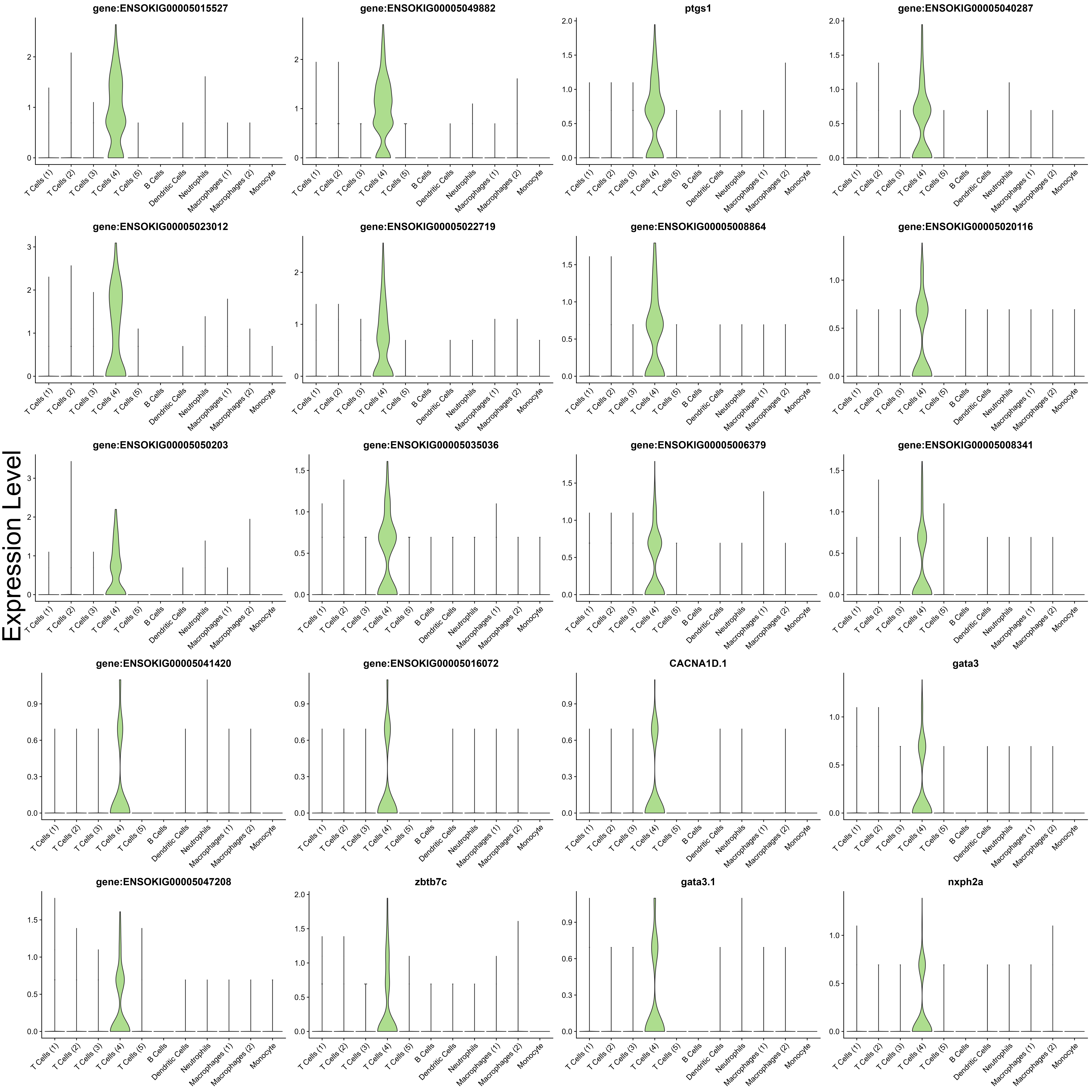

**Fig.S24** Violin plots of expression levels (based on the SCT assay) for the top 20 significant (adjusted p-value < 0.05) marker genes for the T cells (4) cluster (based on log-scale two-fold change in expression) of the coho salmon immune cells only data subset (for UMAP see Fig.3d).

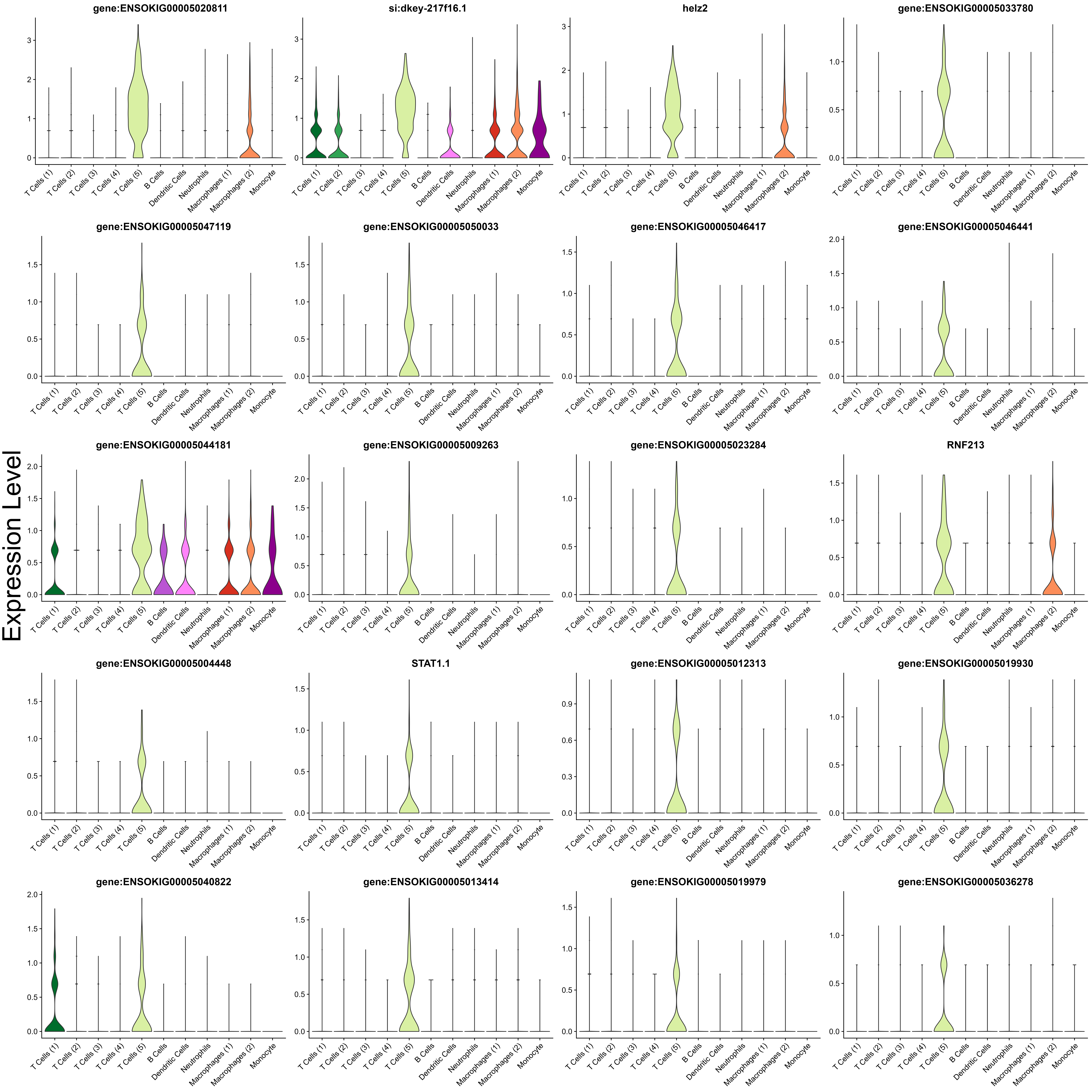

**Fig.S25** Violin plots of expression levels (based on the SCT assay) for the top 20 significant (adjusted p-value < 0.05) marker genes for the T cells (5) cluster (based on log-scale two-fold change in expression) of the coho salmon immune cells only data subset (for UMAP see Fig.3d).

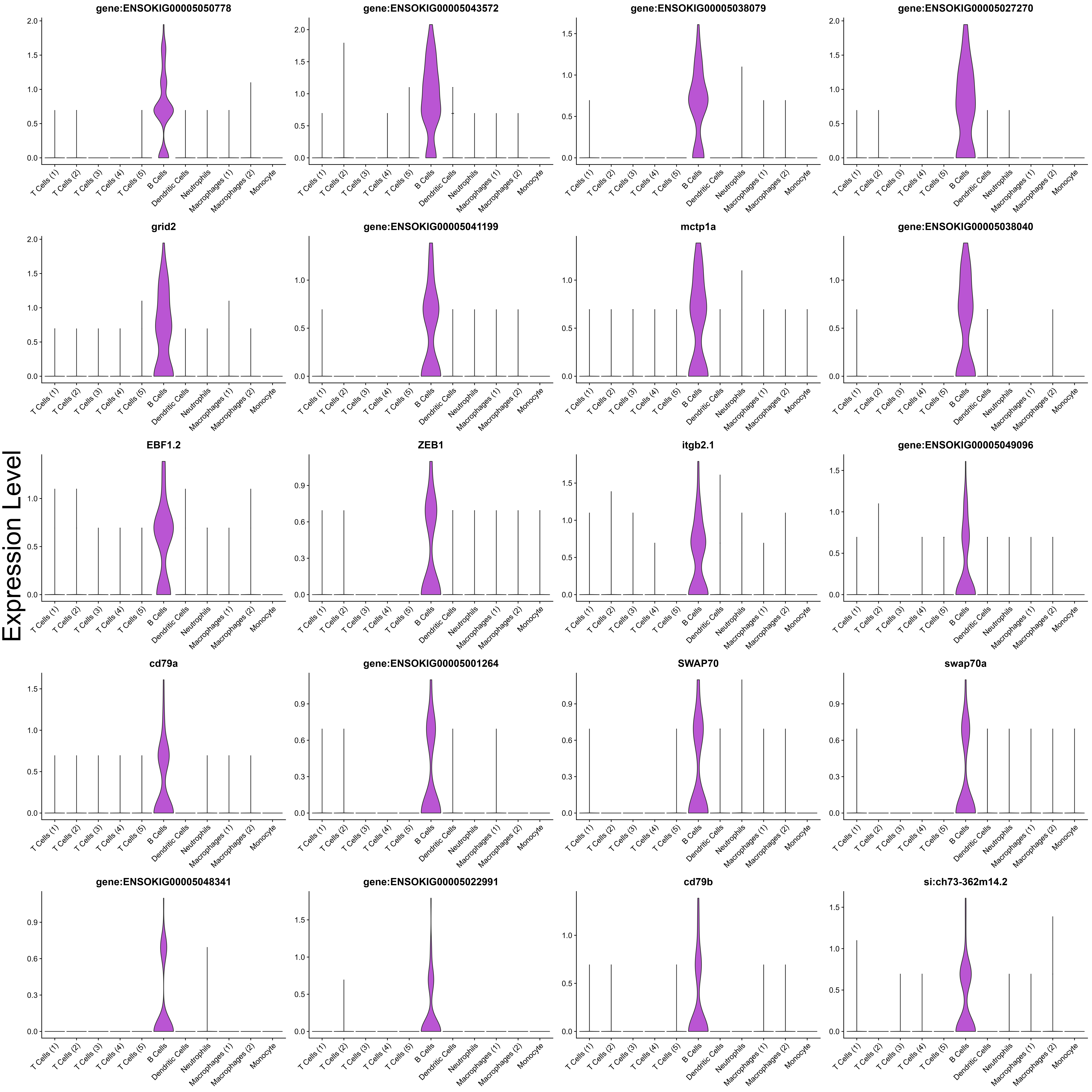

**Fig.S26** Violin plots of expression levels (based on the SCT assay) for the top 20 significant (adjusted p-value < 0.05) marker genes for B cells (based on log-scale two-fold change in expression) of the coho salmon immune cells only data subset (for UMAP see Fig.3d).

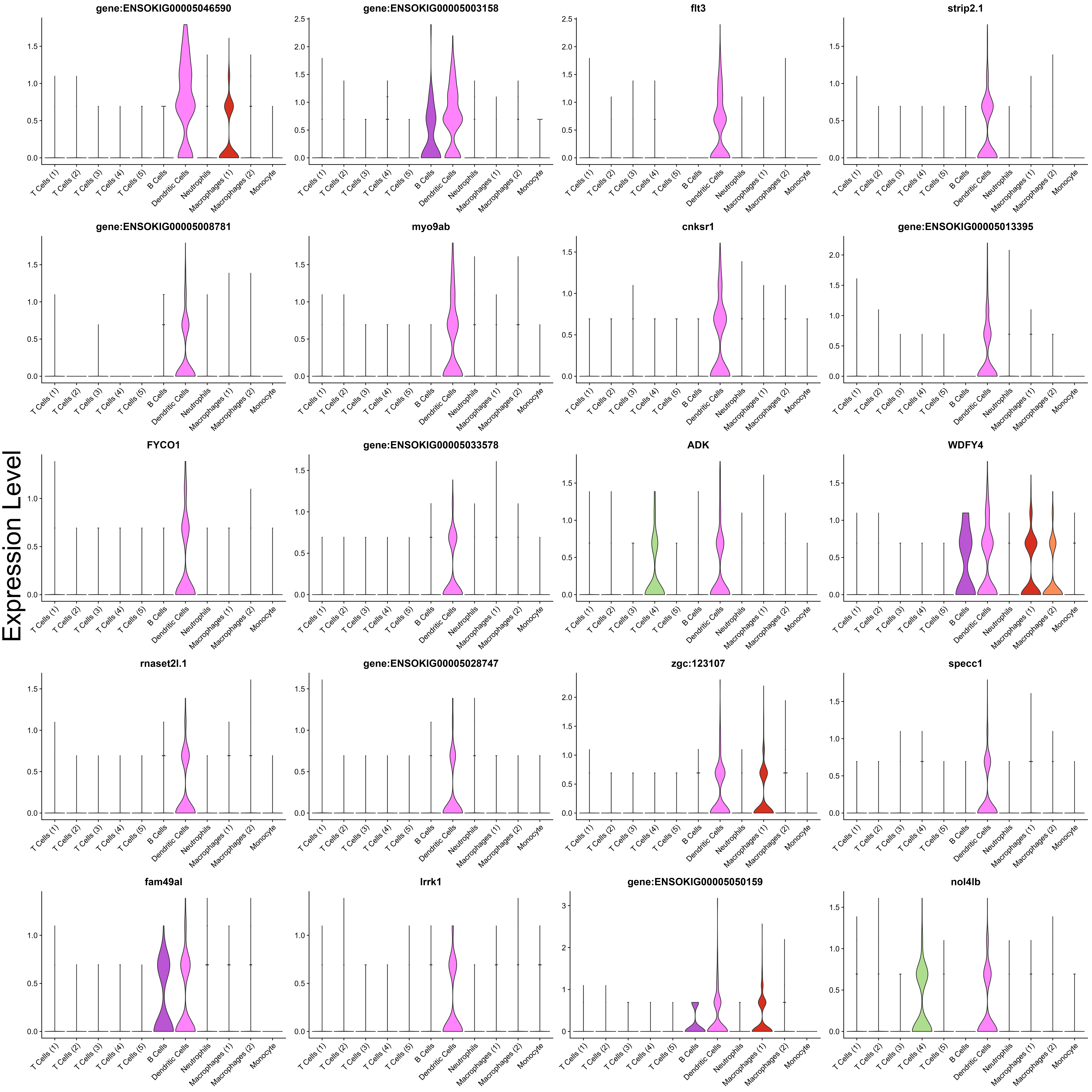

**Fig.S27** Violin plots of expression levels (based on the SCT assay) for the top 20 significant (adjusted p-value < 0.05) marker genes for dendritic cells (based on log-scale two-fold change in expression) of the coho salmon immune cells only data subset (for UMAP see Fig.3d).

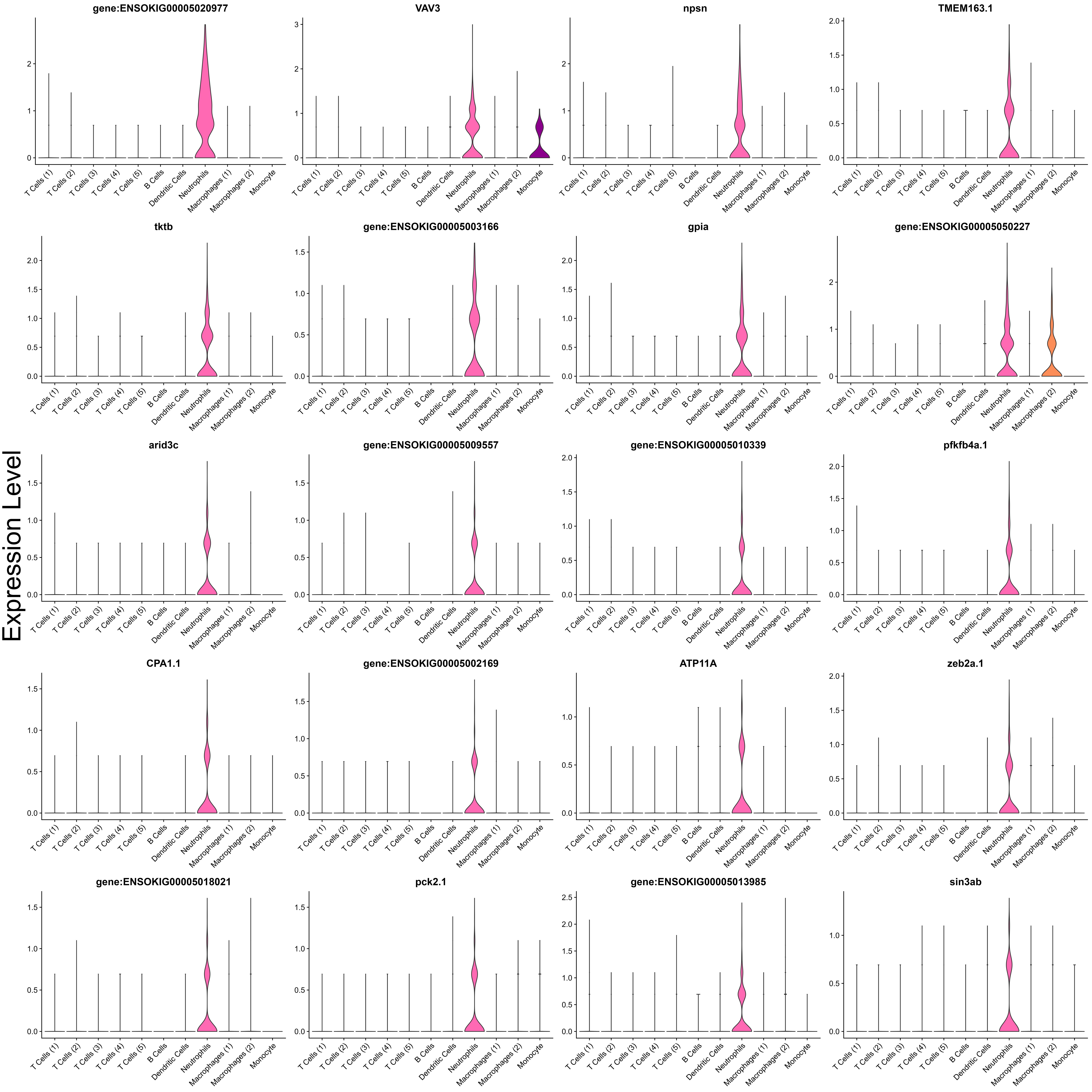

**Fig.S28** Violin plots of expression levels (based on the SCT assay) for the top 20 significant (adjusted p-value < 0.05) marker genes for neutrophils (based on log-scale two-fold change in expression) of the coho salmon immune cells only data subset (for UMAP see Fig.3d).

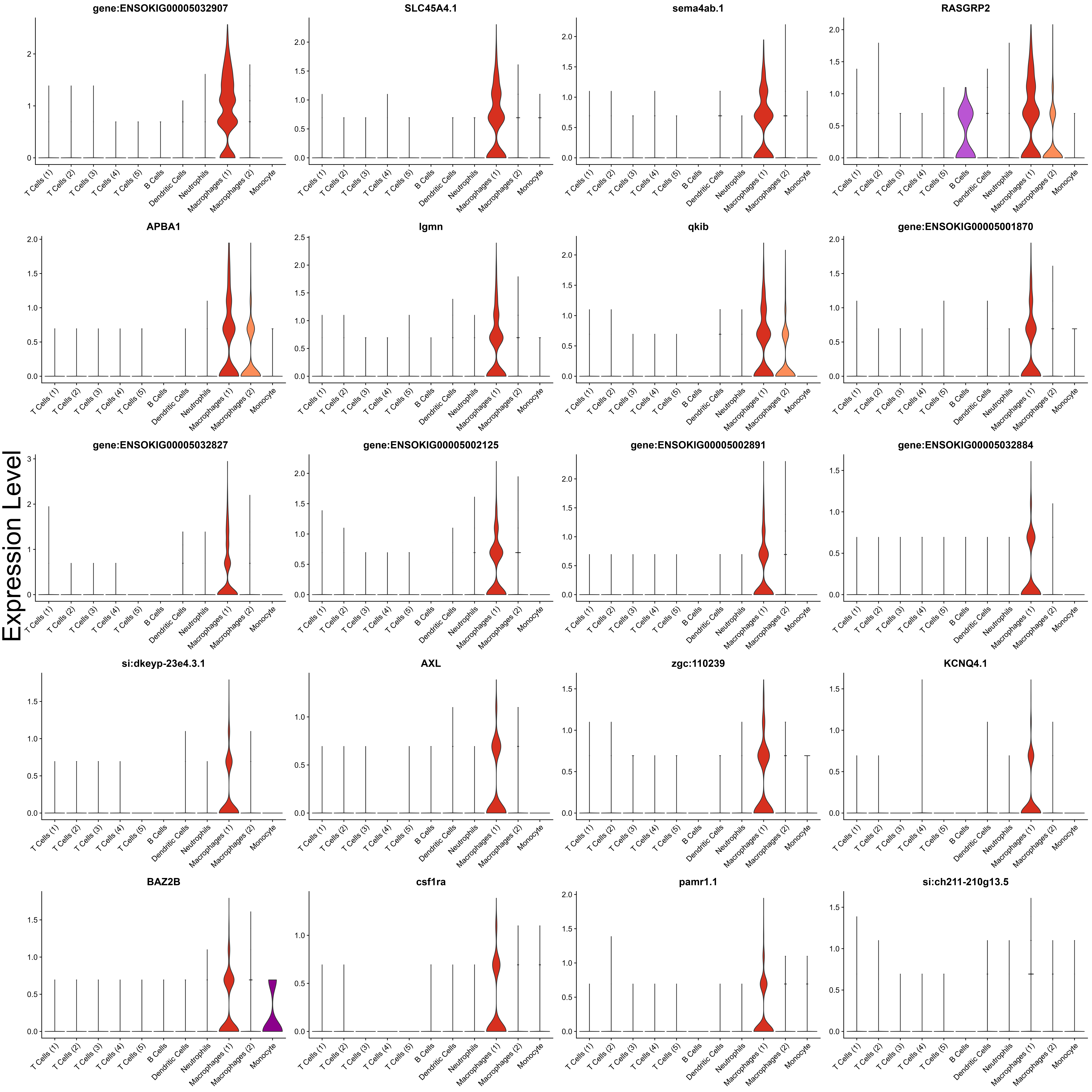

**Fig.S29** Violin plots of expression levels (based on the SCT assay) for the top 20 significant (adjusted p-value < 0.05) marker genes for the macrophages (1) cluster (based on log-scale two-fold change in expression) of the coho salmon immune cells only data subset (for UMAP see Fig.3d).

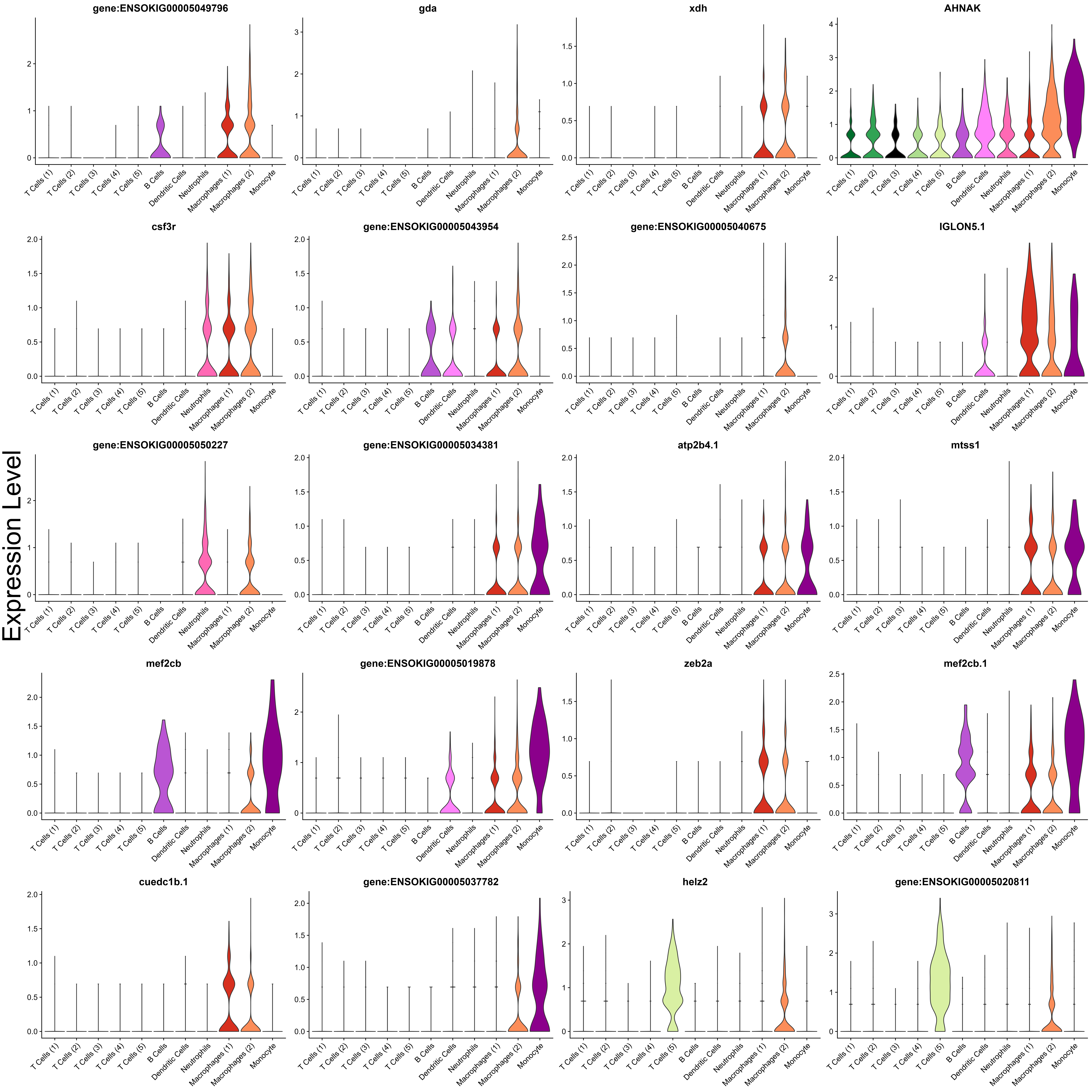

**Fig.S30** Violin plots of expression levels (based on the SCT assay) for the top 20 significant (adjusted p-value < 0.05) marker genes for the macrophages (2) cluster (based on log-scale two-fold change in expression) of the coho salmon immune cells only data subset (for UMAP see Fig.3d).

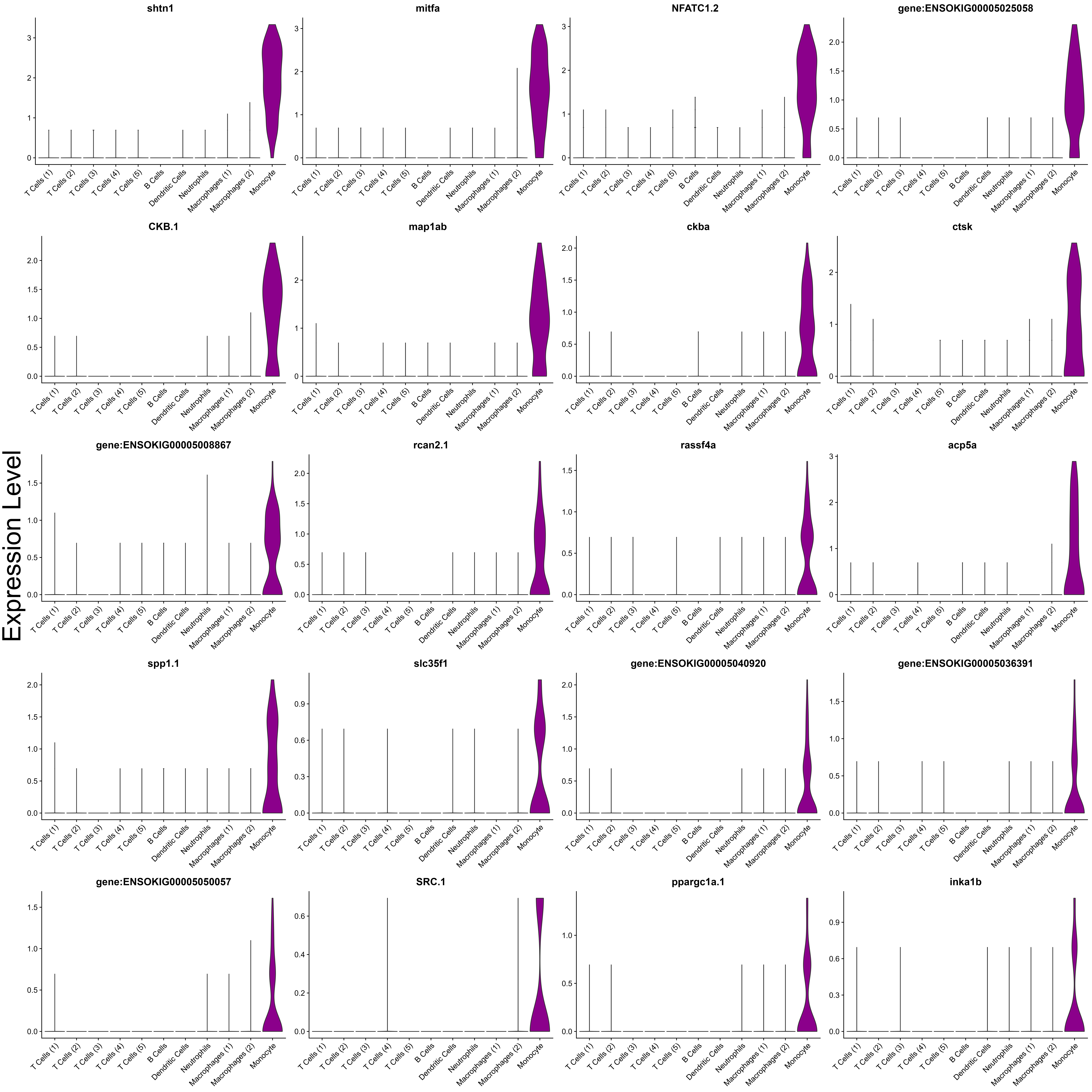

**Fig.S31** Violin plots of expression levels (based on the SCT assay) for the top 20 significant (adjusted p-value < 0.05) marker genes for monocytes (based on log-scale two-fold change in expression) of the coho salmon immune cells only data subset (for UMAP see Fig.3d).

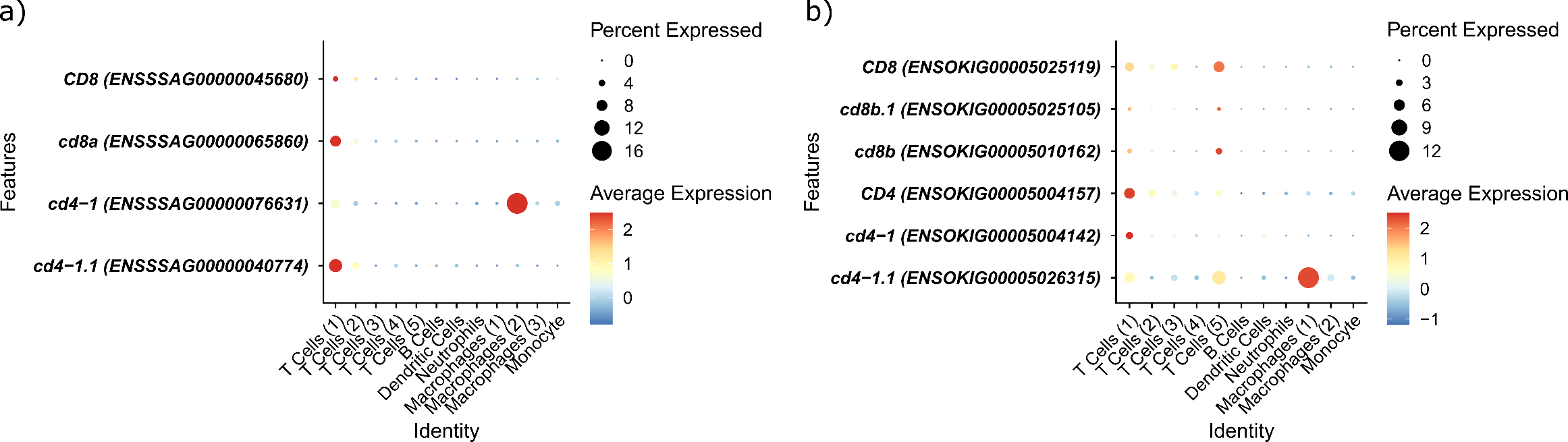

**Fig.S32** Expression of CD4 and CD8 paralogs in the immune cell subclusters identified within a) Atlantic salmon and b) coho salmon.

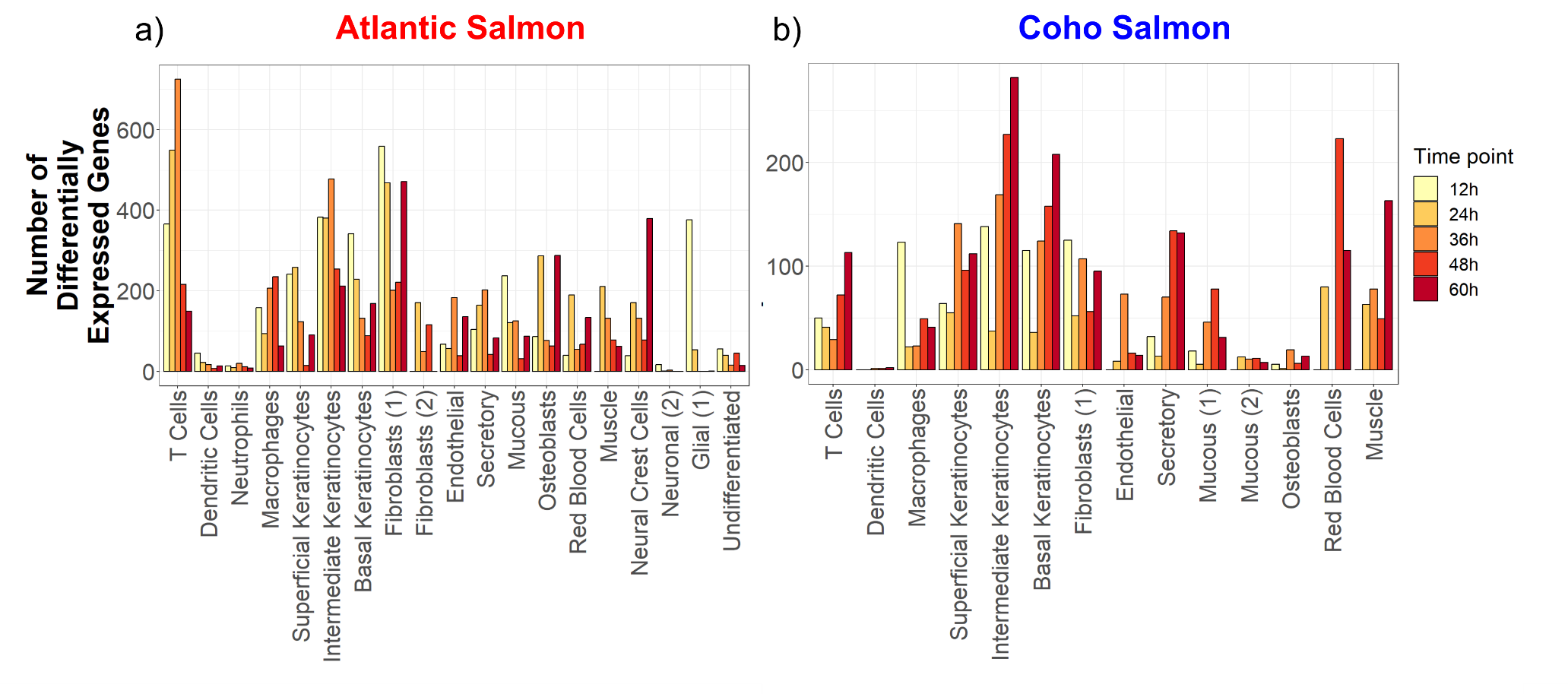

Fig.S33 Number of genes detected as differentially expressed between the control samples and each of the infected time points (12h, 24h, 36h, 48h, 60h) for each cell type for a) Atlantic salmon and b) coho salmon.

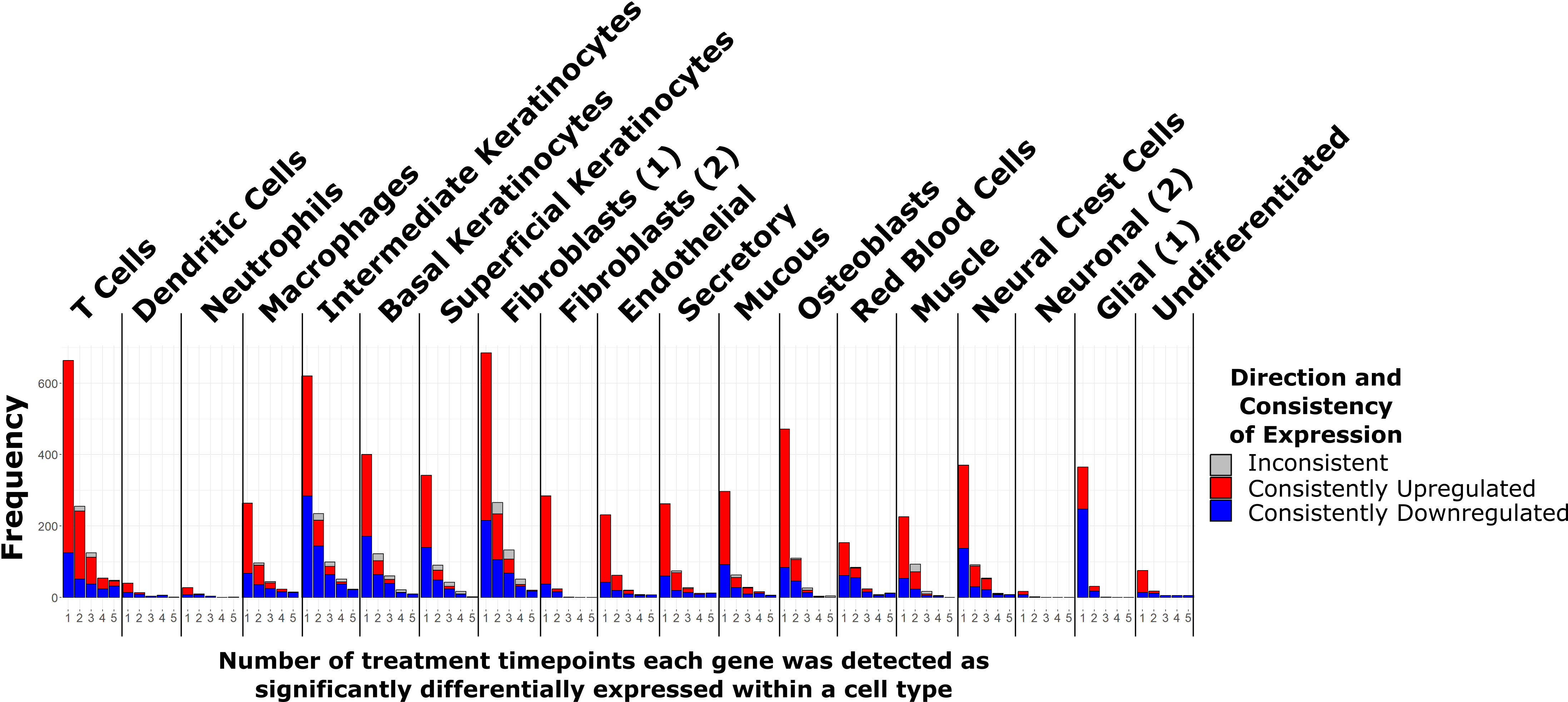

**Fig.S34** Number of times each gene was detected as differentially expressed (1-5 time points) between any of the treatment time points and the control sample for a given cell type detected within the Atlantic salmon samples. Counts are coloured by whether differentially expressed genes were either consistently upregulated (red), downregulated (blue), or were detected within the same cell type as being significantly upregulated and downregulated at different time points (grey).

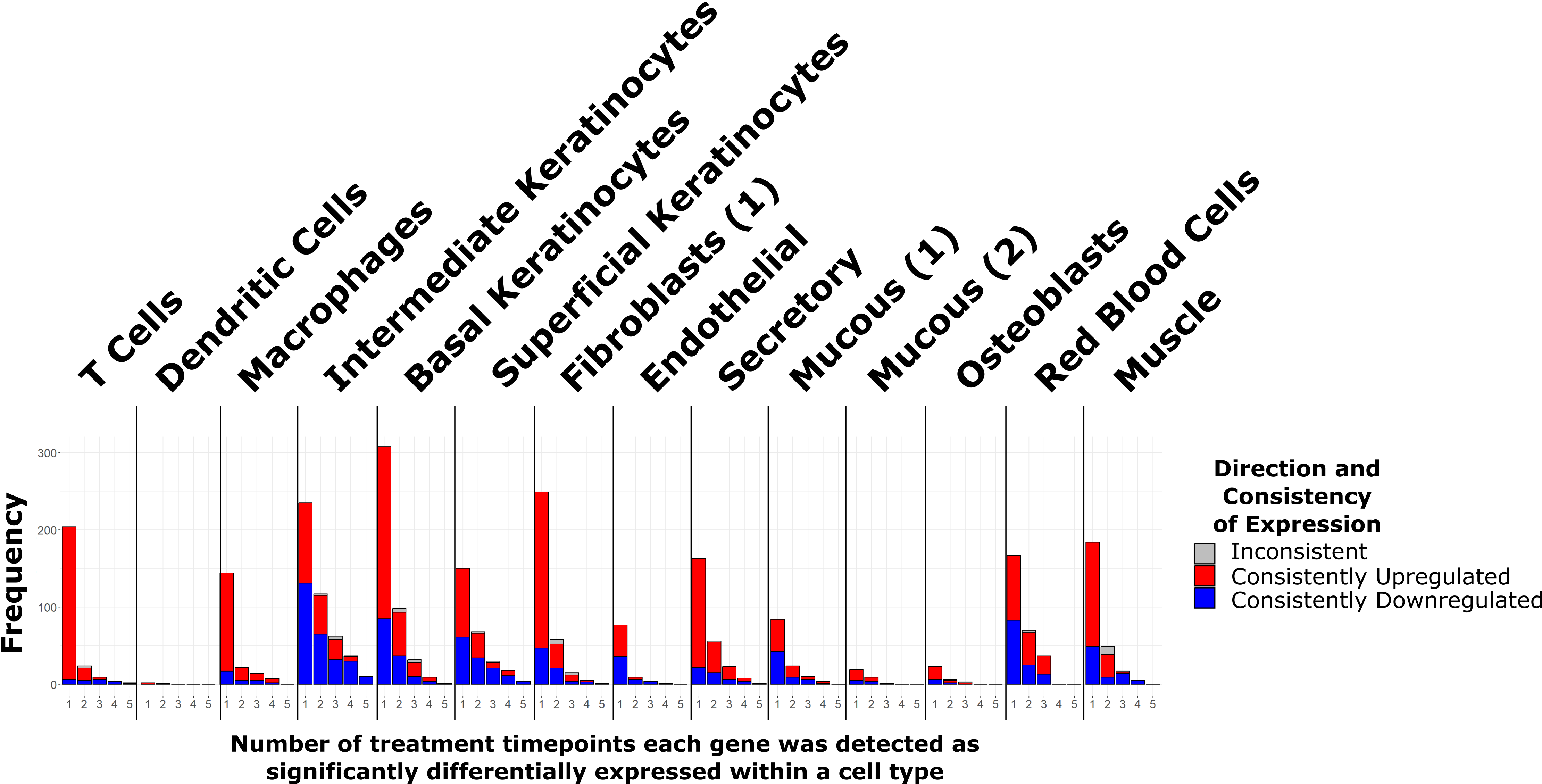

**Fig.S35** Number of times each gene was detected as differentially expressed (1-5 time points) between any of the treatment time points and the control sample for a given cell type detected within the coho salmon samples. Counts are coloured by whether differentially expressed genes were either consistently upregulated (red), downregulated (blue), or were detected within the same cell type as being significantly upregulated and downregulated at different time points (grey)

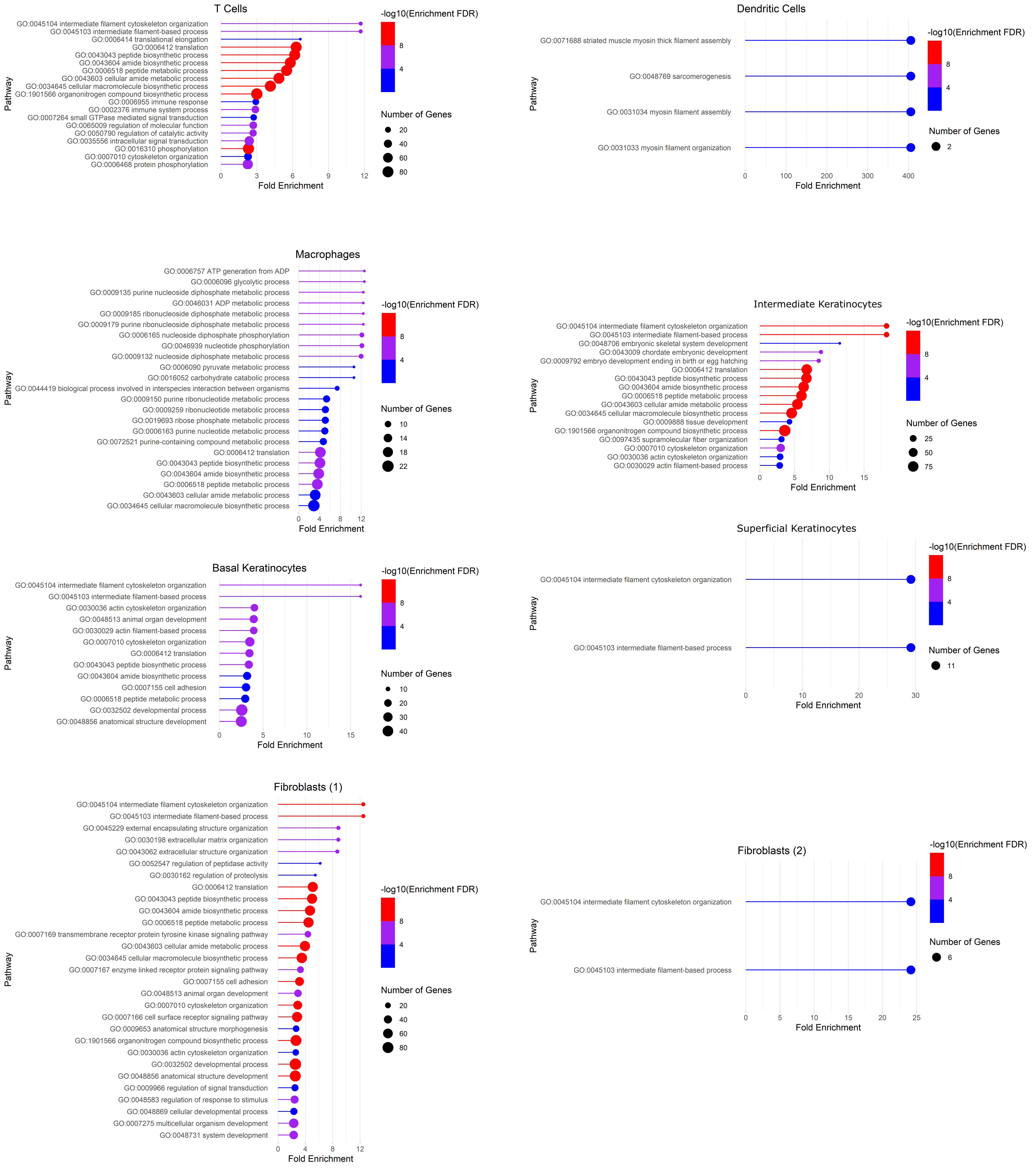

**Fig.S36** Significantly (FDR-adjusted p-value < 0.001) enriched Biological GO terms for each Atlantic salmon cell type based on the significantly differentially expressed genes detected between the control samples and any of the five treatment time points (12h, 24h, 36h, 48h, 60h). Please note that no significant GO terms were detected for Endothelial, Glial (1), Mucous, Neuronal (2), and Neutrophils. (Continued on next page.)

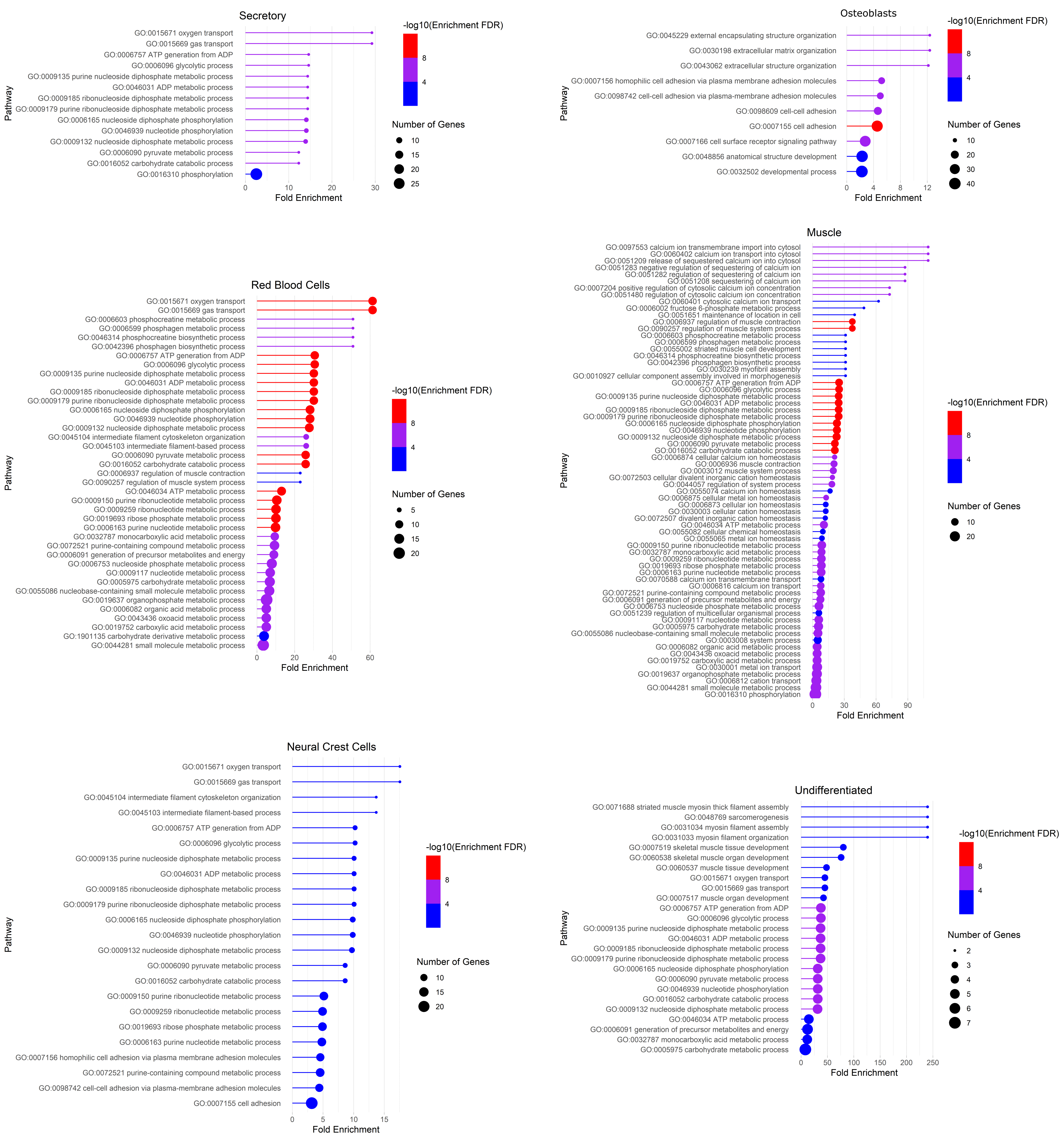

**Fig.S36** Continued.

**Fig.S37** Significantly (FDR-adjusted p-value < 0.001) enriched Biological GO terms for each coho salmon cell type based on the significantly differentially expressed genes detected between the control samples and any of the five treatment time points (12h, 24h, 36h, 48h, 60h). Please note that no significant GO terms were detected for Mucous (1), Mucous (2), and Dendritic Cells.

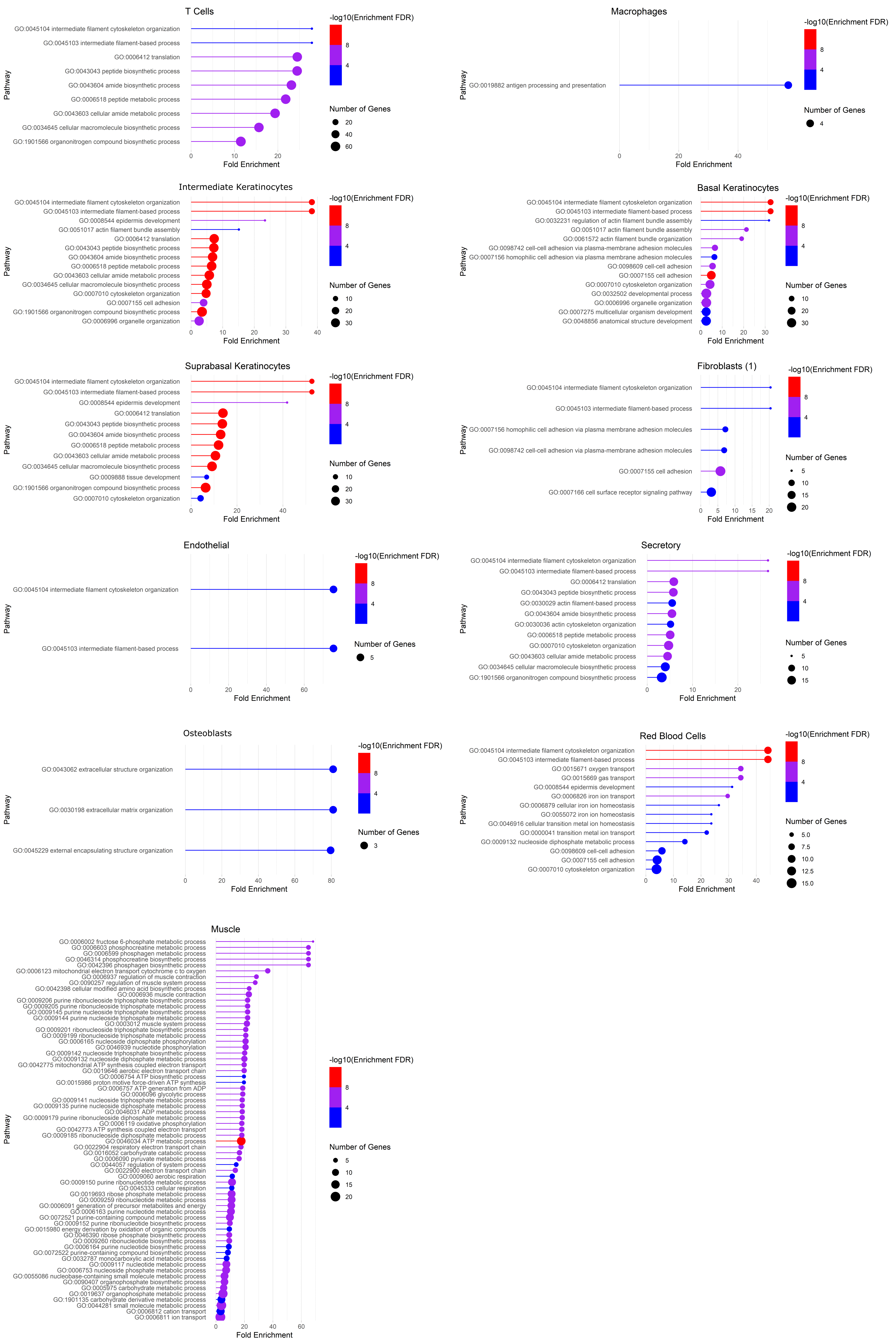

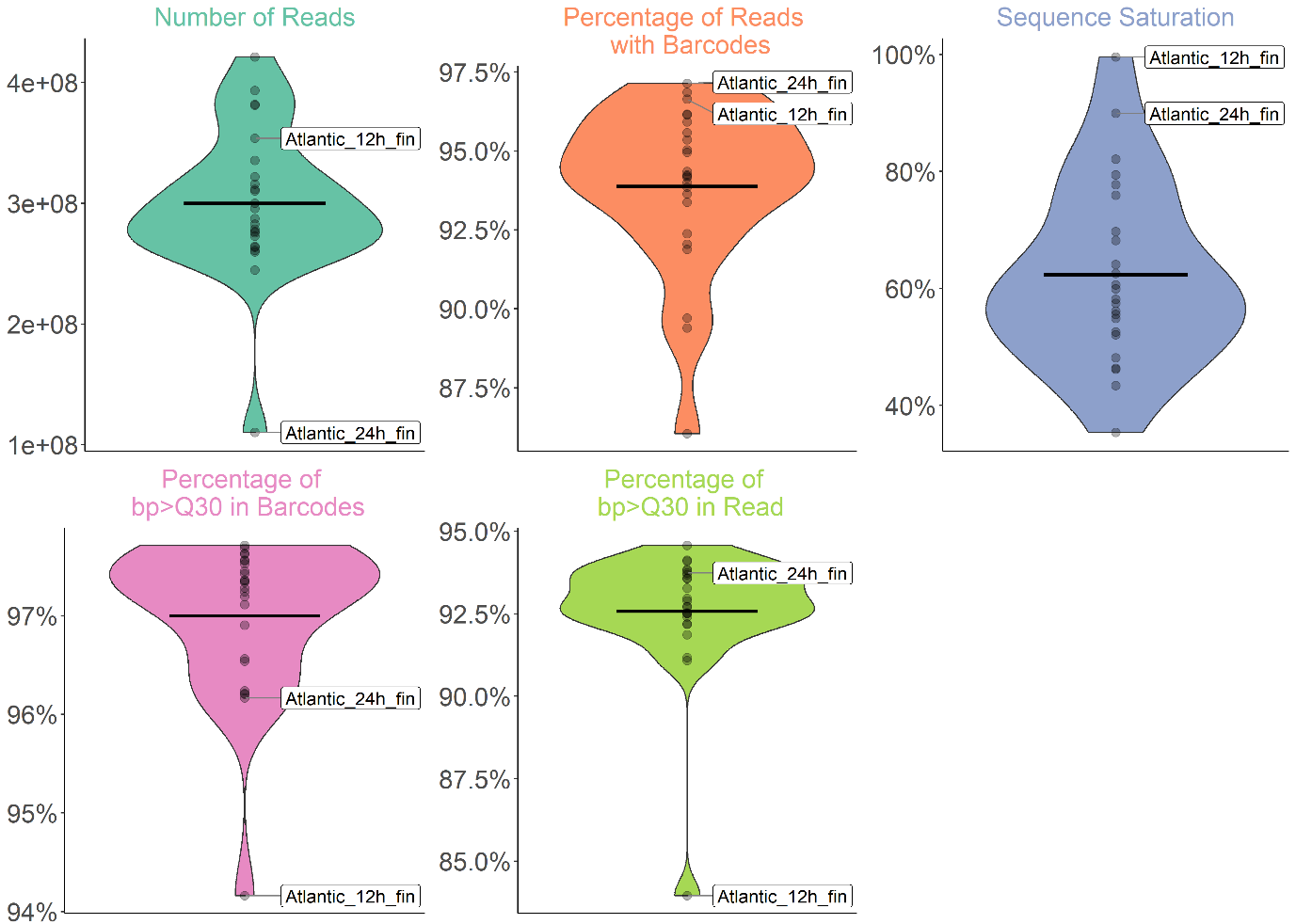

**Fig.S38** Summary statistics for all Atlantic and coho salmon libraries. Samples which were removed in downstream quality control filtering in Seurat are highlighted (i.e., Atlantic_12h_fin, Atlantic_24_fin).

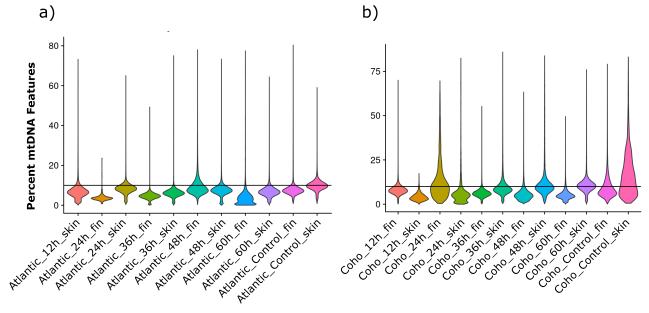

**Fig.S39** Percent of UMIs identified as mtDNA features for a) Atlantic salmon and b) coho salmon samples, with the 10% maximum threshold used for subsequent filtering indicated by a horizontal black line.

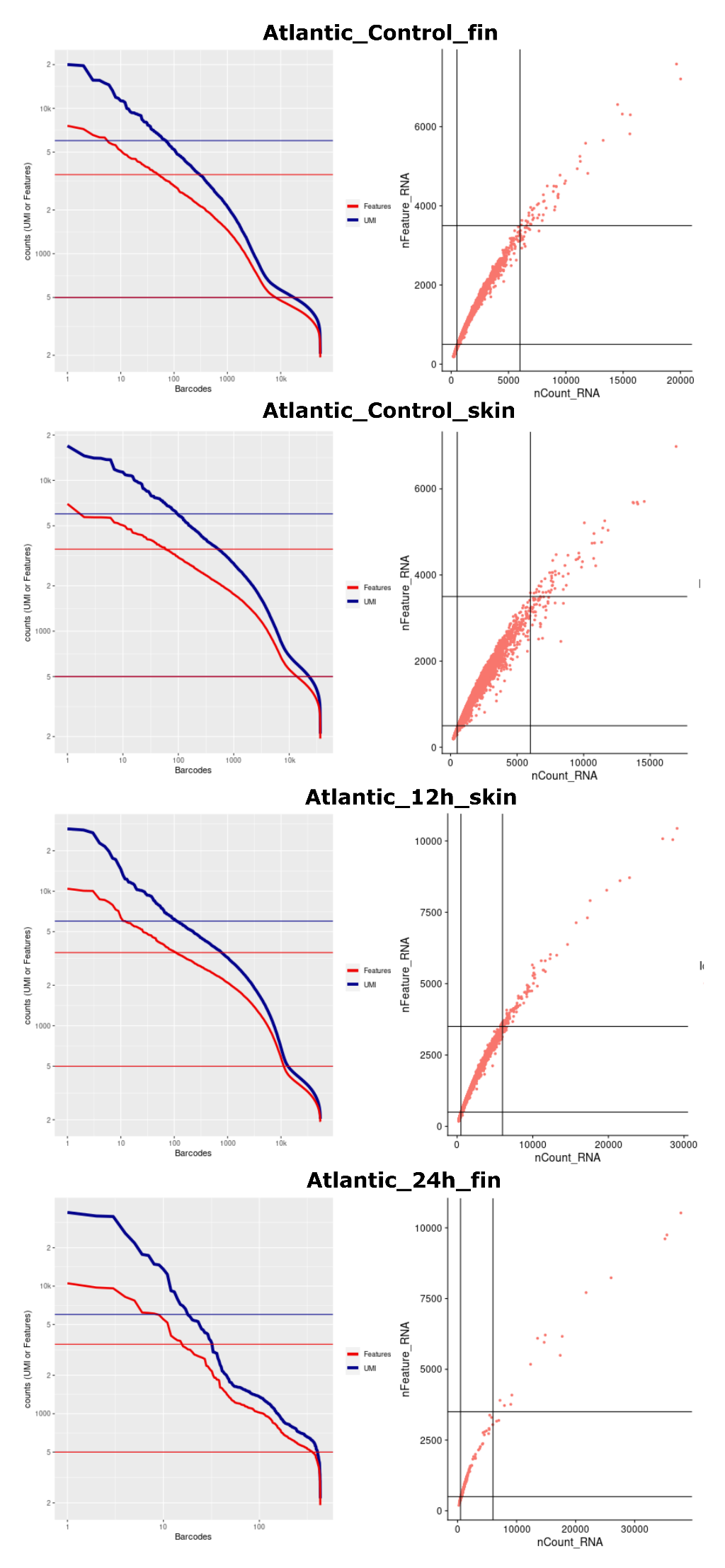

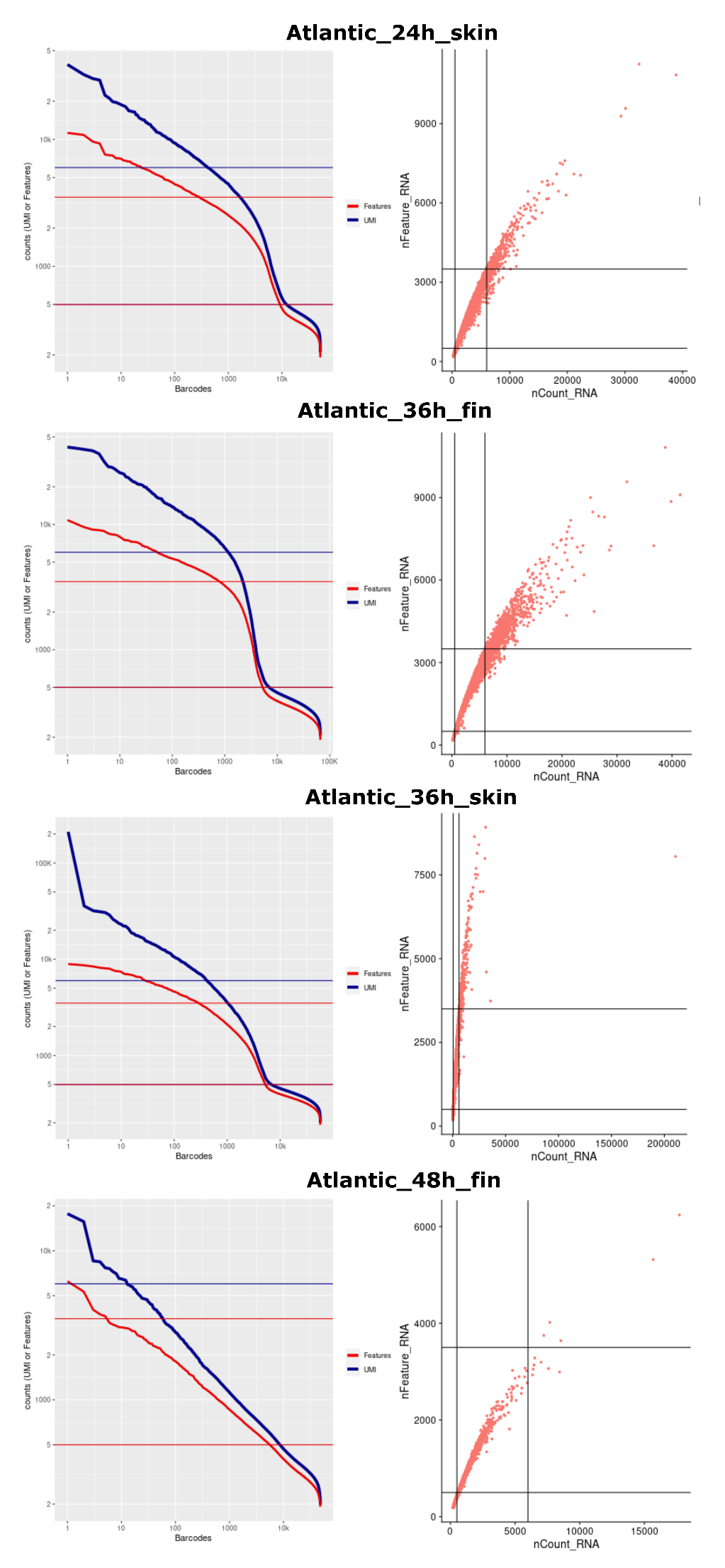

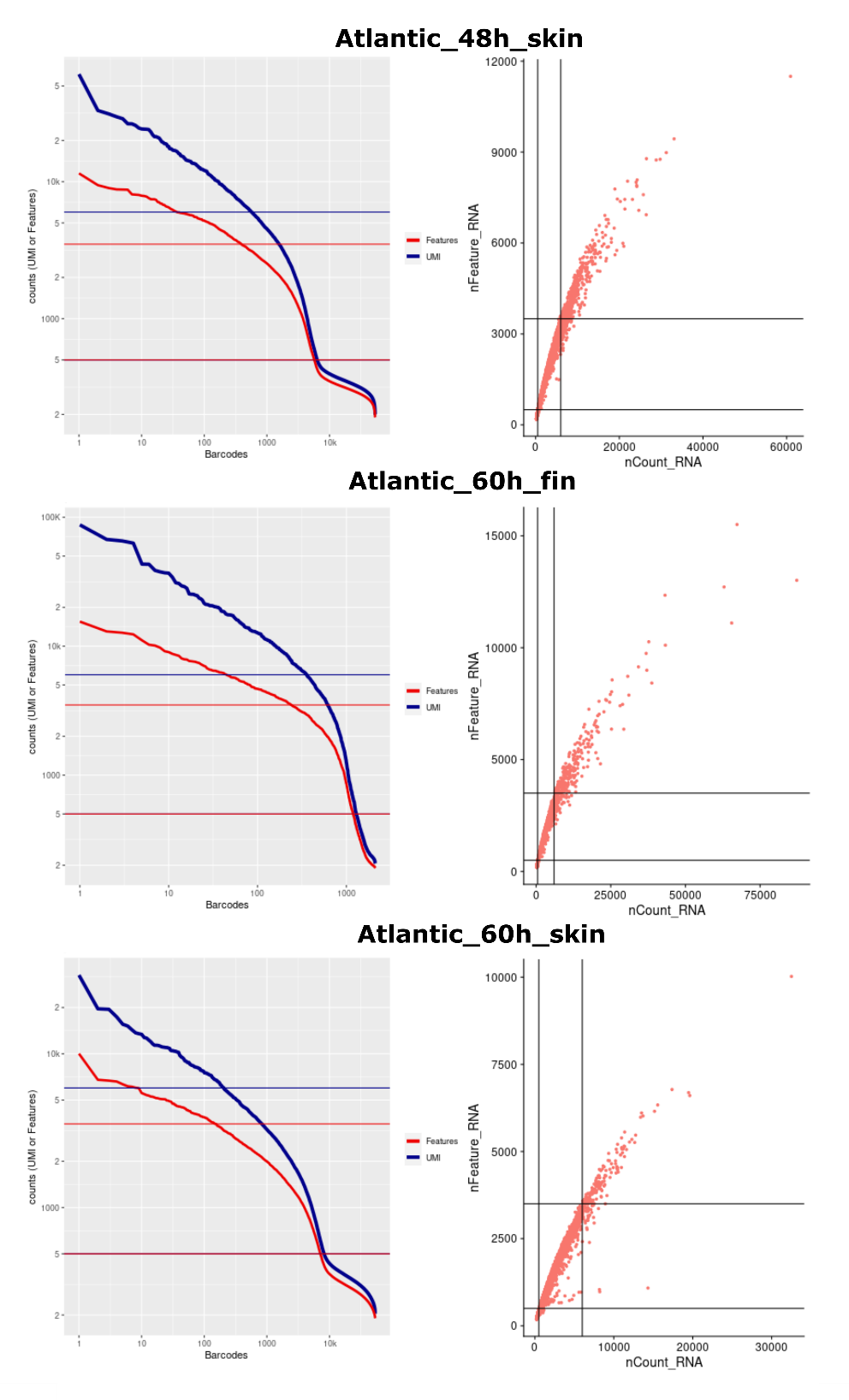

**Fig.S40** UMI and feature counts per cell barcode and feature counts vs. UMI counts for each Atlantic salmon sample.

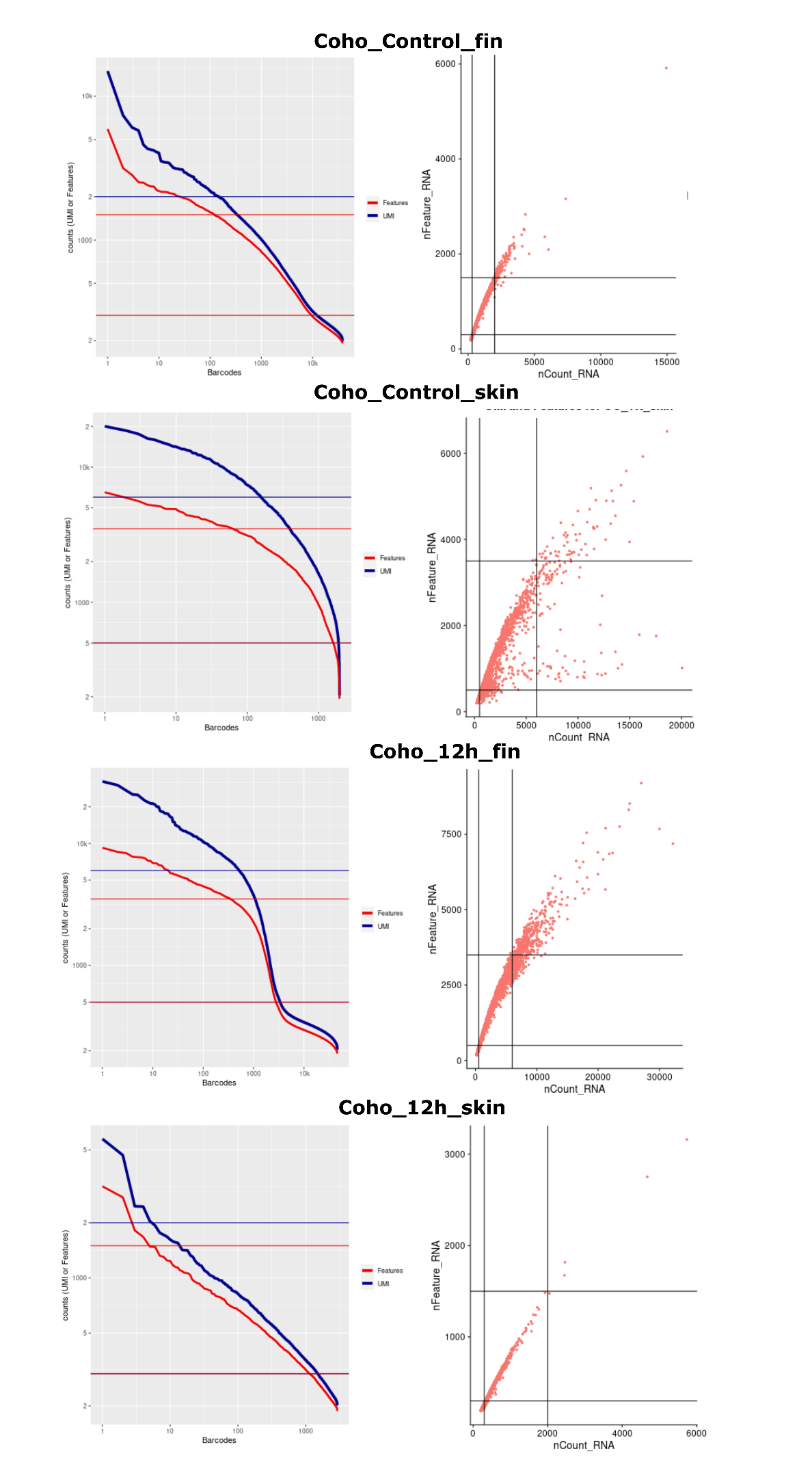

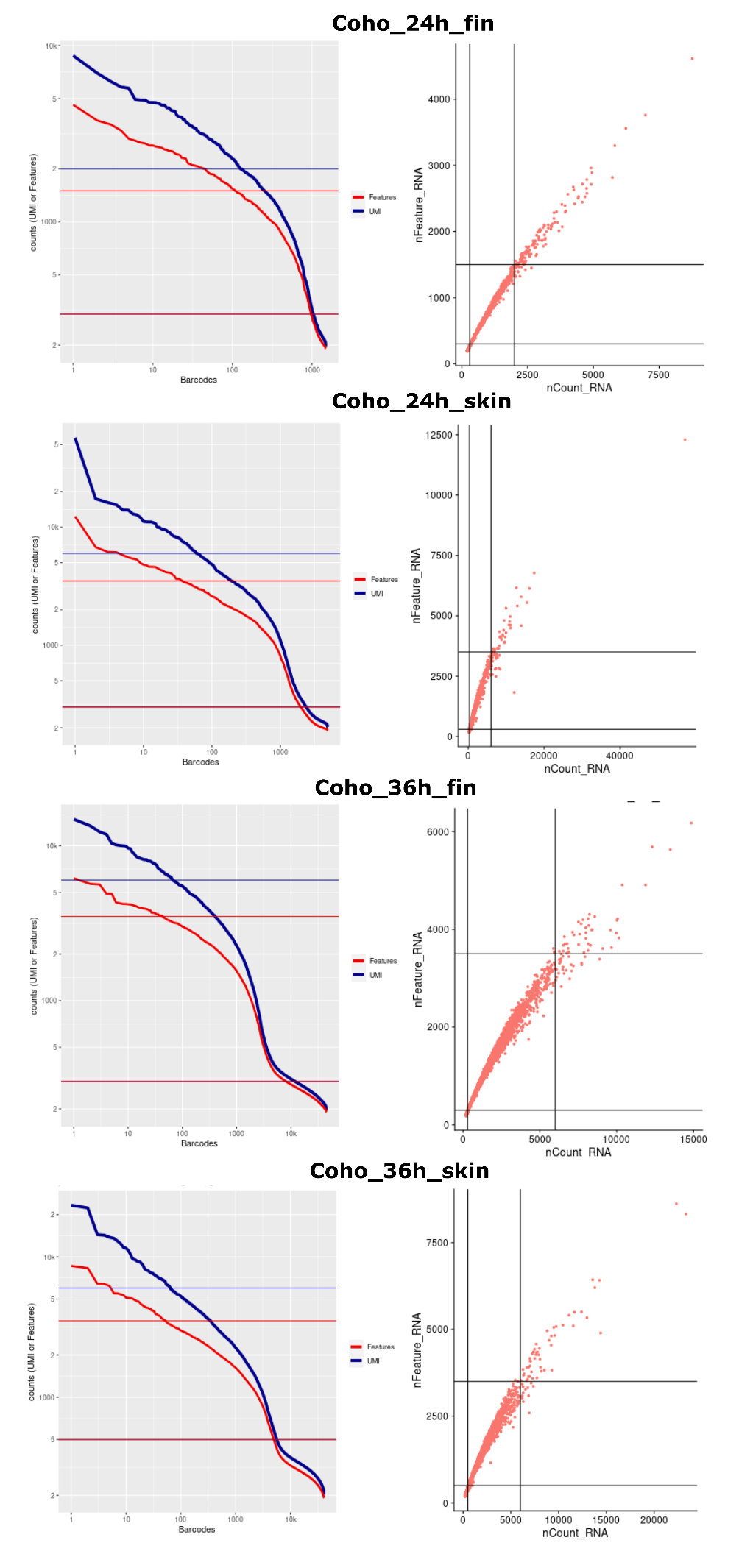

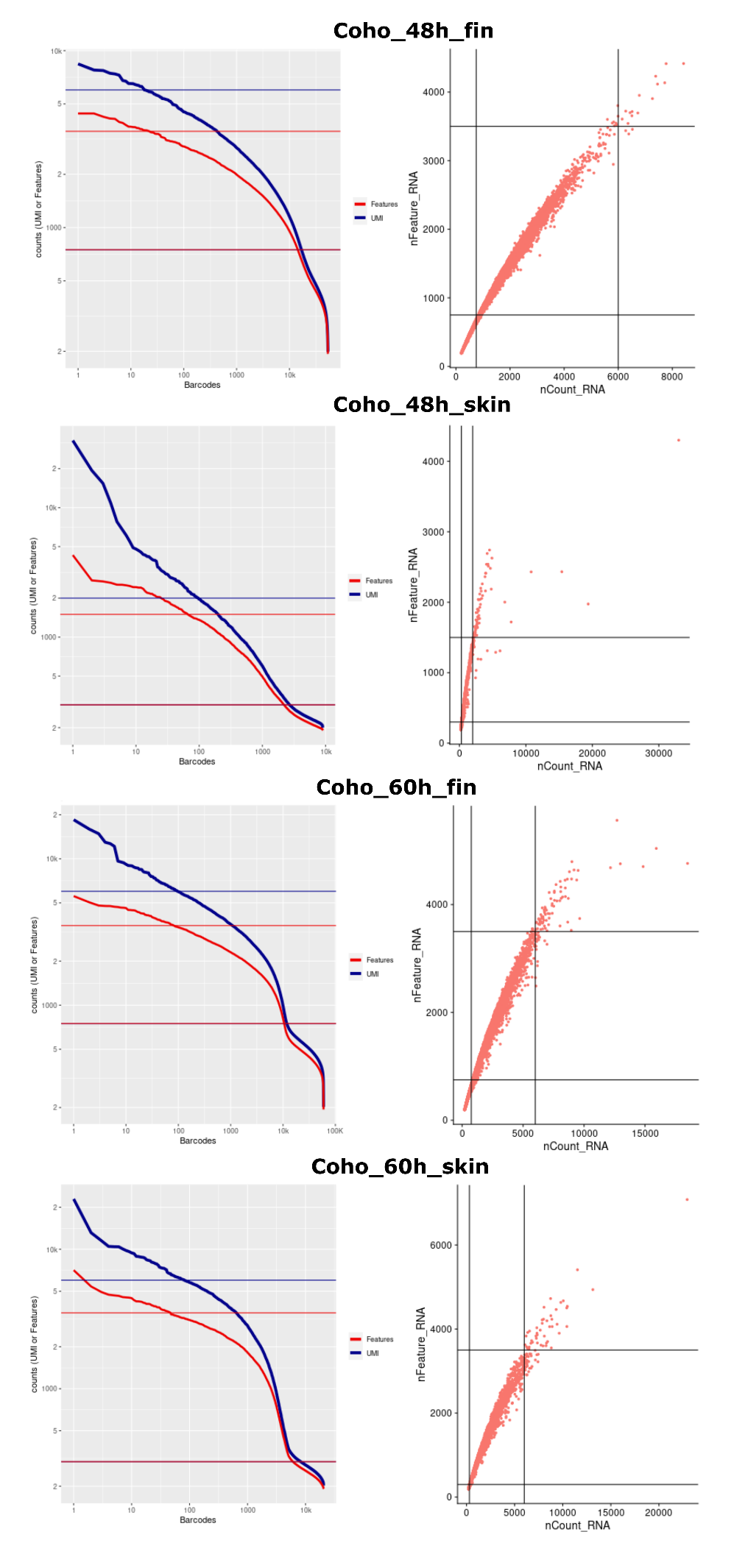

**Fig.S41** UMI and feature counts per cell barcode and feature counts vs. UMI counts for each coho salmon sample.

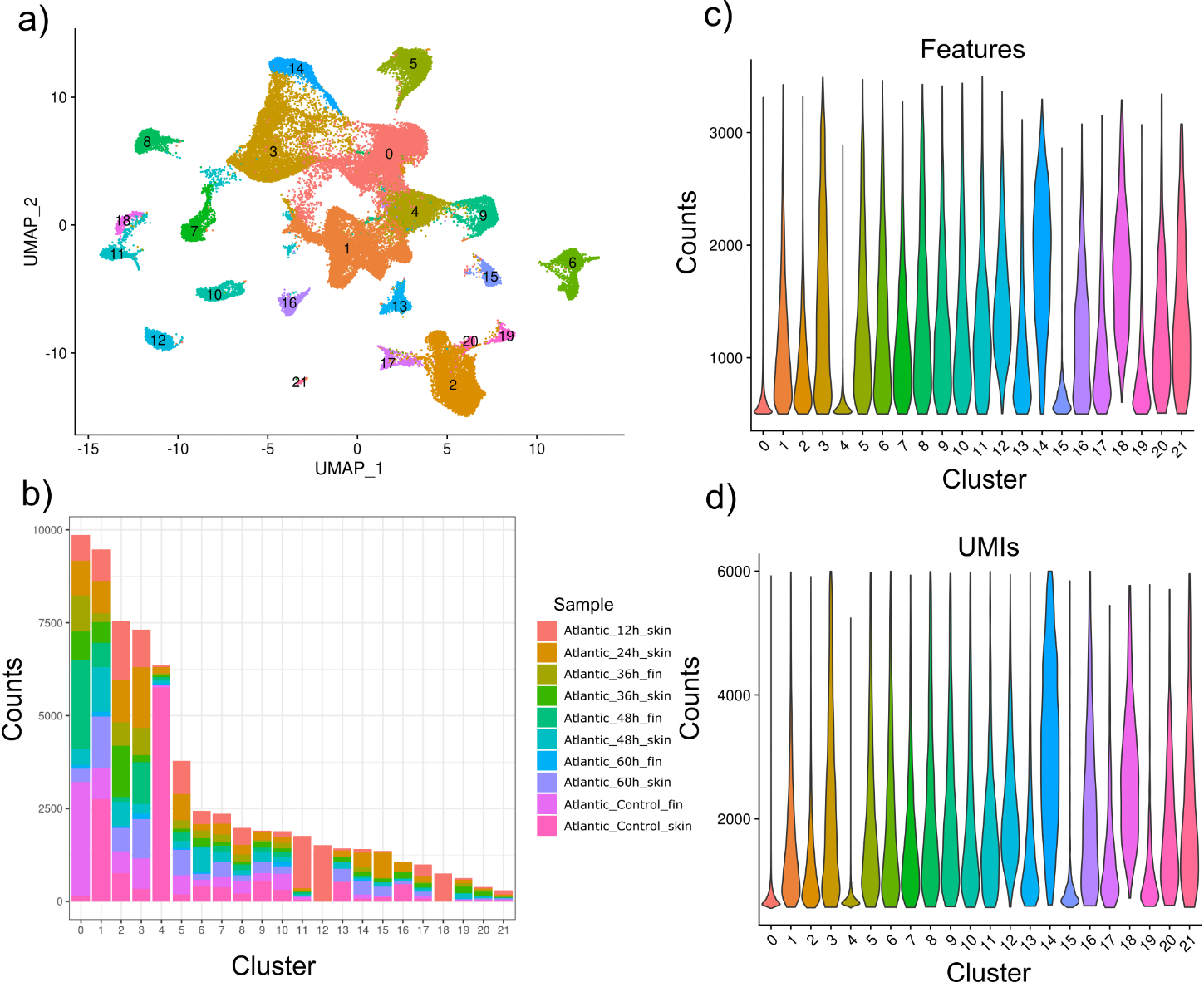

**Fig.S42** Cell clusters after initial integration of Atlantic salmon samples: a) UMAP, b) number of cells per cluster per sample, c) Violin plot of the distribution of feature counts per cluster, d) Violin plot of the distribution of UMI counts per cluster. Note that clusters 0 and 4 were removed from subsequent analysis due to low average UMI/feature counts. Cluster 4 was also found almost exclusively in a single sample (Atlantic_Control_skin).

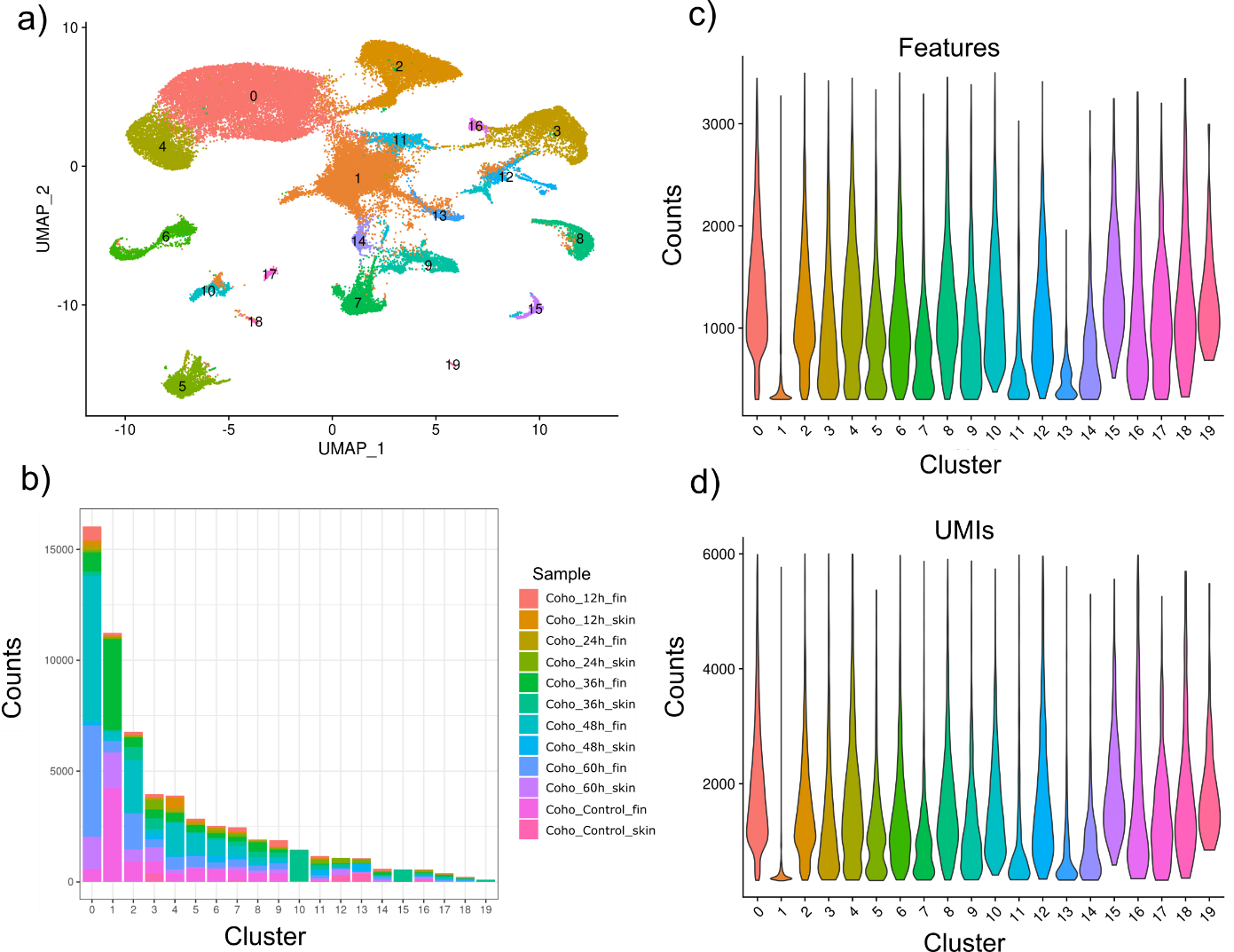

**Fig.S43** Cell clusters after initial integration of coho salmon samples: a) UMAP, b) number of cells per cluster per sample, c) Violin plot of the distribution of feature counts per cluster, d) Violin plot of the distribution of UMI counts per cluster. Note that cluster 1 was removed from subsequent analysis due to low average UMI/feature counts.

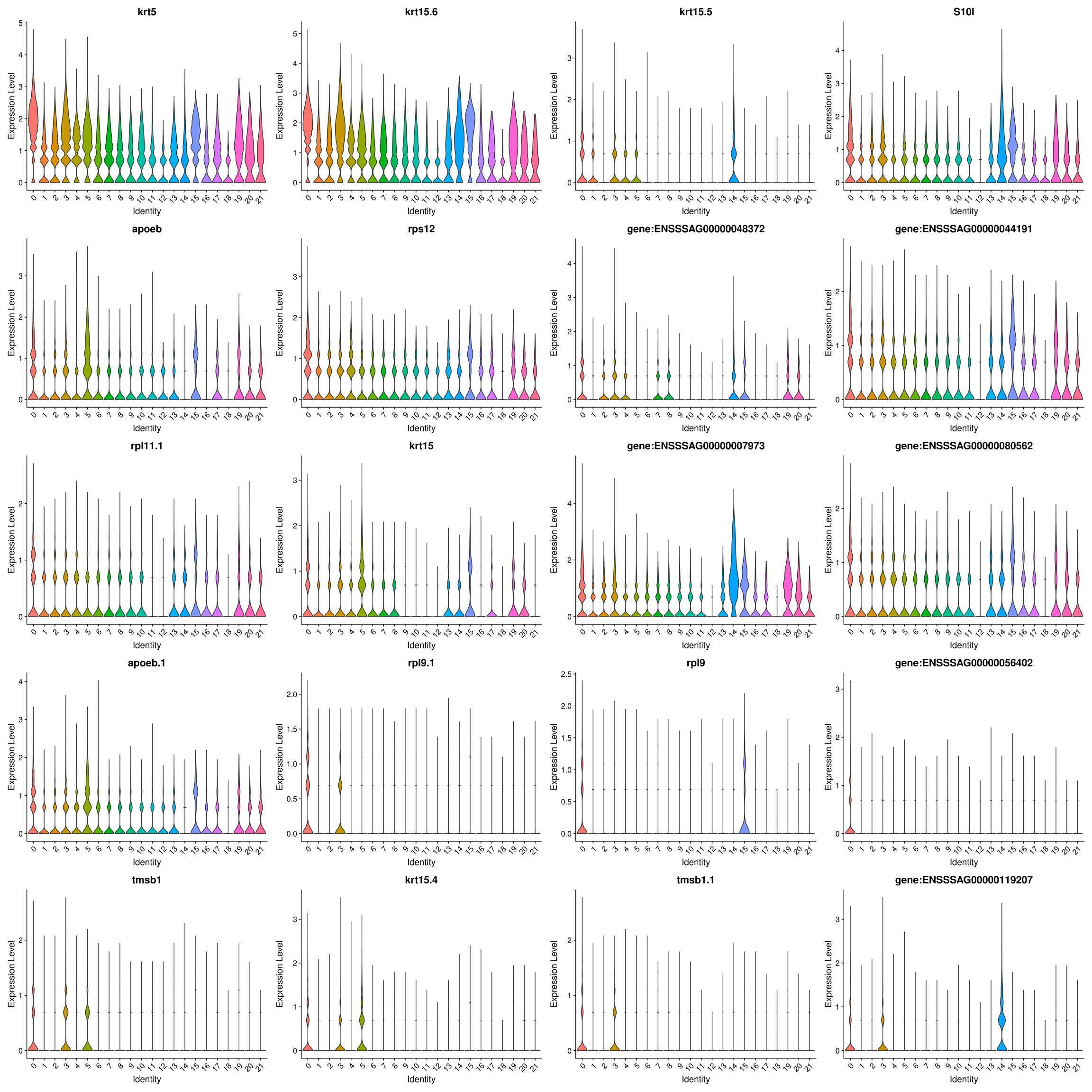

**Fig.S44** Violin plots of expression levels (based on the SCT assay) for the top 20 significant (p-adjusted < 0.05) marker genes for cluster 0 (based on log-scale two-fold change in expression) after the initial integration of Atlantic salmon samples (see Fig.S42 for UMAP). Note the abundance of ribosomal proteins.

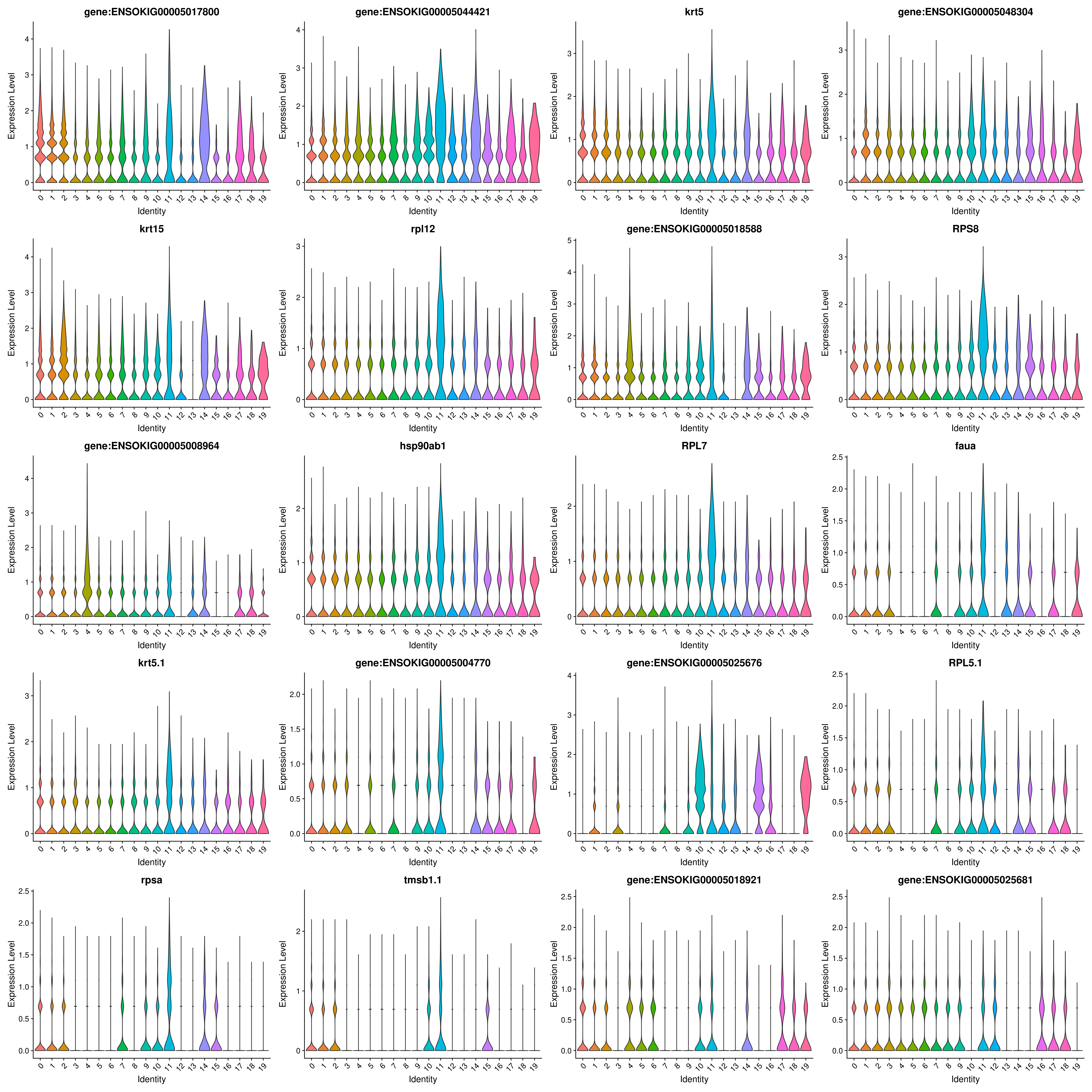

**Fig.S45** Violin plots of expression levels (based on the SCT assay) for the top 20 significant (p-adjusted < 0.05) marker genes for cluster 1 (based on log-scale two-fold change in expression) after the initial integration of coho salmon samples (see Fig.S43 for UMAP). Note the abundance of ribosomal proteins.

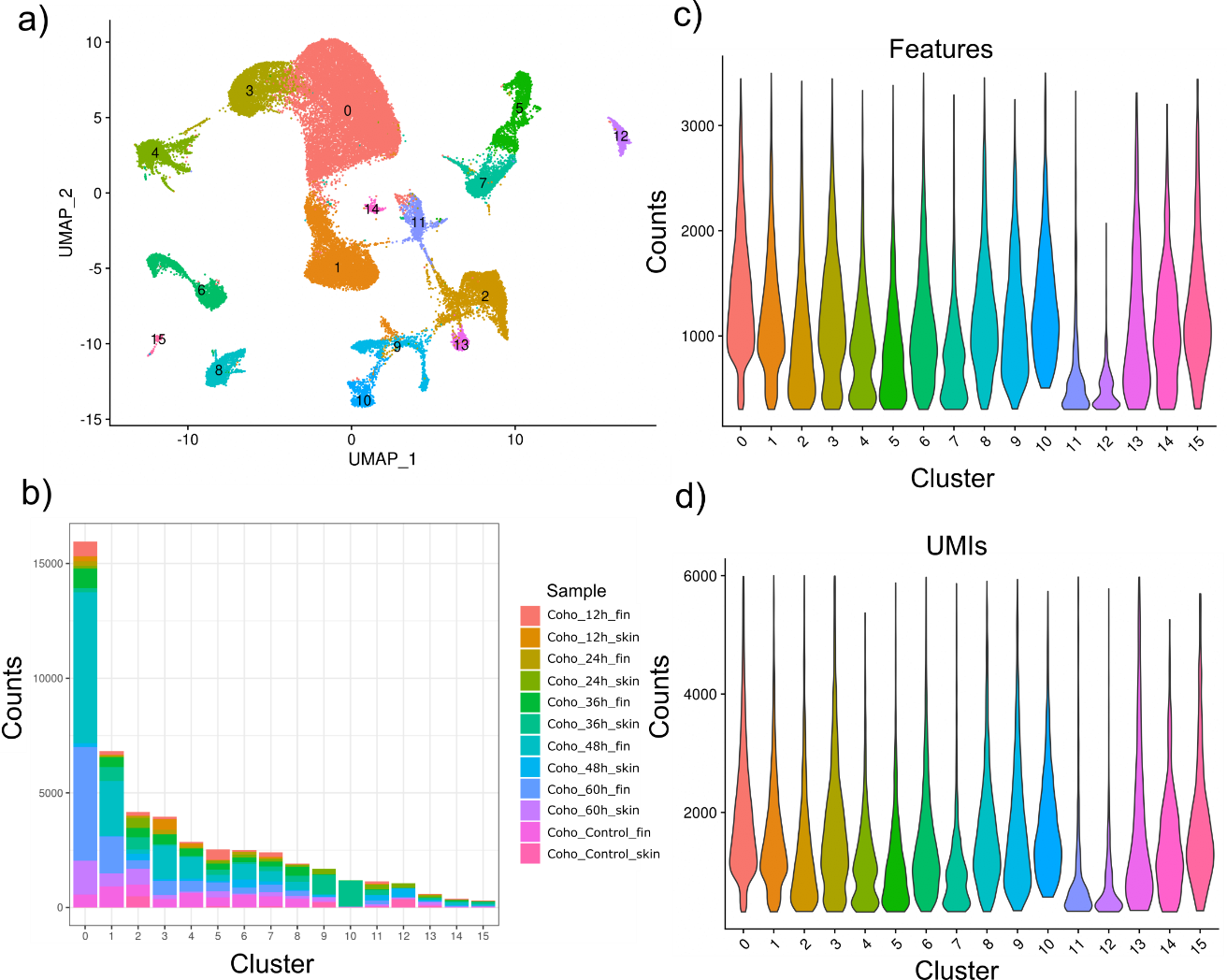

**Fig.S46** Cell clusters after removing one cluster (cluster 1 from Fig.S42) and re-integrating/clustering coho salmon samples: a) UMAP, b) number of cells per cluster per sample, c) Violin plot of the distribution of feature counts per cluster, d) Violin plot of the distribution of UMI counts per cluster.

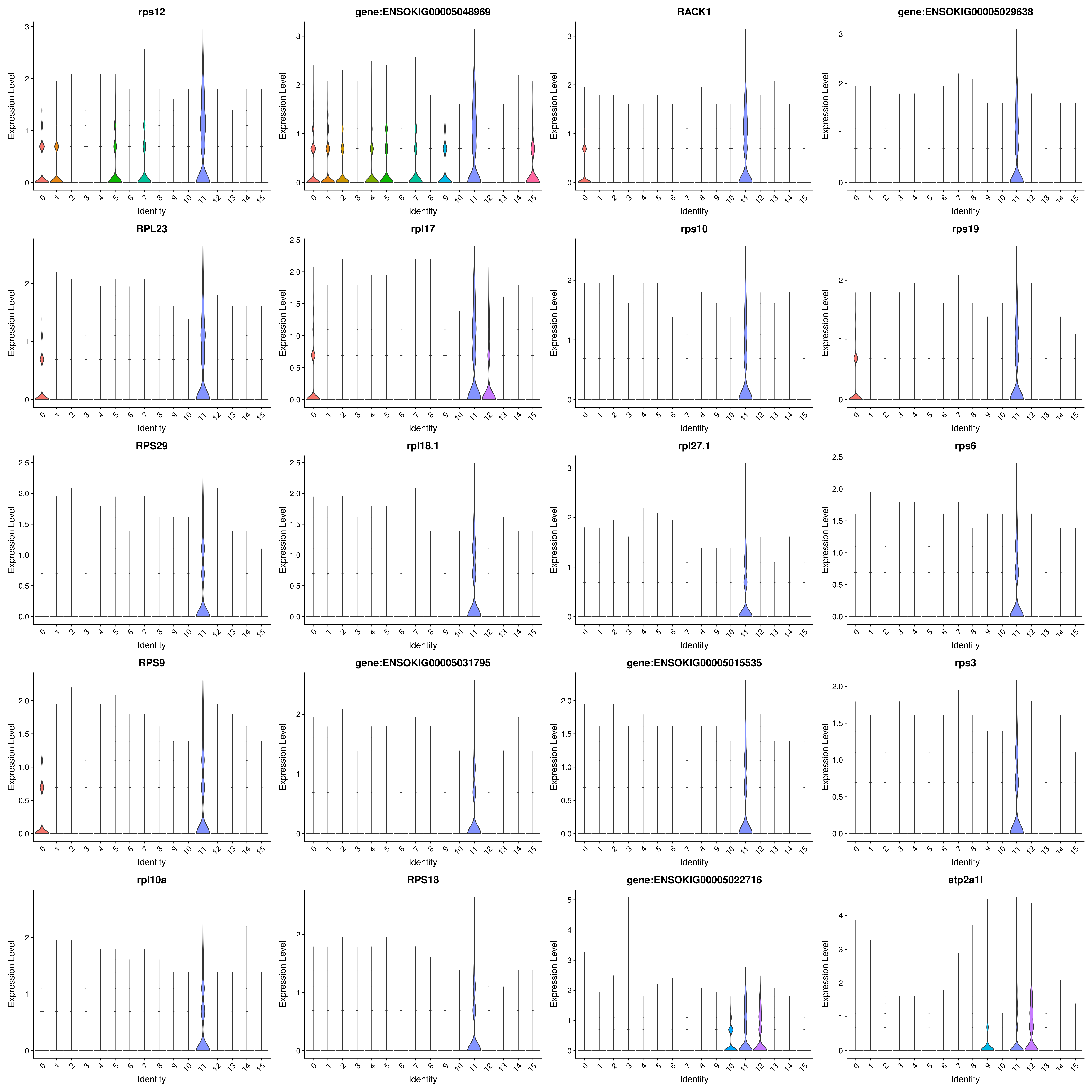
**Fig.S47** Violin plots of expression levels (based on the SCT assay) for the top 20 significant (p-adjusted < 0.05) marker genes for cluster 11 (based on log-scale two-fold change in expression) after removing one cluster (cluster 1 from Fig.S43) and re-integrating/clustering coho salmon samples (see Fig.S46 for UMAP). Note the abundance of ribosomal proteins.

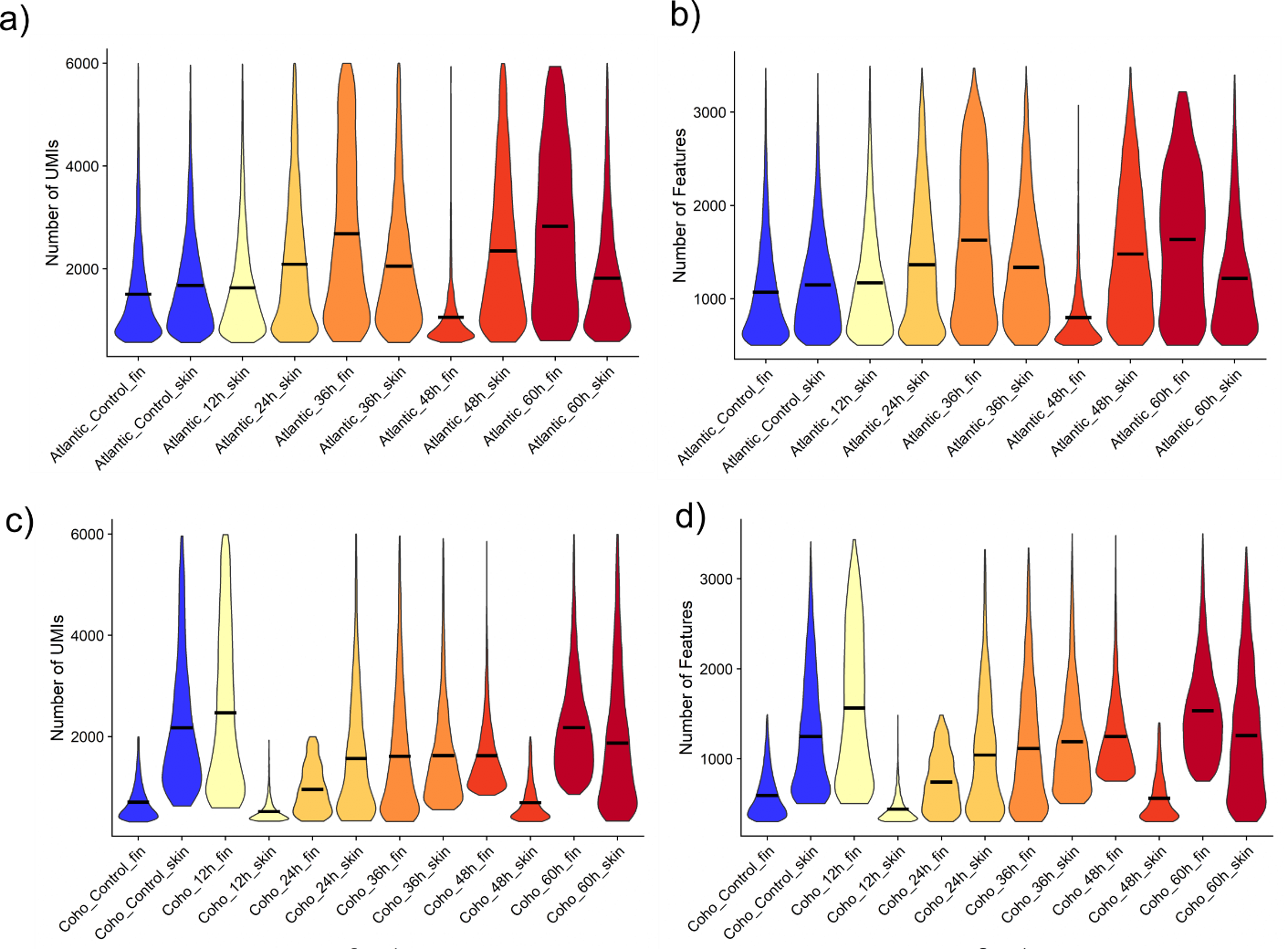

**Fig.S48** Distribution of UMIs and features for Atlantic salmon (a, b) and coho salmon (c, d). Horizontal black bar indicates the mean value for each sample.

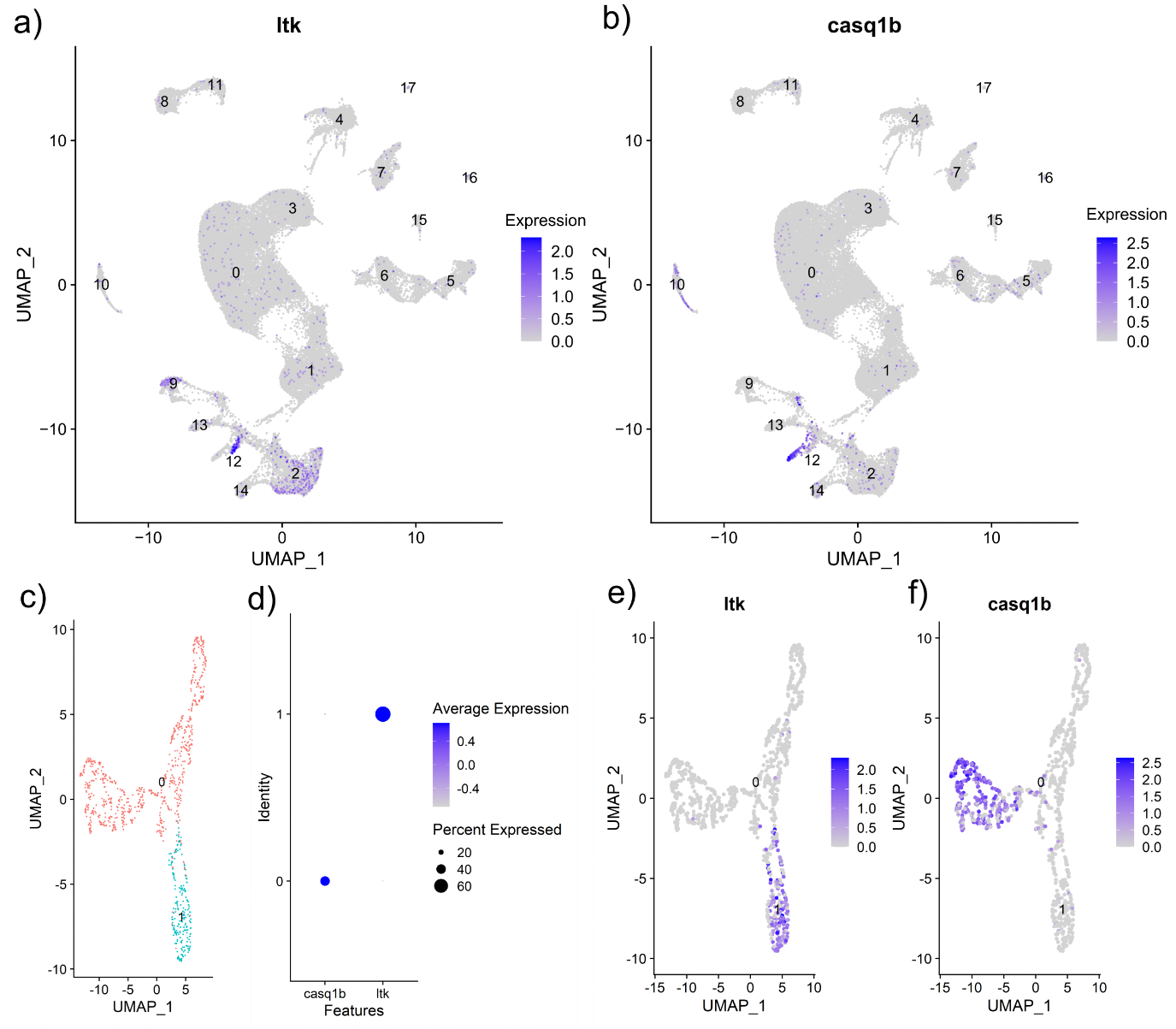

**Fig.S49** Subclustering of cluster 12 within coho salmon. Feature Plots indicate expression (based on the SCT assay) of *ltk* (a) and *casq1b* (b) in different cells within cluster 12. Re-clustering those cells within cluster 12 using 3 PCs and a resolution of 0.02 revealed two clusters as visualized in a UMAP (c). One cluster expressed *casq1b*, the other expressed *ltk* as shown in a Dot Plot (d) and Feature Plots (e, f).

**Fig.S50** Distribution of UMIs and features for each cluster identified in Atlantic salmon (a, b) and coho salmon (c, d). Horizontal black bar indicates the mean value for each sample.

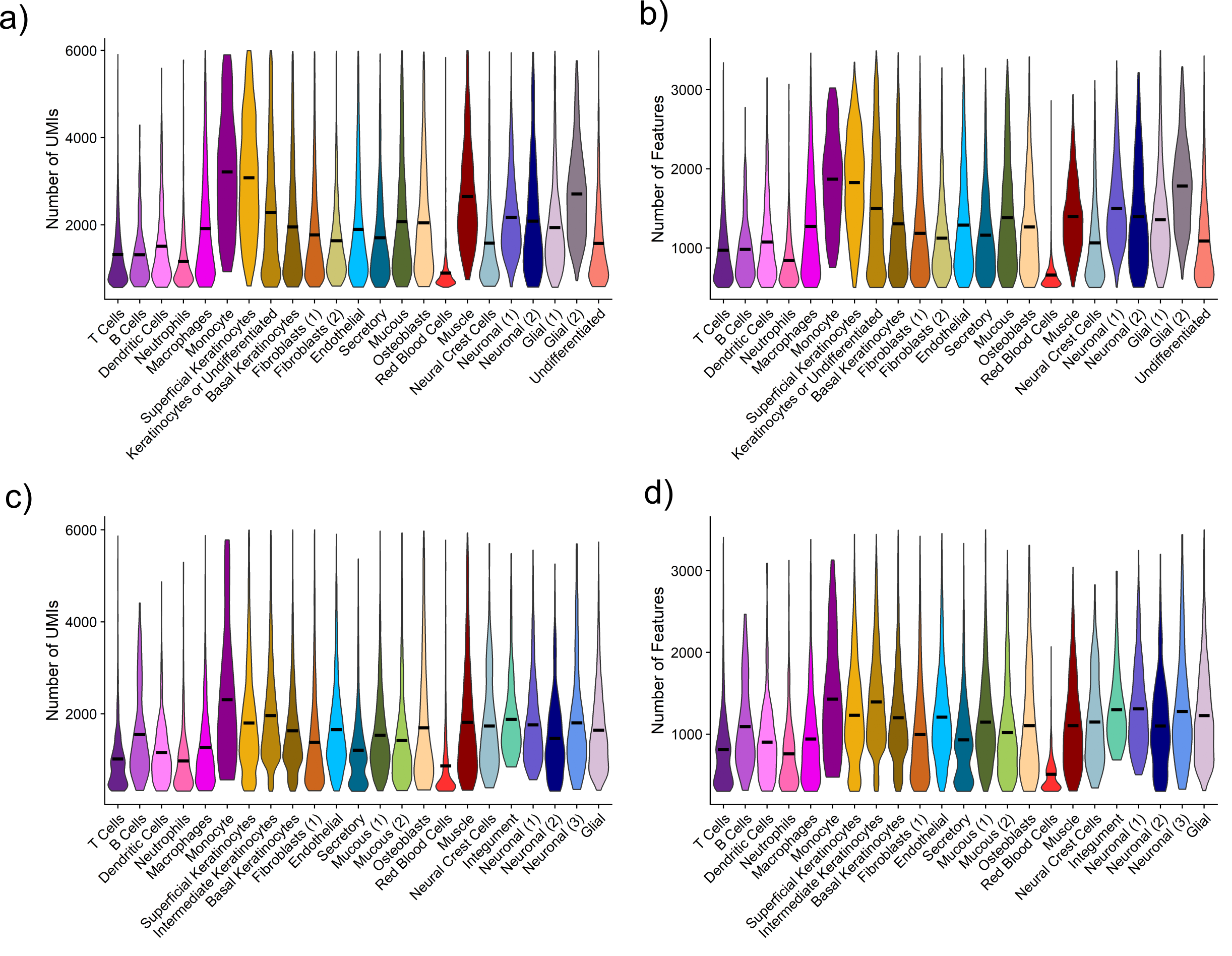

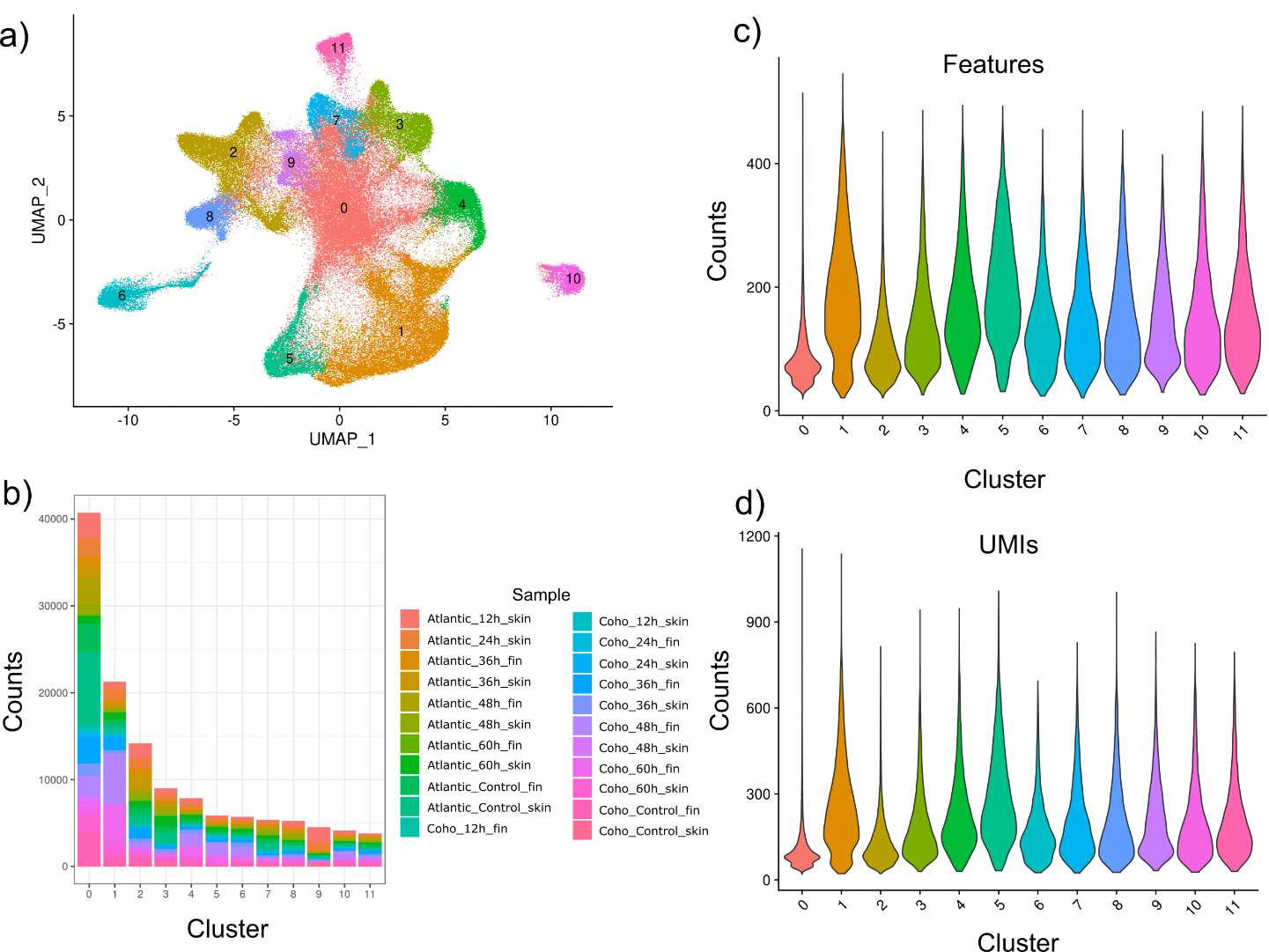

**Fig.S51** Cell clusters after initial integration of Atlantic and coho salmon samples: a) UMAP, b) number of cells per cluster per sample, c) Violin plot of the distribution of feature counts per cluster, d) Violin plot of the distribution of UMI counts per cluster. Note that cluster 0 was removed from subsequent analysis.

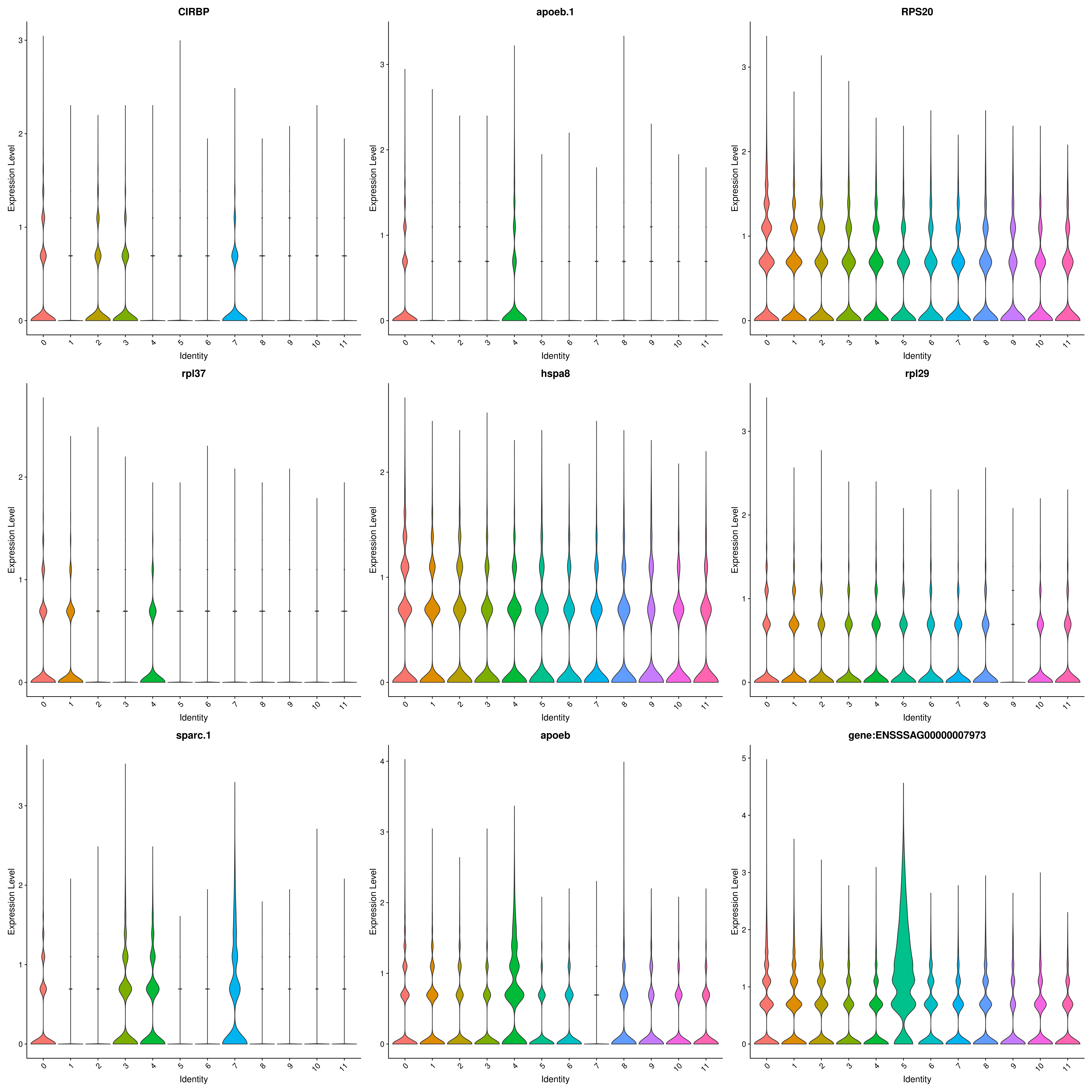
**Fig.S52** Violin plots of expression levels (based on the SCT assay) for the 9 significant (adjusted p-value < 0.05) marker genes for cluster 0 (based on log-scale two-fold change in expression) of the dataset integrating both Atlantic salmon and coho salmon samples (for UMAP see Fig.S51).

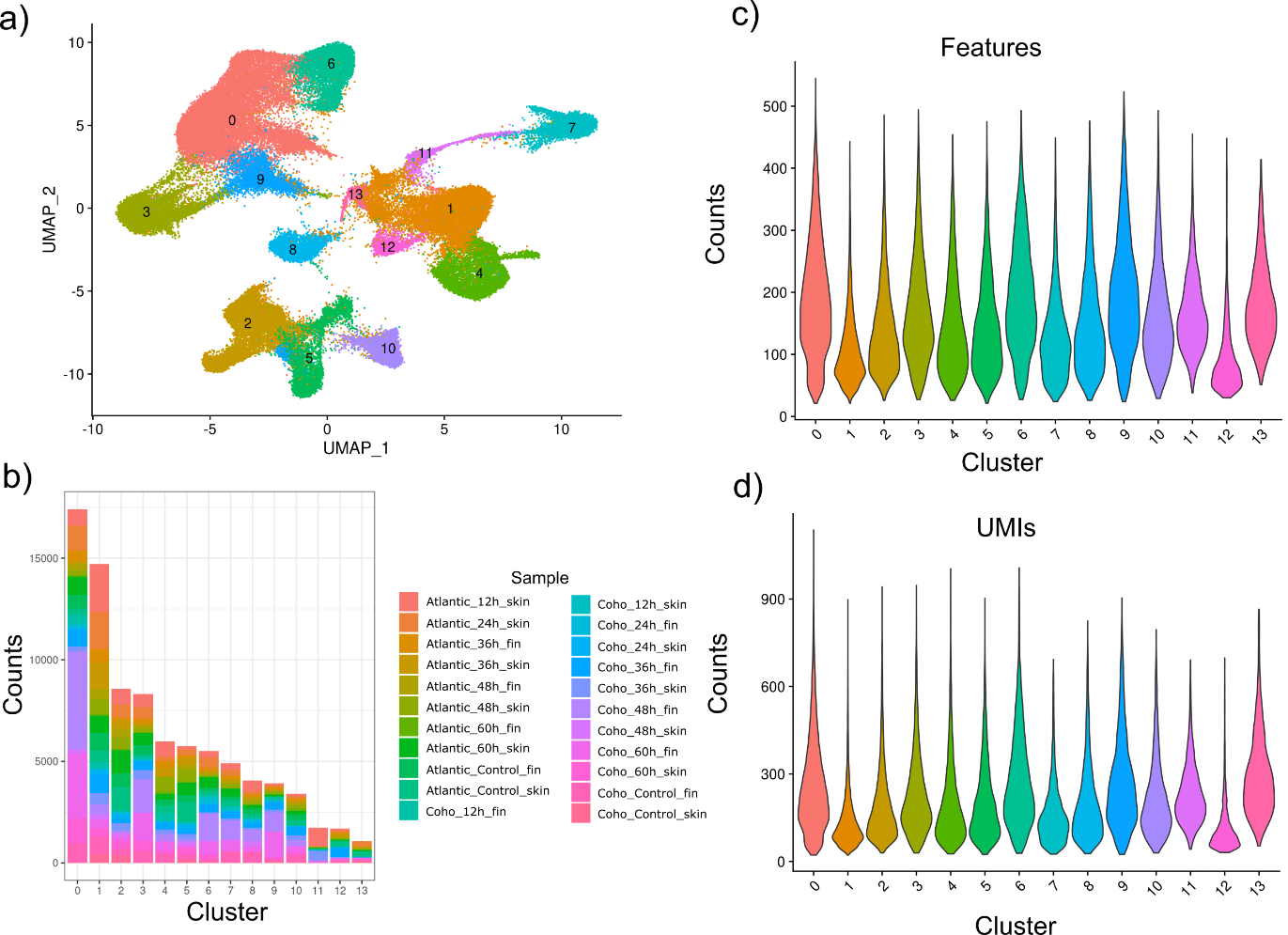

**Fig.S53** Cell clusters after removing one cluster (cluster 0 from Fig.S51) and re-integrating/clustering: a) UMAP, b) number of cells per cluster per sample, c) Violin plot of the distribution of feature counts per cluster, d) Violin plot of the distribution of UMI counts per cluster. Note that cluster 1 was removed from subsequent analysis.

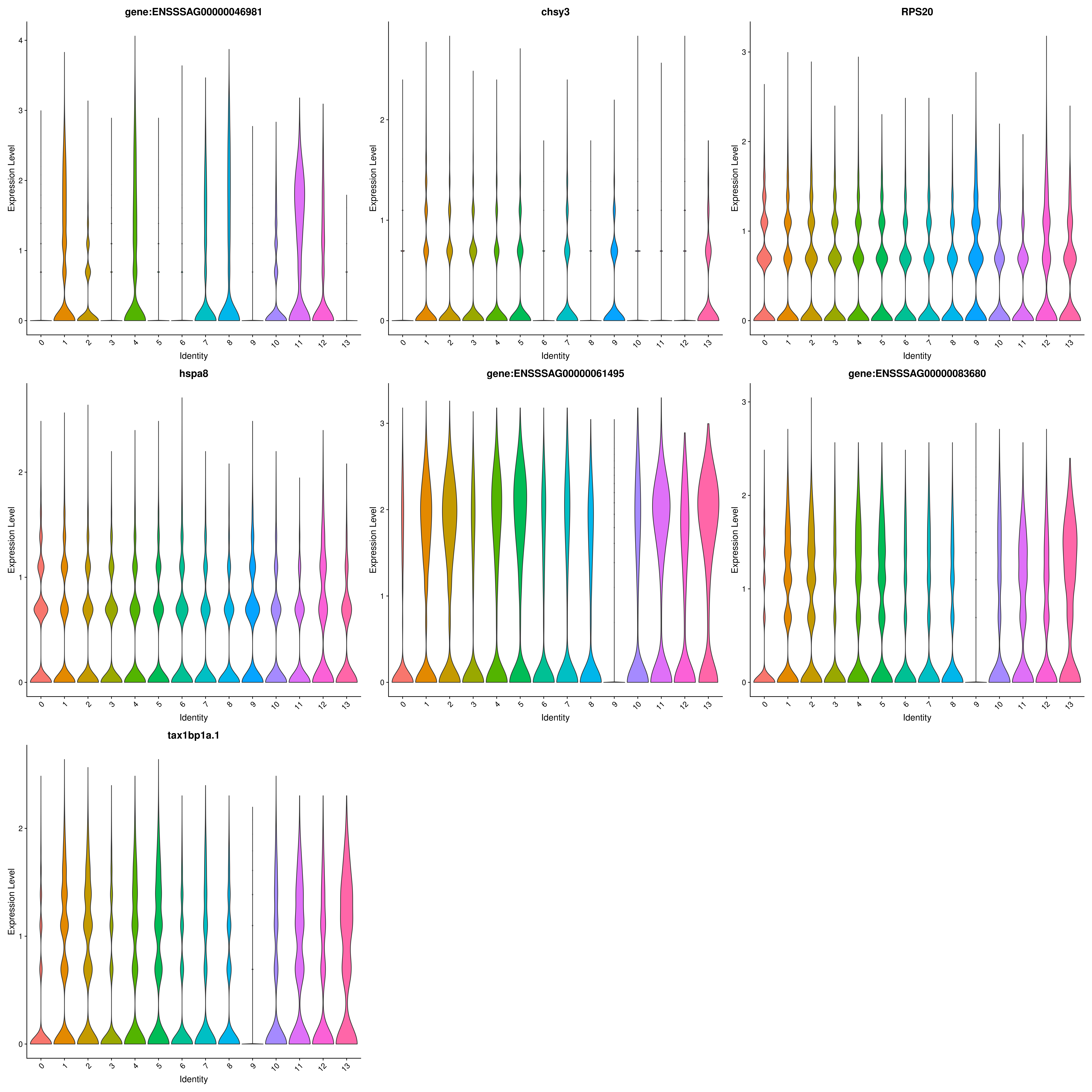

**Fig.S54** Violin plots of expression levels (based on the SCT assay) for the 7 significant (adjusted p-value < 0.05) marker genes for cluster 1 (based on log-scale two-fold change in expression) of the dataset integrating both Atlantic and Coho samples (for UMAP see Fig.S53).

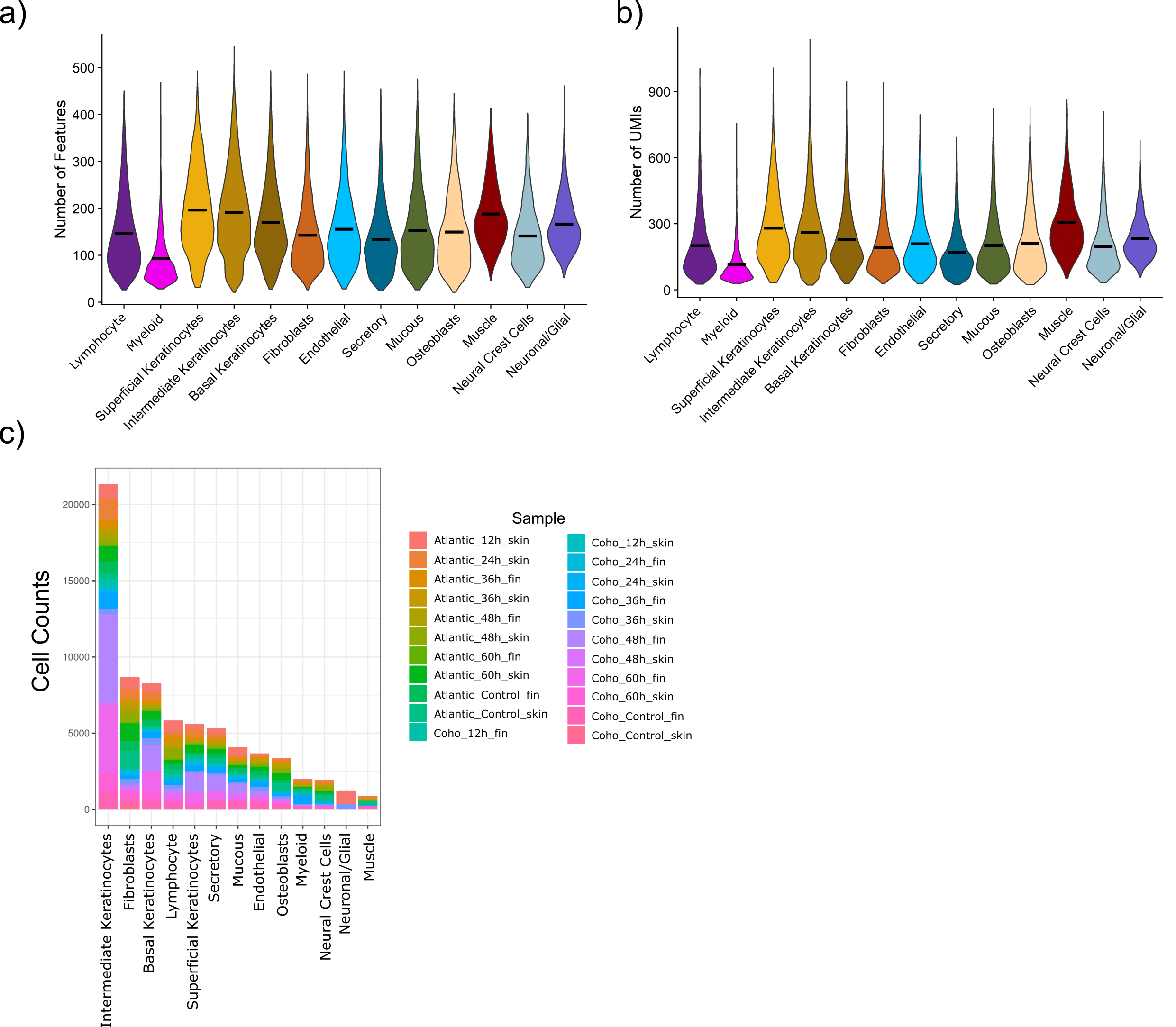

**Fig.S55** Cell clusters after removing one cluster (cluster 1 from Fig.S53) and re-integrating/clustering: a) Violin plot of the distribution of feature counts per cluster, b) Violin plot of the distribution of UMI counts per cluster, c) number of cells per cluster per sample. Horizontal white bar indicates the mean value for each sample in a and b.

**Table S1** Summary statistics for STAR outputs for each Atlantic salmon sample. Please note that raw files were used for downstream analysis (which includes all barcodes, not just those estimated to be viable cells by STAR). Therefore, the statistics relating to STAR-assigned cells are not necessarily relevant, but are provided for the reader’s interest.

| **Stats** | **Atlantic_**  **Control_**  **skin** | **Atlantic_**  **Control_**  **fin** | **Atlantic_**  **12h_**  **skin** | **Atlantic_**  **12h_**  **fin** | **Atlantic_**  **24h_**  **skin** | **Atlantic_**  **24h_**  **fin** | **Atlantic_**  **36h_**  **skin** | **Atlantic_**  **36h_**  **fin** | **Atlantic_**  **48h_**  **skin** | **Atlantic_**  **48h_**  **fin** | **Atlantic_**  **60h_**  **skin** | **Atlantic_**  **60h_**  **fin** |
| --- | --- | --- | --- | --- | --- | --- | --- | --- | --- | --- | --- | --- |
| **Number of Reads** | 321955411 | 295551337 | 287042718 | 353678800 | 282562462 | 110112712 | 315629173 | 393313790 | 264259178 | 310071740 | 263338701 | 381111089 |
| **Reads With Valid Barcodes** | 0.936286 | 0.93991 | 0.918735 | 0.966361 | 0.950211 | 0.971276 | 0.968462 | 0.955684 | 0.933665 | 0.938435 | 0.943417 | 0.961351 |
| **Sequencing Saturation** | 0.43391 | 0.481264 | 0.461568 | 0.995379 | 0.525175 | 0.899236 | 0.599346 | 0.697027 | 0.561006 | 0.54836 | 0.555422 | 0.759148 |
| **Q30 Bases in CB+UMI** | 0.965611 | 0.965352 | 0.977091 | 0.941605 | 0.969034 | 0.961646 | 0.976242 | 0.962104 | 0.976833 | 0.973577 | 0.97554 | 0.976305 |
| **Q30 Bases in RNA read** | 0.927049 | 0.925024 | 0.938372 | 0.839489 | 0.932749 | 0.937356 | 0.945648 | 0.921885 | 0.935666 | 0.928847 | 0.938162 | 0.941215 |
| **Reads Mapped to Genome: Unique+Multiple** | 0.456254 | 0.472516 | 0.469762 | 0.599451 | 0.598245 | 0.775026 | 0.47207 | 0.641069 | 0.564607 | 0.463942 | 0.501395 | 0.277742 |
| **Reads Mapped to Genome: Unique** | 0.352346 | 0.364219 | 0.360404 | 0.522241 | 0.483287 | 0.663975 | 0.368411 | 0.513612 | 0.459498 | 0.336774 | 0.391177 | 0.232388 |
| **Reads Mapped to GeneFull: Unique+Multiple GeneFull** | 0.349119 | 0.362229 | 0.344158 | 0.471235 | 0.449156 | 0.609792 | 0.360462 | 0.457549 | 0.419502 | 0.357637 | 0.386086 | 0.204369 |
| **Reads Mapped to GeneFull: Unique GeneFull** | 0.273161 | 0.286393 | 0.271067 | 0.414637 | 0.372626 | 0.532902 | 0.289934 | 0.378298 | 0.350904 | 0.262615 | 0.309893 | 0.173523 |
| **Estimated Number of Cells** | 9866 | 4708 | 8982 | 2081 | 6245 | 426 | 2795 | 3068 | 3313 | 6464 | 5732 | 878 |
| **Unique Reads in Cells Mapped to GeneFull** | 32658055 | 30082325 | 36733011 | 37934797 | 46115933 | 7627481 | 31262241 | 67110528 | 40149817 | 22545541 | 36527251 | 27350059 |
| **Fraction of Unique Reads in Cells** | 0.371344 | 0.355399 | 0.4721 | 0.258679 | 0.43799 | 0.129986 | 0.34162 | 0.451043 | 0.432977 | 0.276872 | 0.447601 | 0.41357 |
| **Mean Reads per Cell** | 3310 | 6389 | 4089 | 18229 | 7384 | 17904 | 11185 | 21874 | 12118 | 3487 | 6372 | 31150 |
| **Median Reads per Cell** | 2554 | 4893 | 3320 | 10845 | 6016 | 6720 | 8064 | 18625 | 9398 | 2513 | 5026 | 21397 |
| **UMIs in Cells** | 17226103 | 13155840 | 18750612 | 217030 | 20834285 | 775112 | 11320805 | 18310523 | 16177823 | 9416816 | 14685030 | 6378850 |
| **Mean UMI per Cell** | 1746 | 2794 | 2087 | 104 | 3336 | 1819 | 4050 | 5968 | 4883 | 1456 | 2561 | 7265 |
| **Median UMI per Cell** | 1373 | 2182 | 1701 | 77 | 2732 | 844 | 2908 | 5137 | 3803 | 1084 | 2052 | 5129 |
| **Mean GeneFull per Cell** | 1124 | 1543 | 1394 | 83 | 1906 | 1044 | 2135 | 2898 | 2480 | 898 | 1535 | 3021 |
| **Median GeneFull per Cell** | 958 | 1327 | 1211 | 68 | 1692 | 653 | 1786 | 2728 | 2155 | 752 | 1328 | 2577 |
| **Total GeneFull Detected** | 45581 | 43079 | 46794 | 21227 | 45651 | 30251 | 44217 | 44983 | 44453 | 42483 | 44293 | 40529 |

**Table S2** Summary statistics for STAR outputs for each coho salmon sample. Please note that raw files were used for downstream analysis (which includes all barcodes, not just those estimated to be viable cells by STAR). Therefore, the statistics relating to STAR-assigned cells are not necessarily relevant, but are provided for the reader’s interest.

| **Stats** | **Coho_**  **Control_**  **skin** | **Coho_**  **Control_**  **fin** | **Coho_**  **12h_**  **skin** | **Coho_**  **12h_**  **fin** | **Coho_**  **24h_**  **skin** | **Coho_**  **24h_**  **fin** | **Coho_**  **36h_**  **skin** | **Coho_**  **36h_**  **fin** | **Coho_**  **48h_**  **skin** | **Coho_**  **48h_**  **fin** | **Coho_**  **60h_**  **skin** | **Coho_**  **60h_**  **fin** |
| --- | --- | --- | --- | --- | --- | --- | --- | --- | --- | --- | --- | --- |
| **Number of Reads** | 272572603 | 276464716 | 278541953 | 275427587 | 299891579 | 244684146 | 335320286 | 381640818 | 259657264 | 311390827 | 260830113 | 420924570 |
| **Reads With Valid Barcodes** | 0.941577 | 0.920432 | 0.860409 | 0.94946 | 0.953507 | 0.893923 | 0.941946 | 0.923695 | 0.942316 | 0.897022 | 0.961607 | 0.959087 |
| **Sequencing Saturation** | 0.776908 | 0.463412 | 0.820663 | 0.606011 | 0.793786 | 0.625526 | 0.520646 | 0.681679 | 0.64073 | 0.353565 | 0.581034 | 0.574321 |
| **Q30 Bases in CB+UMI** | 0.974244 | 0.973359 | 0.972798 | 0.975214 | 0.974505 | 0.971095 | 0.973501 | 0.962014 | 0.971916 | 0.975593 | 0.972391 | 0.962382 |
| **Q30 Bases in RNA read** | 0.940897 | 0.925045 | 0.921822 | 0.936799 | 0.935628 | 0.92402 | 0.925214 | 0.911675 | 0.927226 | 0.929605 | 0.918603 | 0.910774 |
| **Reads Mapped to Genome: Unique+Multiple** | 0.535037 | 0.318965 | 0.234634 | 0.516494 | 0.486003 | 0.243276 | 0.374254 | 0.379116 | 0.371447 | 0.46242 | 0.414026 | 0.565078 |
| **Reads Mapped to Genome: Unique** | 0.473984 | 0.282777 | 0.202723 | 0.457842 | 0.428401 | 0.209409 | 0.331012 | 0.335661 | 0.31976 | 0.414474 | 0.367817 | 0.503185 |
| **Reads Mapped to GeneFull: Unique+Multiple GeneFull** | 0.383204 | 0.198197 | 0.128836 | 0.32536 | 0.296861 | 0.141313 | 0.229477 | 0.223146 | 0.216348 | 0.267375 | 0.26304 | 0.338743 |
| **Reads Mapped to GeneFull: Unique GeneFull** | 0.33425 | 0.17752 | 0.111288 | 0.289536 | 0.261617 | 0.123318 | 0.205097 | 0.19756 | 0.18971 | 0.241075 | 0.236002 | 0.30292 |
| **Estimated Number of Cells** | 1851 | 9314 | 8364 | 2424 | 1351 | 1997 | 5178 | 2937 | 2676 | 27719 | 4562 | 12757 |
| **Unique Reads in Cells Mapped to GeneFull** | 28060242 | 17302048 | 12889879 | 32016959 | 24601862 | 7203141 | 25099571 | 24465648 | 11997431 | 52586806 | 26501624 | 63480222 |
| **Fraction of Unique Reads in Cells** | 0.307991 | 0.352541 | 0.415825 | 0.401486 | 0.313572 | 0.23872 | 0.364961 | 0.324492 | 0.243555 | 0.700518 | 0.430525 | 0.497859 |
| **Mean Reads per Cell** | 15159 | 1857 | 1541 | 13208 | 18210 | 3606 | 4847 | 8330 | 4483 | 1897 | 5809 | 4976 |
| **Median Reads per Cell** | 10737 | 1378 | 1130 | 10670 | 13369 | 2832 | 3809 | 6559 | 3223 | 1503 | 4709 | 4358 |
| **UMIs in Cells** | 5979997 | 8830040 | 1977143 | 11436790 | 4554764 | 2510949 | 11232707 | 6661804 | 3817768 | 33617687 | 10393018 | 26132182 |
| **Mean UMI per Cell** | 3230 | 948 | 236 | 4718 | 3371 | 1257 | 2169 | 2268 | 1426 | 1212 | 2278 | 2048 |
| **Median UMI per Cell** | 2291 | 717 | 181 | 3877 | 2492 | 992 | 1725 | 1842 | 1026 | 972 | 1847 | 1803 |
| **Mean GeneFull per Cell** | 1465 | 667 | 205 | 2401 | 1780 | 819 | 1376 | 1441 | 805 | 940 | 1413 | 1404 |
| **Median GeneFull per Cell** | 1248 | 554 | 162 | 2178 | 1550 | 689 | 1209 | 1274 | 663 | 797 | 1247 | 1312 |
| **Total GeneFull Detected** | 35062 | 34812 | 35919 | 35011 | 34321 | 30685 | 37853 | 34492 | 32828 | 38580 | 36130 | 38153 |

**Table S3** Number of nuclei detected per sample after several filtering stages: 1) initial filtering removing nuclei with fewer than 200 features, 2) removing nuclei where mtDNA features account for 10% or more of all UMIs, 3) removing nuclei that exceeded UMI/feature upper and lower bounds, 4) removing doublet cells identified with DoubletFinder, 5) removing cells in poor quality clusters. Note that sample Atlantic_12h_fin was removed from subsequent analysis after the initial filtering due to a low cell count (highlighted). Sample Atlantic_24h_fin was removed from subsequent analysis after removing nuclei based on mtDNA features and upper and lower UMI/feature counts due to a low cell count (highlighted). Note that because only 1:1 orthologs were used for SCTransformation prior to running DoubletFinder fo the species-integrated dataset, cell counts differ for stage 4 (after removing doublet cells identified with DoubletFinder) and 5 (removing cells in poor quality clusters) between the species-integrated and species-specific datasets and are noted in separate columns. However, filtering for stages 1-3 were identical for the species-specific and species-integrated datasets.

|  | **Cell Count** | | | | | | |
| --- | --- | --- | --- | --- | --- | --- | --- |
|  | **All Datasets** | | | **Species-Specific Datasets** | | **Species-Integrated Dataset** | |
| **Sample ID** | **1) Nuclei with > 200 features** | **2) Nuclei with < 10% of UMIs as mtDNA features** | **3) Nuclei passing upper and lower UMI/feature thresholds** | **4) Nuclei after removing doublets** | **5) Nuclei after removing poor-quality clusters** | **4) Nuclei after removing doublets** | **5) Nuclei after removing poor-quality clusters** |
| **Atlantic_Control_fin** | 61866 | 55206 | 7896 | 7647 | 4555 | 7683 | 3531 |
| **Atlantic_Control_skin** | 66543 | 36325 | 13648 | 13217 | 7290 | 13242 | 4263 |
| **Atlantic_12h_fin** | 60 |  |  |  |  |  |  |
| **Atlantic_12h_skin** | 60811 | 54419 | 11139 | 10730 | 9962 | 10765 | 5515 |
| **Atlantic_24h_fin** | 434 | 431 | 338 |  |  |  |  |
| **Atlantic_24h_skin** | 64345 | 51918 | 8804 | 8488 | 7398 | 8514 | 4544 |
| **Atlantic_36h_fin** | 66375 | 66193 | 4136 | 3995 | 2981 | 4001 | 2063 |
| **Atlantic_36h_skin** | 58441 | 57179 | 4817 | 4648 | 3820 | 4656 | 2076 |
| **Atlantic_48h_fin** | 69318 | 47909 | 5547 | 5383 | 2924 | 5484 | 1916 |
| **Atlantic_48h_skin** | 60731 | 53298 | 5140 | 4961 | 4443 | 4959 | 3084 |
| **Atlantic_60h_fin** | 2190 | 2106 | 831 | 804 | 671 | 807 | 572 |
| **Atlantic_60h_skin** | 57553 | 53337 | 6928 | 6683 | 6284 | 6691 | 4987 |
| **Coho_Control_fin** | 57310 | 38435 | 9344 | 9078 | 4811 | 9145 | 4348 |
| **Coho_Control_skin** | 5427 | 1961 | 1444 | 1394 | 1341 | 1394 | 840 |
| **Coho_12h_fin** | 54254 | 45595 | 2269 | 2192 | 1969 | 2193 | 1592 |
| **Coho_12h_skin** | 2969 | 2920 | 1128 | 1096 | 1020 | 118 | 557 |
| **Coho_24h_fin** | 3040 | 1481 | 844 | 819 | 799 | 826 | 741 |
| **Coho_24h_skin** | 5642 | 4875 | 1920 | 1854 | 1579 | 1859 | 1074 |
| **Coho_36h_fin** | 46548 | 44540 | 8025 | 7803 | 3642 | 7820 | 3987 |
| **Coho_36h_skin** | 59963 | 40633 | 4901 | 4727 | 4438 | 4731 | 2682 |
| **Coho_48h_fin** | 57552 | 53458 | 14364 | 13921 | 13524 | 13967 | 11153 |
| **Coho_48h_skin** | 17454 | 8991 | 2097 | 2025 | 1755 | 2042 | 1150 |
| **Coho_60h_fin** | 62865 | 59425 | 10397 | 10051 | 9380 | 10122 | 8246 |
| **Coho_60h_skin** | 45109 | 20449 | 5952 | 5751 | 4083 | 5779 | 3418 |

**Table S4** Number of cells per cell type in each Atlantic salmon sample. Note that cell types with fewer than 50 cells in all control samples and time points with fewer than 50 cells were excluded from differential expression analyses. These excluded cell types and times are highlighted in yellow.

|  | **Samples** | | | | | | | | | |
| --- | --- | --- | --- | --- | --- | --- | --- | --- | --- | --- |
| **Cell Type** | **Atlantic_60h_fin** | **Atlantic_60h_skin** | **Atlantic_48h_fin** | **Atlantic_48h_skin** | **Atlantic_36h_fin** | **Atlantic_36h_skin** | **Atlantic_24h_skin** | **Atlantic_12h_skin** | **Atlantic_Control_fin** | **Atlantic_Control_skin** |
| Monocyte | 1 | 2 | 0 | 80 | 1 | 8 | 4 | 3 | 2 | 4 |
| Neuronal (2) | 2 | 33 | 14 | 11 | 41 | 14 | 40 | 118 | 37 | 16 |
| B Cells | 0 | 27 | 1 | 26 | 3 | 22 | 66 | 163 | 3 | 34 |
| Neutrophils | 28 | 14 | 7 | 123 | 87 | 180 | 70 | 50 | 43 | 17 |
| Muscle | 0 | 64 | 0 | 53 | 0 | 154 | 167 | 0 | 0 | 312 |
| Dendritic Cells | 9 | 94 | 37 | 53 | 26 | 97 | 86 | 191 | 72 | 95 |
| Glial (2) | 0 | 0 | 0 | 0 | 0 | 0 | 1 | 771 | 0 | 0 |
| Undifferentiated | 16 | 52 | 114 | 138 | 32 | 51 | 80 | 97 | 44 | 274 |
| Fibroblasts (2) | 0 | 0 | 3 | 200 | 1 | 107 | 3 | 1 | 6 | 927 |
| Neural Crest Cells | 0 | 344 | 0 | 178 | 0 | 139 | 170 | 47 | 2 | 506 |
| Neuronal (1) | 0 | 0 | 0 | 0 | 0 | 0 | 2 | 1510 | 0 | 0 |
| Red Blood Cells | 2 | 266 | 0 | 127 | 8 | 114 | 785 | 33 | 30 | 219 |
| Superficial Keratinocytes | 21 | 372 | 67 | 66 | 131 | 110 | 401 | 223 | 112 | 154 |
| Glial (1) | 8 | 51 | 8 | 30 | 25 | 6 | 89 | 1475 | 91 | 15 |
| Endothelial | 115 | 201 | 115 | 155 | 152 | 96 | 162 | 166 | 438 | 318 |
| Osteoblasts | 12 | 306 | 117 | 254 | 37 | 168 | 229 | 29 | 198 | 602 |
| Mucous | 19 | 165 | 161 | 124 | 183 | 85 | 274 | 458 | 332 | 212 |
| Secretory | 16 | 370 | 180 | 216 | 174 | 114 | 278 | 224 | 224 | 357 |
| Macrophages | 54 | 164 | 33 | 575 | 210 | 211 | 168 | 340 | 163 | 401 |
| Basal Keratinocytes | 52 | 680 | 357 | 195 | 235 | 100 | 714 | 926 | 534 | 191 |
| Intermediate Keratinocytes | 145 | 1075 | 1054 | 237 | 740 | 179 | 1618 | 795 | 806 | 256 |
| Fibroblasts (1) | 99 | 1342 | 498 | 912 | 215 | 420 | 794 | 746 | 811 | 1655 |
| T Cells | 72 | 662 | 158 | 690 | 680 | 1445 | 1197 | 1596 | 607 | 725 |

|  | **Samples** | | | | | | | | | | | |
| --- | --- | --- | --- | --- | --- | --- | --- | --- | --- | --- | --- | --- |
| **Cell Type** | **Coho_60h_fin** | **Coho_60h_skin** | **Coho_48h_fin** | **Coho_48h_skin** | **Coho_36h_fin** | **Coho_36h_skin** | **Coho_24h_fin** | **Coho_24h_skin** | **Coho_12h_fin** | **Coho_12h_skin** | **Coho_Control_fin** | **Coho_Control_skin** |
| B Cells | 0 | 4 | 0 | 13 | 0 | 9 | 1 | 5 | 0 | 0 | 1 | 30 |
| Monocyte | 0 | 30 | 1 | 2 | 0 | 34 | 0 | 0 | 3 | 0 | 0 | 0 |
| Integument | 0 | 6 | 0 | 0 | 0 | 87 | 0 | 0 | 0 | 0 | 0 | 0 |
| Neural Crest Cells | 2 | 4 | 0 | 30 | 2 | 5 | 0 | 117 | 5 | 0 | 1 | 44 |
| Neuronal (3) | 26 | 9 | 61 | 13 | 22 | 9 | 3 | 5 | 24 | 11 | 31 | 0 |
| Dendritic Cells | 52 | 30 | 51 | 7 | 18 | 45 | 2 | 18 | 43 | 6 | 50 | 5 |
| Neuronal (2) | 49 | 18 | 118 | 19 | 64 | 27 | 7 | 10 | 56 | 3 | 17 | 1 |
| Osteoblasts | 20 | 132 | 8 | 65 | 25 | 83 | 9 | 74 | 15 | 37 | 50 | 53 |
| Neutrophils | 134 | 80 | 39 | 11 | 120 | 27 | 2 | 51 | 116 | 1 | 34 | 6 |
| Neuronal (1) | 1 | 6 | 1 | 0 | 0 | 654 | 0 | 0 | 0 | 0 | 0 | 0 |
| Muscle | 18 | 181 | 9 | 61 | 21 | 81 | 0 | 108 | 1 | 1 | 48 | 191 |
| Mucous (2) | 74 | 45 | 136 | 345 | 26 | 91 | 43 | 115 | 23 | 2 | 116 | 16 |
| Red Blood Cells | 0 | 94 | 0 | 376 | 0 | 32 | 1 | 197 | 4 | 0 | 1 | 364 |
| Glial | 3 | 29 | 7 | 0 | 9 | 1254 | 0 | 17 | 3 | 0 | 15 | 21 |
| Macrophages | 217 | 132 | 185 | 26 | 103 | 117 | 9 | 67 | 269 | 14 | 282 | 19 |
| Mucous (1) | 187 | 17 | 544 | 0 | 177 | 5 | 65 | 5 | 54 | 15 | 418 | 1 |
| Endothelial | 234 | 122 | 330 | 44 | 399 | 265 | 34 | 70 | 26 | 1 | 362 | 19 |
| T Cells | 321 | 191 | 455 | 127 | 225 | 190 | 25 | 115 | 209 | 63 | 457 | 53 |
| Secretory | 484 | 63 | 945 | 91 | 328 | 43 | 61 | 23 | 64 | 132 | 602 | 18 |
| Superficial Keratinocytes | 585 | 204 | 1358 | 59 | 413 | 57 | 195 | 21 | 105 | 418 | 222 | 5 |
| Fibroblasts (1) | 356 | 655 | 191 | 276 | 405 | 502 | 83 | 428 | 156 | 22 | 518 | 466 |
| Basal Keratinocytes | 1620 | 589 | 2422 | 15 | 424 | 631 | 59 | 32 | 146 | 27 | 904 | 11 |
| Intermediate Keratinocytes | 4997 | 1442 | 6663 | 175 | 861 | 190 | 200 | 101 | 647 | 267 | 682 | 18 |

**Table S5** Number of cells per cell type in each coho salmon sample. Note that cell types with fewer than 50 cells in all control samples and time points with fewer than 50 cells were excluded from differential expression analyses. These excluded cell types and times are highlighted in yellow.

**Table S6** Number of cells per cluster and sampling time point after reclustering only the immune cells within the Atlantic salmon samples.

|  | **Sample Time Point** | | | | | |
| --- | --- | --- | --- | --- | --- | --- |
|  | **12h** | **24h** | **36h** | **48h** | **60h** | **Control** |
| T Cells (1) | 909 | 528 | 976 | 410 | 371 | 491 |
| T Cells (2) | 499 | 432 | 467 | 206 | 113 | 486 |
| Macrophages (1) | 121 | 30 | 168 | 429 | 97 | 268 |
| Macrophages (2) | 183 | 89 | 181 | 110 | 111 | 155 |
| T Cells (3) | 57 | 61 | 367 | 57 | 99 | 133 |
| Dendritic Cells | 191 | 86 | 123 | 90 | 103 | 167 |
| T Cells (4) | 92 | 121 | 157 | 90 | 108 | 153 |
| Neutrophils | 50 | 70 | 267 | 130 | 42 | 60 |
| T Cells (5) | 39 | 55 | 158 | 85 | 43 | 69 |
| Macrophages (3) | 36 | 49 | 72 | 69 | 10 | 141 |
| B Cells | 163 | 66 | 25 | 27 | 27 | 37 |
| Monocyte | 3 | 4 | 9 | 80 | 3 | 6 |

**Table S7** Number of cells per cluster after reclustering only the immune cells within the coho salmon samples.

|  | **Sample Time Point** | | | | | |
| --- | --- | --- | --- | --- | --- | --- |
| **Cluster** | **12h** | **24h** | **36h** | **48h** | **60h** | **Control** |
| T Cells (1) | 155 | 101 | 213 | 271 | 239 | 236 |
| Macrophages (1) | 200 | 17 | 118 | 181 | 188 | 61 |
| Macrophages (2) | 83 | 59 | 102 | 30 | 161 | 240 |
| Neutrophils | 117 | 53 | 147 | 50 | 214 | 40 |
| T Cells (2) | 36 | 31 | 124 | 30 | 173 | 209 |
| Dendritic Cells | 49 | 20 | 63 | 58 | 82 | 55 |
| T Cells (3) | 1 | 0 | 7 | 229 | 22 | 2 |
| T Cells (4) | 17 | 5 | 41 | 36 | 61 | 17 |
| T Cells (5) | 63 | 3 | 30 | 16 | 17 | 46 |
| Monocyte | 3 | 0 | 34 | 3 | 30 | 0 |
| B Cells | 0 | 6 | 9 | 13 | 4 | 31 |

**Table S8** Genes used for cell cycle scoring for Atlantic salmon.

| **Human_Symbol** | **Ensembl_Symbol** | **Ensembl_ID** | **Cycle** |
| --- | --- | --- | --- |
| MCM5 | mcm5 | ENSSSAG00000040979 | S.gene |
| MCM5 | mcm5.1 | ENSSSAG00000005359 | S.gene |
| PCNA | pcna | ENSSSAG00000009453 | S.gene |
| TYMS | tyms | ENSSSAG00000073844 | S.gene |
| TYMS | TYMS | ENSSSAG00000070708 | S.gene |
| MCM2 | mcm2 | ENSSSAG00000042717 | S.gene |
| MCM4 | mcm4 | ENSSSAG00000081704 | S.gene |
| MCM4 | mcm4.1 | ENSSSAG00000070777 | S.gene |
| GINS2 | psf2 | ENSSSAG00000117735 | S.gene |
| CDCA7 | cdca7a | ENSSSAG00000071616 | S.gene |
| DTL | DTL | ENSSSAG00000074405 | S.gene |
| DTL | DTL.1 | ENSSSAG00000058407 | S.gene |
| UHRF1 | uhrf1 | ENSSSAG00000072643 | S.gene |
| UHRF1 | uhrf1.1 | ENSSSAG00000046494 | S.gene |
| RFC2 | rfc2 | ENSSSAG00000066661 | S.gene |
| NASP | nasp | ENSSSAG00000049742 | S.gene |
| RAD51AP1 | rad51ap1 | ENSSSAG00000046281 | S.gene |
| RAD51AP1 | rad51ap1.1 | ENSSSAG00000002591 | S.gene |
| GMNN | gene:ENSSSAG00000052775 | ENSSSAG00000052775 | S.gene |
| WDR76 | gene:ENSSSAG00000052378 | ENSSSAG00000052378 | S.gene |
| SLBP | slbp | ENSSSAG00000063884 | S.gene |
| CCNE2 | ccne2 | ENSSSAG00000059043 | S.gene |
| CCNE2 | ccne2.1 | ENSSSAG00000057367 | S.gene |
| UBR7 | ubr7 | ENSSSAG00000040956 | S.gene |
| POLD3 | gene:ENSSSAG00000005314 | ENSSSAG00000005314 | S.gene |
| MSH2 | msh2 | ENSSSAG00000074238 | S.gene |
| ATAD2 | ATAD2 | ENSSSAG00000059621 | S.gene |
| ATAD2 | ATAD2.1 | ENSSSAG00000003726 | S.gene |
| RAD51 | rad51 | ENSSSAG00000045336 | S.gene |
| RRM2 | RRM2 | ENSSSAG00000083858 | S.gene |
| CDC45 | cdc45 | ENSSSAG00000047205 | S.gene |
| CDC45 | cdc45.1 | ENSSSAG00000047454 | S.gene |
| CDC6 | cdc6 | ENSSSAG00000041769 | S.gene |
| EXO1 | exo1 | ENSSSAG00000103371 | S.gene |
| TIPIN | tipin | ENSSSAG00000082602 | S.gene |
| TIPIN | tipin.1 | ENSSSAG00000043439 | S.gene |
| DSCC1 | dscc1 | ENSSSAG00000088925 | S.gene |
| DSCC1 | dscc1.1 | ENSSSAG00000104335 | S.gene |
| DSCC1 | DSCC1 | ENSSSAG00000098087 | S.gene |
| BLM | blm | ENSSSAG00000078058 | S.gene |
| CASP8AP2 | casp8ap2 | ENSSSAG00000045001 | S.gene |
| CASP8AP2 | casp8ap2.1 | ENSSSAG00000101584 | S.gene |
| USP1 | usp1 | ENSSSAG00000044868 | S.gene |
| CLSPN | CLSPN | ENSSSAG00000054174 | S.gene |
| CLSPN | CLSPN.1 | ENSSSAG00000078544 | S.gene |
| POLA1 | pola1 | ENSSSAG00000078390 | S.gene |
| CHAF1B | chaf1b | ENSSSAG00000042565 | S.gene |
| BRIP1 | brip1 | ENSSSAG00000051666 | S.gene |
| E2F8 | e2f8 | ENSSSAG00000065538 | S.gene |
| E2F8 | e2f8.1 | ENSSSAG00000073983 | S.gene |
| HMGB2 | hmgb2a | ENSSSAG00000054548 | G2M.gene |
| HMGB2 | hmgb2a.1 | ENSSSAG00000109524 | G2M.gene |
| CDK1 | cdk1 | ENSSSAG00000006872 | G2M.gene |
| NUSAP1 | nusap1 | ENSSSAG00000000078 | G2M.gene |
| NUSAP1 | nusap1.1 | ENSSSAG00000063363 | G2M.gene |
| BIRC5 | birc5a | ENSSSAG00000001277 | G2M.gene |
| BIRC5 | birc5b | ENSSSAG00000068971 | G2M.gene |
| BIRC5 | birc5b.1 | ENSSSAG00000095509 | G2M.gene |
| TPX2 | gene:ENSSSAG00000075691 | NSSSAG00000075691 | G2M.gene |
| TOP2A | gene:ENSSSAG00000066721 | ENSSSAG00000066721 | G2M.gene |
| NDC80 | ndc80 | ENSSSAG00000075612 | G2M.gene |
| CKS2 | cks2 | ENSSSAG00000063439 | G2M.gene |
| CKS2 | CKS2 | ENSSSAG00000053543 | G2M.gene |
| NUF2 | nuf2 | ENSSSAG00000091386 | G2M.gene |
| TMPO | tmpoa | ENSSSAG00000097715 | G2M.gene |
| TMPO | lap2 | ENSSSAG00000070216 | G2M.gene |
| TMPO | tmpob | ENSSSAG00000079549 | G2M.gene |
| TMPO | tmpob.1 | ENSSSAG00000071470 | G2M.gene |
| CENPF | gene:ENSSSAG00000002402 | ENSSSAG00000002402 | G2M.gene |
| TACC3 | gene:ENSSSAG00000043775 | ENSSSAG00000043775 | G2M.gene |
| SMC4 | smc4 | ENSSSAG00000074267 | G2M.gene |
| CCNB2 | ccnb2 | ENSSSAG00000072975 | G2M.gene |
| CCNB2 | ccnb2.1 | ENSSSAG00000080343 | G2M.gene |
| CKAP2L | ckap2l | ENSSSAG00000045445 | G2M.gene |
| KIF11 | kif11 | ENSSSAG00000006907 | G2M.gene |
| KIF11 | kif11.1 | ENSSSAG00000071263 | G2M.gene |
| ANP32E | ANP32E | ENSSSAG00000068348 | G2M.gene |
| ANP32E | ANP32E.1 | ENSSSAG00000041648 | G2M.gene |
| TUBB4B | tubb4b | ENSSSAG00000101622 | G2M.gene |
| KIF20B | kif20bb | ENSSSAG00000079056 | G2M.gene |
| KIF20B | kif20bb.1 | ENSSSAG00000068302 | G2M.gene |
| KIF20B | kif20ba | ENSSSAG00000078645 | G2M.gene |
| KIF20B | kif20ba.1 | ENSSSAG00000054281 | G2M.gene |
| CDCA3 | si:ch211-69g19.2 | ENSSSAG00000040417 | G2M.gene |
| CDC20 | cdc20 | ENSSSAG00000075816 | G2M.gene |
| CDC20 | cdc20.1 | ENSSSAG00000075603 | G2M.gene |
| TTK | ttk | ENSSSAG00000049671 | G2M.gene |
| TTK | ttk.1 | ENSSSAG00000009759 | G2M.gene |
| CDC25C | CDC25C | ENSSSAG00000006736 | G2M.gene |
| KIF2C | kif2c | ENSSSAG00000053000 | G2M.gene |
| KIF2C | kif2c.1 | ENSSSAG00000080527 | G2M.gene |
| RANGAP1 | RANGAP1 | ENSSSAG00000002274 | G2M.gene |
| RANGAP1 | RANGAP1.1 | ENSSSAG00000067438 | G2M.gene |
| RANGAP1 | rangap1b | ENSSSAG00000002137 | G2M.gene |
| DLGAP5 | dlgap5 | ENSSSAG00000000679 | G2M.gene |
| DLGAP5 | dlgap5.1 | ENSSSAG00000057319 | G2M.gene |
| CDCA8 | cdca8.1 | ENSSSAG00000040950 | G2M.gene |
| CDCA8 | cdca8.1 | ENSSSAG00000078302 | G2M.gene |
| ECT2 | ect2 | ENSSSAG00000070771 | G2M.gene |
| ECT2 | ect2.1 | ENSSSAG00000068867 | G2M.gene |
| KIF23 | KIF23 | ENSSSAG00000120697 | G2M.gene |
| KIF23 | KIF23.1 | ENSSSAG00000095510 | G2M.gene |
| HMMR | hmmr | ENSSSAG00000041733 | G2M.gene |
| AURKA | gene:ENSSSAG00000031237 | ENSSSAG00000031237 | G2M.gene |
| ANLN | ANLN | ENSSSAG00000073106 | G2M.gene |
| ANLN | ANLN.1 | ENSSSAG00000086325 | G2M.gene |
| LBR | lbr | ENSSSAG00000044615 | G2M.gene |
| LBR | lbr.1 | ENSSSAG00000076862 | G2M.gene |
| CKAP5 | ckap5 | ENSSSAG00000008344 | G2M.gene |
| CTCF | ctcf | ENSSSAG00000064848 | G2M.gene |
| CTCF | ctcf.1 | ENSSSAG00000067932 | G2M.gene |
| NEK2 | nek2 | ENSSSAG00000106767 | G2M.gene |
| NEK2 | nek2.1 | ENSSSAG00000056109 | G2M.gene |
| G2E3 | G2E3 | ENSSSAG00000064280 | G2M.gene |
| G2E3 | g2e3 | ENSSSAG00000071501 | G2M.gene |
| CBX5 | cbx5 | ENSSSAG00000096644 | G2M.gene |
| CBX5 | cbx5.1 | ENSSSAG00000110153 | G2M.gene |
| CENPA | gene:ENSSSAG00000078816 | ENSSSAG00000078816 | G2M.gene |

**Table S9** Genes used for cell cycle scoring for coho salmon.

| **Human_Symbol** | **Ensembl_Symbol** | **Ensembl_ID** | **Cycle** |
| --- | --- | --- | --- |
| MCM5 | mcm5 | ENSOKIG00005014560 | S.gene |
| MCM5 | mcm5.1 | ENSOKIG00005018292 | S.gene |
| PCNA | pcna | ENSOKIG00005011301 | S.gene |
| PCNA | PCNA.1 | ENSOKIG00005018627 | S.gene |
| PCNA | PCNA | ENSOKIG00005029428 | S.gene |
| FEN1 | FEN1 | ENSOKIG00005024494 | S.gene |
| FEN1 | FEN1.1 | ENSOKIG00005024547 | S.gene |
| FEN1 | FEN1.2 | ENSOKIG00005024619 | S.gene |
| MCM2 | mcm2 | ENSOKIG00005004073 | S.gene |
| MCM2 | MCM2 | ENSOKIG00005017128 | S.gene |
| MCM4 | mcm4 | ENSOKIG00005012623 | S.gene |
| MCM4 | mcm4.1 | ENSOKIG00005042602 | S.gene |
| RRM1 | rrm1.1 | ENSOKIG00005024782 | S.gene |
| RRM1 | rrm1 | ENSOKIG00005046893 | S.gene |
| UNG | unga | ENSOKIG00005050249 | S.gene |
| GINS2 | gins2 | ENSOKIG00005005401 | S.gene |
| MCM6 | MCM6 | ENSOKIG00005040687 | S.gene |
| CDCA7 | cdca7a | ENSOKIG00005013961 | S.gene |
| DTL | DTL | ENSOKIG00005026623 | S.gene |
| DTL | dtl | ENSOKIG00005013017 | S.gene |
| PRIM1 | prim1 | ENSOKIG00005010009 | S.gene |
| UHRF1 | uhrf1.1 | ENSOKIG00005000817 | S.gene |
| UHRF1 | uhrf1 | ENSOKIG00005015452 | S.gene |
| MLF1IP | gene:ENSOKIG00005012480 | ENSOKIG00005012480 | S.gene |
| HELLS | hells.1 | ENSOKIG00005009326 | S.gene |
| HELLS | hells | ENSOKIG00005047721 | S.gene |
| RFC2 | rfc2 | ENSOKIG00005029854 | S.gene |
| NASP | NASP | ENSOKIG00005038416 | S.gene |
| NASP | nasp | ENSOKIG00005015929 | S.gene |
| RAD51AP1 | rad51ap1 | ENSOKIG00005006131 | S.gene |
| RAD51AP1 | rad51ap1.1 | ENSOKIG00005031433 | S.gene |
| GMNN | GMNN | ENSOKIG00005004827 | S.gene |
| GMNN | gmnn | ENSOKIG00005010054 | S.gene |
| WDR76 | wdr76 | ENSOKIG00005007563 | S.gene |
| SLBP | SLBP | ENSOKIG00005007467 | S.gene |
| SLBP | slbp | ENSOKIG00005019708 | S.gene |
| CCNE2 | ccne2 | ENSOKIG00005032974 | S.gene |
| CCNE2 | ccne2.1 | ENSOKIG00005011102 | S.gene |
| UBR7 | gene:ENSOKIG00005002397 | ENSOKIG00005002397 | S.gene |
| UBR7 | ubr7 | ENSOKIG00005004167 | S.gene |
| POLD3 | gene:ENSOKIG00005049692 | ENSOKIG00005049692 | S.gene |
| MSH2 | msh2 | ENSOKIG00005029611 | S.gene |
| ATAD2 | ATAD2.1 | ENSOKIG00005045862 | S.gene |
| ATAD2 | ATAD2 | ENSOKIG00005004525 | S.gene |
| ATAD2 | atad2 | ENSOKIG00005047340 | S.gene |
| RAD51 | rad51 | ENSOKIG00005006788 | S.gene |
| RAD51 | RAD51 | ENSOKIG00005029092 | S.gene |
| RRM2 | RRM2 | ENSOKIG00005002061 | S.gene |
| RRM2 | RRM2.1 | ENSOKIG00005036867 | S.gene |
| CDC45 | cdc45 | ENSOKIG00005023527 | S.gene |
| CDC45 | cdc45.1 | ENSOKIG00005020780 | S.gene |
| CDC6 | cdc6.1 | ENSOKIG00005005964 | S.gene |
| CDC6 | cdc6 | ENSOKIG00005004472 | S.gene |
| EXO1 | exo1 | ENSOKIG00005011806 | S.gene |
| DSCC1 | dscc1.1 | ENSOKIG00005047842 | S.gene |
| DSCC1 | dscc1 | ENSOKIG00005048820 | S.gene |
| BLM | blm | ENSOKIG00005006801 | S.gene |
| CASP8AP2 | casp8ap2.1 | ENSOKIG00005009010 | S.gene |
| CASP8AP2 | casp8ap2 | ENSOKIG00005002883 | S.gene |
| USP1 | usp1 | ENSOKIG00005045064 | S.gene |
| CLSPN | CLSPN | ENSOKIG00005015558 | S.gene |
| CLSPN | CLSPN.1 | ENSOKIG00005048701 | S.gene |
| POLA1 | pola1 | ENSOKIG00005036207 | S.gene |
| CHAF1B | chaf1b.1 | ENSOKIG00005013623 | S.gene |
| CHAF1B | chaf1b | ENSOKIG00005003095 | S.gene |
| BRIP1 | brip1 | ENSOKIG00005015939 | S.gene |
| E2F8 | e2f8 | ENSOKIG00005005287 | S.gene |
| E2F8 | e2f8.1 | ENSOKIG00005018575 | S.gene |
| HMGB2 | hmgb2a | ENSOKIG00005013022 | G2M.gene |
| HMGB2 | hmgb2b | ENSOKIG00005037789 | G2M.gene |
| HMGB2 | hmgb2a.1 | ENSOKIG00005007373 | G2M.gene |
| CDK1 | cdk1.1 | ENSOKIG00005046533 | G2M.gene |
| CDK1 | cdk1 | ENSOKIG00005025794 | G2M.gene |
| NUSAP1 | nusap1 | ENSOKIG00005007844 | G2M.gene |
| NUSAP1 | nusap1.1 | ENSOKIG00005015733 | G2M.gene |
| UBE2C | ube2c | ENSOKIG00005000462 | G2M.gene |
| UBE2C | ube2c.1 | ENSOKIG00005000467 | G2M.gene |
| BIRC5 | birc5a | ENSOKIG00005003843 | G2M.gene |
| BIRC5 | birc5b | ENSOKIG00005039805 | G2M.gene |
| TPX2 | tpx2 | ENSOKIG00005017081 | G2M.gene |
| TOP2A | top2a | ENSOKIG00005014247 | G2M.gene |
| NDC80 | ndc80 | ENSOKIG00005001262 | G2M.gene |
| CKS2 | cks2.1 | ENSOKIG00005039691 | G2M.gene |
| CKS2 | CKS2 | ENSOKIG00005033377 | G2M.gene |
| CKS2 | cks2 | ENSOKIG00005022273 | G2M.gene |
| NUF2 | nuf2 | ENSOKIG00005000381 | G2M.gene |
| CKS1B | cks1b | ENSOKIG00005020717 | G2M.gene |
| CKS1B | CKS1B | ENSOKIG00005027388 | G2M.gene |
| MKI67 | mki67 | ENSOKIG00005013958 | G2M.gene |
| TMPO | tmpob.1 | ENSOKIG00005045692 | G2M.gene |
| TMPO | tmpoa | ENSOKIG00005012700 | G2M.gene |
| TMPO | tmpob | ENSOKIG00005016372 | G2M.gene |
| CENPF | cenpf | ENSOKIG00005011884 | G2M.gene |
| TACC3 | tacc3 | ENSOKIG00005001785 | G2M.gene |
| SMC4 | smc4 | ENSOKIG00005048554 | G2M.gene |
| CCNB2 | ccnb2.1 | ENSOKIG00005049632 | G2M.gene |
| CCNB2 | ccnb2 | ENSOKIG00005050238 | G2M.gene |
| CKAP2L | ckap2l | ENSOKIG00005002616 | G2M.gene |
| AURKB | AURKB | ENSOKIG00005021484 | G2M.gene |
| AURKB | aurkb | ENSOKIG00005050742 | G2M.gene |
| BUB1 | bub1 | ENSOKIG00005002111 | G2M.gene |
| KIF11 | kif11.1 | ENSOKIG00005046642 | G2M.gene |
| KIF11 | kif11 | ENSOKIG00005023299 | G2M.gene |
| ANP32E | ANP32E.2 | ENSOKIG00005000095 | G2M.gene |
| ANP32E | ANP32E | ENSOKIG00005013394 | G2M.gene |
| ANP32E | ANP32E.3 | ENSOKIG00005029241 | G2M.gene |
| ANP32E | ANP32E.1 | ENSOKIG00005034814 | G2M.gene |
| GTSE1 | gtse1 | ENSOKIG00005022873 | G2M.gene |
| KIF20B | kif20bb | ENSOKIG00005007330 | G2M.gene |
| KIF20B | kif20bb.1 | ENSOKIG00005022101 | G2M.gene |
| KIF20B | kif20ba | ENSOKIG00005041578 | G2M.gene |
| KIF20B | kif20ba.1 | ENSOKIG00005039441 | G2M.gene |
| CDCA3 | si:ch211-69g19.2 | ENSOKIG00005005809 | G2M.gene |
| CDC20 | cdc20 | ENSOKIG00005017478 | G2M.gene |
| TTK | ttk.1 | ENSOKIG00005000616 | G2M.gene |
| TTK | ttk.1 | ENSOKIG00005012015 | G2M.gene |
| CDC25C | ccdc25 | ENSOKIG00005044242 | G2M.gene |
| KIF2C | kif2c.1 | ENSOKIG00005038836 | G2M.gene |
| KIF2C | kif2c | ENSOKIG00005026504 | G2M.gene |
| RANGAP1 | RANGAP1 | ENSOKIG00005007651 | G2M.gene |
| RANGAP1 | rangap1a | ENSOKIG00005012910 | G2M.gene |
| RANGAP1 | rangap1b | ENSOKIG00005004716 | G2M.gene |
| NCAPD2 | ncapd2 | ENSOKIG00005019394 | G2M.gene |
| DLGAP5 | dlgap5 | ENSOKIG00005009969 | G2M.gene |
| CDCA2 | gene:ENSOKIG00005000746 | ENSOKIG00005000746 | G2M.gene |
| CDCA8 | cdca8 | ENSOKIG00005014820 | G2M.gene |
| CDCA8 | cdca8.1 | ENSOKIG00005048853 | G2M.gene |
| ECT2 | ect2.1 | ENSOKIG00005015498 | G2M.gene |
| ECT2 | ect2 | ENSOKIG00005012999 | G2M.gene |
| KIF23 | KIF23.1 | ENSOKIG00005024942 | G2M.gene |
| KIF23 | KIF23 | ENSOKIG00005038931 | G2M.gene |
| HMMR | hmmr | ENSOKIG00005028767 | G2M.gene |
| AURKA | aurka | ENSOKIG00005026781 | G2M.gene |
| ANLN | ANLN | ENSOKIG00005016220 | G2M.gene |
| ANLN | anln | ENSOKIG00005012931 | G2M.gene |
| LBR | lbr | ENSOKIG00005028443 | G2M.gene |
| LBR | lbr.1 | ENSOKIG00005049627 | G2M.gene |
| CKAP5 | ckap5.1 | ENSOKIG00005021670 | G2M.gene |
| CKAP5 | ckap5 | ENSOKIG00005006293 | G2M.gene |
| CTCF | CTCF | ENSOKIG00005020058 | G2M.gene |
| CTCF | CTCF.1 | ENSOKIG00005050634 | G2M.gene |
| CTCF | ctcf.1 | ENSOKIG00005001291 | G2M.gene |
| CTCF | ctcf | ENSOKIG00005031329 | G2M.gene |
| NEK2 | nek2 | ENSOKIG00005012940 | G2M.gene |
| NEK2 | nek2.1 | ENSOKIG00005025708 | G2M.gene |
| G2E3 | g2e3 | ENSOKIG00005017163 | G2M.gene |
| G2E3 | g2e3.1 | ENSOKIG00005013265 | G2M.gene |
| GAS2L3 | gas2l3 | ENSOKIG00005003841 | G2M.gene |
| CBX5 | cbx5.1 | ENSOKIG00005012086 | G2M.gene |
| CBX5 | cbx5 | ENSOKIG00005031647 | G2M.gene |
